# Low-intensity focused ultrasound to human amygdala reveals a causal role in ambiguous emotion processing and alters local and network-level activity

**DOI:** 10.1101/2025.08.14.670358

**Authors:** Johannes Algermissen, Miruna Rascu, Lilian A. Weber, Tim den Boer, Eleanor Martin, Bradley Treeby, Michael Gray, Robin O. Cleveland, Marco K. Wittmann, William T Clarke, Elsa Fouragnan, Matthew FS Rushworth, Miriam C Klein-Flügge

## Abstract

The amygdala is a core region changed in depression, a disorder characterized by compromised emotion, motivation and learning processes. However, lesion studies in humans examining amygdala function have largely focused on its role in processing fear. It currently remains unclear what causal role the human amygdala plays in more complex emotion, motivation and learning processes in daily life. This is because it has not been possible to reversibly modulate amygdala activity non-invasively in humans. Here, we used transcranial focused ultrasound stimulation (TUS) to bilaterally perturb neural activity in the basolateral amygdala (BLA). In separate sessions (n = 87), 29 healthy volunteers received offline-TUS to bilateral BLA, mid-insula or sham before playing a novel emotional learning task validated in an independent large cohort online (n = 210). 7T-resting-state and metabolite signals clearly demonstrated target engagement. BLA-TUS reduced the BLA’s connectivity fingerprint and decreased its excitation/inhibition balance, suggesting an inhibitory effect of our TUS protocol on BLA activity. In behaviour, BLA-TUS caused a stimulation volume-dependent increase in the tendency to approach neutral, emotionally ambiguous faces, treating them more similarly to happy faces, and a slowing of reaction times for those two emotion categories. These effects were functionally and regionally specific and suggest a causal role for the amygdala in resolving emotional ambiguity. Our results provide important insights for future studies into mood disorders where ambiguous situations might be harder to resolve, which could contribute to existing emotional and learning biases.

## Introduction

The human amygdala is often thought of as the brain’s centre for emotional processing(Adolphs et al., 1994; Murray, 2007; Phelps and LeDoux, 2005). Its connectional fingerprint positions it as a unique interface between limbic and cognitive circuits(Amaral et al., 1992). Importantly, it is also one of the key regions showing abnormal metabolism in depression(Drevets, 2003; Drevets et al., 1997; Murray et al., 2011; Nord et al., 2021), where functional impairments involve blunted emotional responding(Pechtel et al., 2013; Scholl and Klein-Flügge, 2018; Steele et al., 2007; Whitton et al., 2016) and changes in affective and learning biases(Admon and Pizzagalli, 2015; Browning et al., 2019; Eshel and Roiser, 2010; Harmer et al., 2009a; Korn et al., 2013; Nord et al., 2018). However, whether these changes in cognitive and emotional processes causally relate to changes in amygdala function or might be mediated by other circuits affected in depression remains unclear. This is because until recently, it has not been possible to causally and selectively manipulate activity in deep brain regions non-invasively in human participants to study their precise functional contribution.

Causal investigation of amygdala function in humans has until recently been limited to patient populations with implants or lesions in this region. In line with a large body of animal work focusing on the role of the amygdala in fear conditioning(Janak and Tye, 2015; Krabbe et al., 2018; LeDoux, 2000), studies in human patients with amygdala lesions have predominantly studied impairments in the fear domain, showing decreased fear responses and a reduced ability to recognize fearful facial emotions, with strongest effects following bilateral lesions(Adolphs, 2013; Adolphs et al., 1999, 1994; Feinstein et al., 2011). However, intriguingly, human lesion studies have not highlighted a role for the amygdala in the core processes affected in depression.

By contrast, evidence from animal work using causal manipulations of amygdala circuits, via lesions or optogenetic approaches, emphasizes the importance of the amygdala in several other functional domains. For example, in the context of learning, lesions to the amygdala and connected circuits have highlighted its importance in reinforcement learning, value updating, sensitivity to negative feedback and non-contingent credit assignment(Costa et al., 2016; Izquierdo and Murray, 2007; Morrison and Salzman, 2010; Murray, 2007; Rudebeck et al., 2013; Taswell et al., 2021; Wassum and Izquierdo, 2015). Similarly, in the emotional domain, lesions to amygdala impair emotion recognition, emotion sensitivity and especially processing of ambiguous sensory inputs(Bernardi and Salzman, 2019; Dal Monte et al., 2015; Hadj-Bouziane et al., 2012; Izquierdo et al., 2005; Wang et al., 2017).

Thus, there is an urgent need to study emotion and learning processes using causal approaches in humans to help bridge the gap between causal studies in animals and the functional impairments experienced daily by patients with mood disorders which may, in part, rely on changes in amygdala circuits. Low-intensity transcranial ultrasound stimulation (TUS) offers the opportunity to non-invasively and temporarily perturb neural activity in deep brain regions of healthy human volunteers with millimetre precision. TUS applied offline has repeatedly succeeded in altering neural activity and behavioural read-outs in macaques, including in deep brain regions(Bongioanni et al., 2021; Deffieux et al., 2013; Folloni et al., 2019; Fouragnan et al., 2019; Khalighinejad et al., 2020; Verhagen et al., 2019). More recently, TUS has been shown to have similar effects in humans(In et al., 2024; Nakajima et al., 2022; Yaakub et al., 2024, 2023) and guidelines and standards have been developed to guide its safe use in humans(Aubry et al., 2023; Martin et al., 2024; Murphy et al., 2025b). Compared to other non-invasive brain stimulation techniques, TUS allows unprecedented depth and focality of stimulation, and, unlike DBS, it is non-invasive.

The aim of the current study was to test in healthy volunteers how the basolateral amygdala (BLA) causally contributes to emotion and learning processes, especially emotional biases and learning from probabilistic feedback. Studying the causal role of the amygdala in healthy populations provides an opportunity to understand its role in the absence of compensatory processes or long-term plastic changes present in patients with lesions. We developed and validated (online, n = 210) a novel emotional learning task and examined the functional and spatial specificity of offline TUS effects. We also recorded MR measures of functional resting-state connectivity and metabolite concentration for proof of TUS target engagement. We find that TUS to the BLA changed its connectivity to other regions as well as its E/I balance and altered behaviour, particularly for emotionally ambiguous stimuli. These effects only occurred under BLA, but not active control stimulation (of the mid-insula). Effects were functionally specific, sparing learning from feedback. Overall, our findings suggest a role for BLA in detecting and resolving emotional ambiguity in perceptual processing.

## Results

### Approach-avoidance learning task requires emotion recognition and evokes automatic emotional biases

To study emotional and learning processes relevant to daily life, we developed a task with two key features. First, participants needed to recognize an emotional expression and react to it using an approach or avoid response. Based on prior work relating amygdala function to emotional biases known to be altered in mood disorders(Bertsch et al., 2018; Bramson et al., 2023; Heuer et al., 2007; Radke et al., 2013), participants had to make emotion-congruent (e.g., approach happy), emotion-incongruent (e.g. avoid happy), and emotion-neutral (e.g. approach neutral) responses in separate mini-blocks of trials (**Fig 1a**)(Volman et al., 2011). Second, inspired by work pointing to an important role for the amygdala in learning(Costa et al., 2016; Izquierdo and Murray, 2007; Morrison and Salzman, 2010; Rudebeck et al., 2013; Taswell et al., 2021; Wassum and Izquierdo, 2015), we incorporated the need to learn the correct response from trial-and-error using probabilistic feedback, deciding whether negative feedback warrants a change in behaviour or not. Thus, instead of directly instructing a response, once a facial emotion was shown on the screen, the correct emotion-response mapping of each mini-block had to be learned by trial-and-error based on feedback, akin to reward-learning tasks(Behrens et al., 2008, 2007). A mini-block contained between four and seven consecutive trials showing different faces. Within mini-blocks, the displayed emotion of the different faces remained constant – either happy, neutral or angry. On each trial, participants chose to approach or avoid the stimulus (left or right button press). This would make the face grow or shrink, respectively. Participants were given probabilistic feedback for their response (80% valid). The correct response stayed constant within a given mini-block but could change at mini-block transitions. The six mini-blocks conditions (3 emotions x 2 responses) were presented in pseudo-random order. Transitions between mini-blocks were not instructed but signalled by a change in facial emotion (**Fig 1a**). Thus, participants faced two challenges: they needed to infer the correct response of the current mini-block from probabilistic feedback, while monitoring the facial emotions to infer the start of the next mini-block.

**Figure 1.**
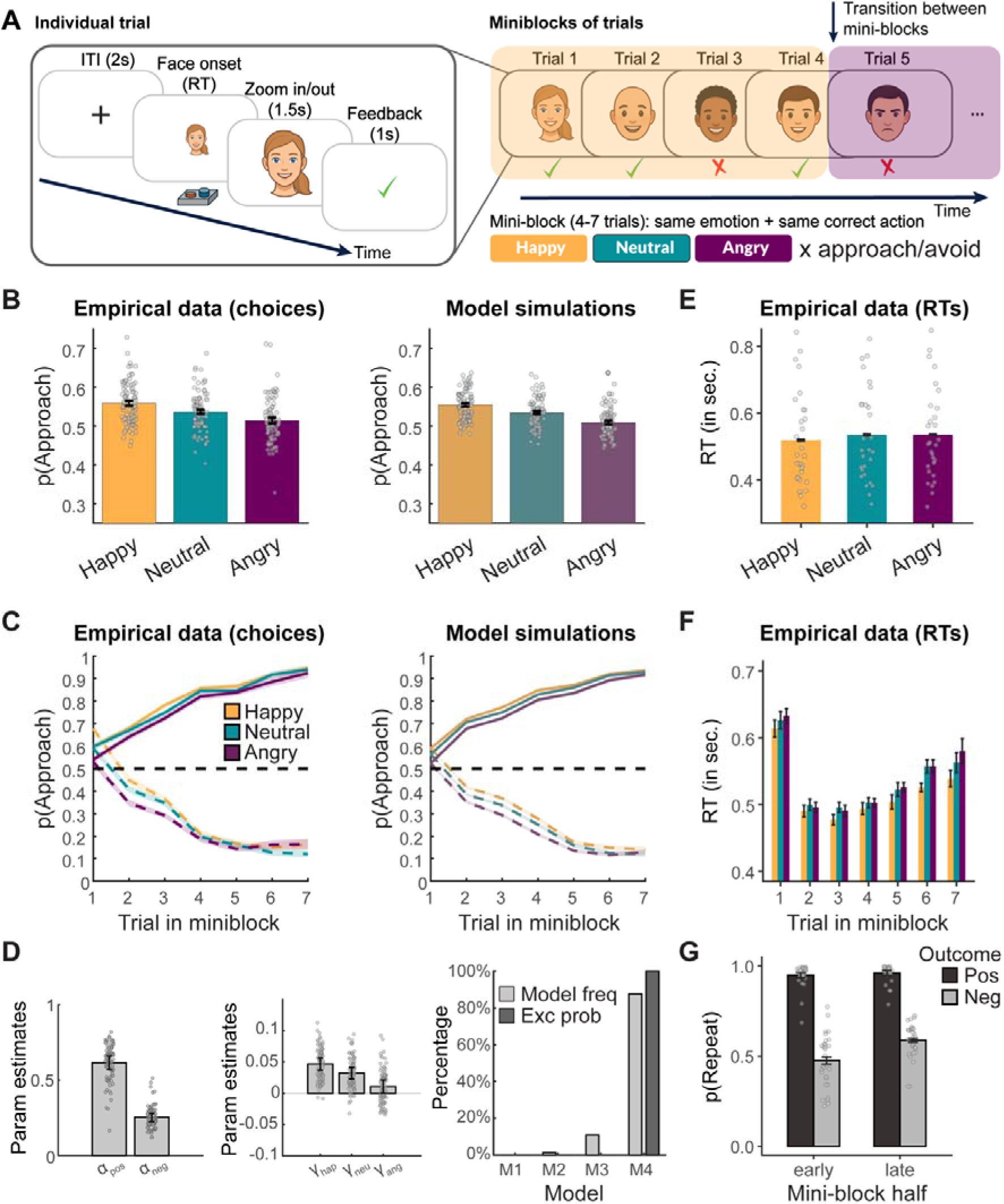
Learning task and RL modelling reveal emotional bias. **A**, *Left:* In each trial, participants (n = 29, Experiment 1; n = 87 sessions) saw a face showing one of three possible emotions – happy, angry or neutral. They pressed one of two buttons (left or right) to approach or avoid the face upon which the face grew or shrank, respectively, before receiving feedback on whether their response was accurate (probabilistic, 80% valid). *Right:* Faces were organised into mini-blocks of 4–7 consecutive trials showing different faces with the same emotion (e.g. here four happy faces in a row). The correct response of each mini-block had to be learned by trial-and-error based on feedback and stayed constant within a given mini-block. The six mini-blocks conditions (3 emotions x 2 responses) were presented in pseudo-random order. Transitions between mini-blocks were not instructed, but signalled by a change in facial emotion. Thus, participants needed to learn the correct response of the current mini-block, while monitoring the facial emotion to infer the start of the next mini-block. **B,** *Left:* Participants showed an emotional bias in their responding. Due to the fully balanced design, the expected p(approach) is 50% for each emotion. However, in our data, participants were more likely to approach happy over neutral and neutral over angry faces. *Right:* A reinforcement learning (RL) model with separate learning rates for positive and negative feedback and separate approach biases for each emotion could mimic participants’ behaviour. **C,** *Left:* Over trials in a mini-block, participants correctly learned when to approach (solid) or avoid (dashed) the faces (happy = yellow; neutral = turquoise; angry = purple). *Right:* Again, the model was able to capture this behaviour. **D**, Parameter estimates from the winning computational model (M4) showed faster learning from positive than negative feedback (left) and decreasing approach biases from happy to neutral and neutral to angry faces (centre). Model comparison shows that M4 best captures behaviour (right). **E,** Reaction times (RTs) mirrored the emotional bias seen in choices in B, with faster RTs in happy than neutral and angry trials. **F,** Reaction times were fastest on trials 2–4 and slowed down with the likelihood of a new mini-block starting (likelihood was highest on trial 1, lowest on trials 2–4 and increasing in likelihood from trial 5 to 7). **G,** Participants were more likely to repeat their response following positive compared to negative feedback and late compared to early in the mini-block, showing that feedback was taken into account appropriately. The behaviour of n = 210 participants collected online as part of Experiment 2 confirmed these results in the absence of any TUS (**Fig S1**). Error bars denote standard error of the mean (SEM) computed using Cousineau-Morey correction(Morey, 2008), which removes between-subject variation (i.e. in the per-subject intercept) irrelevant for evaluating within-subject effects.

Two separate samples of participants performed the task either in the laboratory (n = 29 MRI+TUS) or online (n = 210). Online participants completed one session of the task (see Methods). MRI+TUS participants completed four in-person testing sessions on separate days, spread over a period of 4–6 weeks (**Fig 2a**, Methods). This included an initial MRI scan for ultrasound planning, and three TUS+MRI sessions (n = 87 in total). These sessions comprised the behavioural task and a TUS+MRI procedure that was identical in each session apart from the ultrasound stimulation target, which we return to below (basolateral amygdala/BLA: area of interest, mid-insula/mIns: active control, or sham; order counterbalanced; see Methods).

**Figure 2.**
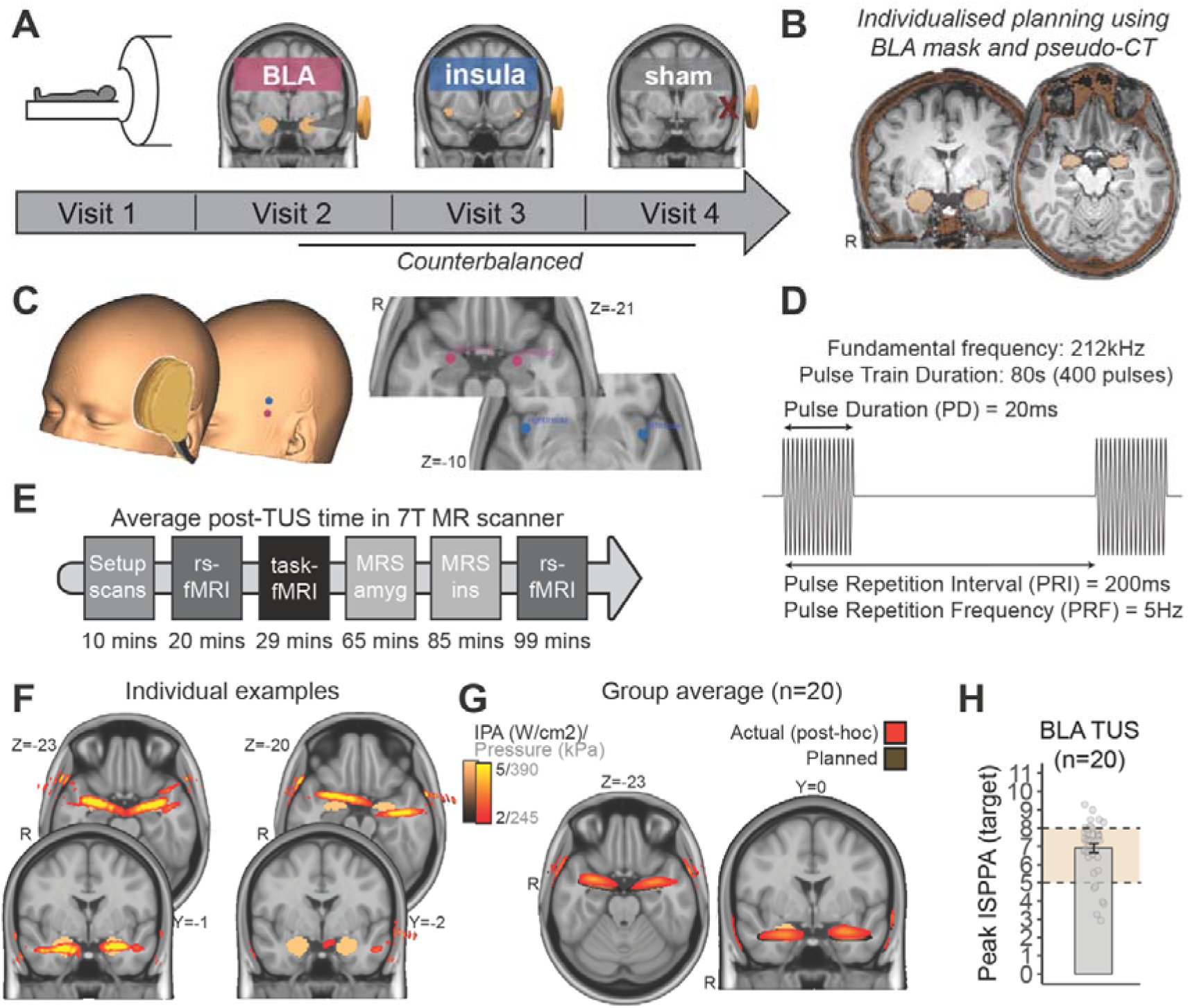
Transcranial focused ultrasound targeting. **A**, Overview of study timeline. Following an initial MR visit to allow for individualized planning of the ultrasound stimulation, participants attended three TUS-MRI sessions that were identical in their procedure apart from the TUS stimulation site (basolateral amygdala (BLA), mid-insula (mIns), or sham; counterbalanced order). **B,** Individual’s structural scan (grey), a pseudo-CT image derived from their PETRA scan as well as a BLA mask projected into participant space (both overlaid in copper) were used to run individualized ultrasound stimulations *ex ante,* between visit 1 and the subsequent TUS visits, to decide on the optimal stimulation trajectory through the temporal window. **C,** Schematic showing approximate placement of the transducer for the left hemisphere (left) and the average skin (middle) and target (right) locations used in the experiment (in MNI space; for full coordinates, see **Tables S7+S8**). **D,** In all TUS visits, a 5 Hz patterned protocol of 80 seconds duration (PD: 20 ms, PRI: 200 ms, 10% duty cycle, 5 Hz PRF, 400 pulses; **Table S2**) was applied to the left hemisphere (BLA, mIns or sham) followed by the right hemisphere (BLA, mIns, or sham) outside the MR scanner. **E,** Following TUS, participants immediately entered a 7T-MRI scanner and completed several MR scans in the order shown. Two blocks of the behavioural task were played during task-fMRI and two during the MRS voxel acquisition. **F,** Following each TUS visit, simulations were repeated *post-hoc* using the actual location of the transducer recorded during the session via neuronavigation. Shown are two example participants: one of the worst who was excluded because the BLA was missed, and one of the best where targeting was accurate. A total of 20 participants out of 29 were retained for analyses. **G,** Comparison of planned *ex ante* (brown) and actual (orange-red) post-hoc simulations averaged across all included BLA-TUS participants (n = 20). Overall, there was good agreement between planned *ex ante* and *post-hoc* simulations, with slightly better accuracy in the right hemisphere. Colour map shows references to Pulse Average Intensity (I_PA_) or Pressure values. **H,** The peak achieved in situ I_SPPA_ of the n = 20 included BLA-TUS participants as inferred from *post-hoc* simulations (planned: yellow range between 5–8 W/cm^2^; achieved: mean of 6.9 W/cm^2^, range 2.9–9.3 W/cm^2^).

First, we characterised participants’ overall choice behaviour, independent of TUS, to provide evidence that our task elicited emotional biases (**Fig 1b**) and probed trial-by-trial learning (**Fig 1c**), the two processes we hypothesized might be changed following TUS to the BLA. A mixed-effects logistic regression predicting approach/avoid responses as a function of the required response, emotion of the displayed face, and mini-block half (trials 1–4 vs. 5– 7) showed across all TUS sessions (n = 87) that participants exhibited an emotional bias in the task (**Fig 1b-left;** main effect of emotion; χ^2^(2) = 23.863, *p* < .001), with overall more approach responses to happy than neutral faces (*b* = 0.059, 95%-CI [0.018 0.010], χ^2^(1) = 7.792, *p* = .005) and more approach responses to neutral than angry faces (*b* = 0.073, 95%-CI [0.033 0.112], χ^2^(1) = 13.074, *p* < .001; **Fig 1c**). Furthermore, participants learned the correct response of each mini-block (**Fig 1c-left and Fig S1b;** main effect of required response, *b* = 1.169, 95%-CI [1.074, 1.263], χ^2^(1) = 586.616, *p* < .001; mean/median accuracy per participant 0.72, range 0.60–0.77), with better performance in the 2^nd^ compared to the 1^st^ half of mini-blocks (2-way interaction required response x mini-block half; *b* = 0.706, 95%-CI [0.627 0.785], χ^2^(1) = 306.010, *p* < .001; for equivalent statistics for the n = 210 online participants, see Supplementary Information and **Fig S1**).

A simple reinforcement learning (RL) model(Barto et al., 1981; Kang et al., 2024; Rescorla and Wagner, 1972) fitted on all TUS sessions was able to reproduce the same two key behaviours, the emotional bias and learning curve (**Fig 1b/c-right;** replicated for online data in **Fig S1**). The winning model (M4) incorporated separate learning rates for positive and negative outcomes, an inverse temperature parameter and three approach biases, one per emotion. This model performed better than simpler models M1–M3 with just one learning rate and no or only one approach bias (see Methods; Bayesian model selection: M4: model frequency: 88%; protected exceedance probability: 100%; **Fig 1d**). Indeed, M4 was the only model that could reproduce the key features observed in participants’ behaviour (**Fig 1b-c** **right**; **Fig S2**). For model parameters, see Fig 1d (associated statistics on model parameters are reported in the Supplementary Information and **Tables S22-S25**).

Second, we analysed participants’ overall reaction times (RTs) to see if they similarly reflected emotional biases and task learning. To test for an emotional bias in RTs, we regressed RTs onto the emotion of the displayed face (happy/neutral/angry), feedback on the previous trials (positive/negative), and mini-block half (first/second). Indeed, participants displayed an emotional bias also in their RTs (main effect of emotion; χ^2^(2) = 12.841, *p* < .001), with faster responses to happy faces than to neutral faces (*b* = -0.019, 95% CI [-0.032 - 0.006], χ^2^(1) = 8.038, *p* = .005) and angry faces (*b* = -0.021, 95% CI [-0.033 -0.009], χ^2^(1) = 11.542, *p* < .001), but no difference between neutral and angry faces (*b* = -0.001, 95% CI [-0.014 0.012], χ^2^(1) = 0.021, *p* = .884; **Fig 1e**). Looking at the RT evolution across trials in a mini-block (**Fig 1f**), we found RTs were fastest on trial #2 of a block, slowed down towards the end of a mini-block, and were slowest on trial #1 of a new mini-block (**Fig 1e**). Thus, RTs slowed down proportionally to the probability of a new mini-block starting, which was lowest on trials #2–4 and then strongly increased up until trial #1 of the next mini-block. This shows that participants learned the task structure and slowed down responses as the onset of a new mini-block became more likely(Kang et al., 2024) (for full model statistics for both online and in-person cohorts, see Supplementary Results and Tables, **Fig S1**).

Third, we found that participants based their choice to repeat or switch their response on whether they had received positive and negative feedback, providing further evidence for their ability to learn from feedback. We regressed choices to repeat/switch responses onto the feedback they had received on the previous trial (positive/negative) and the mini-block half (first/second). Participants showed an overall tendency to repeat responses (intercept: *b* = 1.870, 95% CI [1.635 2.105], χ^2^(1) = 244.115, *p* < .001), repeated their responses more often after positive than negative feedback (main effect of feedback: *b* = 1.743, 95% CI [1.554 1.932], χ^2^(1) = 328.319, *p* < .001) and more in the second than in the first half of mini-blocks (main effect of block-half: *b* = 0.217, 95% CI [0.125 0.308], χ^2^(1) = 21.373, *p* < .001; **Fig 1g**; for the full model for both cohorts, see Supplementary Tables).

In summary, the overall behaviour recorded in our task showed evidence for emotional biases and the ability to learn from feedback. We hypothesized that BLA-TUS might modulate one or both of these processes.

### Targeting the human BLA with ultrasound

Having characterized participants’ overall behaviour in the task, we next turned to the effects of stimulating the BLA using ultrasound. Before comparing behaviour in BLA-TUS sessions with those obtained in no-TUS sham sessions, we inspected whether we were successful in targeting the BLA. We ran acoustic and thermal simulations prior to all TUS sessions using the individual’s structural brain and skull scan acquired during their first visit (**Fig 2b** and Methods). This was important to account for reflection, refraction, and absorption of the ultrasound beam when passing through the skull and aimed to reduce variability in the achieved stimulation across participants. It was also critical for establishing that all ultrasound sessions adhered to ITRUSST safety guidelines (mechanical index for transcranial applications (MI_tc_) <u>≤</u> 1.9 and thermal dose ≤ 0.25 cumulative equivalent minutes at 43° (CEM43): Methods and(Aubry et al., 2024); **Fig S3**). We iteratively refined the transducer positioning to achieve a maximum target in-situ intensity (Spatial Peak Pulse Average Intensity: I_SPPA_) of 5–8 W/cm^2^ in each participant while minimising energy delivery to off-target regions and ensuring mechanical and thermal safety. Our planned targeting was accurate as reflected in the mean Pulse Average Intensity (I_PA_) map shown in brown in **Fig 2g**.

Once acoustic simulations were completed, participants were invited to visits 2–4, where TUS was delivered to the left and right hemispheres (BLA, mIns or sham, counterbalanced; **Fig 2a**) for a total of 80 seconds, following a 5 Hz pulse repetition frequency (PRF) protocol (pulse width: 20 ms, pulse repetition interval: 200 ms; duty cycle: 10%(Yaakub et al., 2023; Zeng et al., 2022); **Fig 2d**; **Table S2**). During TUS delivery, the participant was at rest, and we used neuronavigation to track the position of the transducer and participant for the duration of the sonication (see Methods; mean skin coordinate: **Fig2c**). This allowed us to verify whether we had followed our planned trajectory and precisely targeted the BLA (or mIns in active control sessions). We used the tracked transducer location to run acoustic *post-hoc* simulations (see Methods). These simulations showed that, on average, our targeting was close to the planned trajectory, but slightly better in the right compared to the left hemisphere (red-orange overlay in **Fig 2g**; for visualisation of planned coordinates only, see **Fig S3f-g**). To quantify our TUS targeting precision on a participant-by-participant basis, we used a BLA mask from our previous work(Klein-Flügge et al., 2022) and computed the BLA volume in mm^3^ stimulated in each hemisphere with an in situ I_PA_ of at least 2 W/cm^2^ (see Methods; other choices of threshold led to highly correlated estimates).

Prior work from lesion studies in both animal models and humans suggests behavioural effects are particularly prominent following bilateral amygdala lesions(Costa et al., 2016; De Martino et al., 2010; Ghods-Sharifi et al., 2009; Hampton et al., 2007; Winstanley et al., 2004). Thus, we quantified the extent to which both left and right BLA were stimulated with TUS in each participant. Values of bilateral stimulation were expressed as the BLA volume targeted with at least 2 W/cm^2^ I_PA_ in the less stimulated hemisphere (while somewhat arbitrary, other choices of threshold lead to highly correlated covariates, e.g. correlation of covariates thresholded at 2 W/cm^2^ and 3 W/cm^2^: *r* = 0.981; thresholded at 2 W/cm^2^ and 1 W/cm^2^: *r* = 0.990). In other words, if the volume of stimulated tissue differed across hemispheres, this value quantifies the lower bound of the volume successfully stimulated in either hemisphere. Two representative participants with good and bad bilateral amygdala targeting, respectively, are shown in **Fig 2f**. Overall, nine participants received less than 20% bilateral BLA stimulation (corresponding to < 340 mm^3^ in stimulated bilateral volume > 2 W/cm^2^ I_PA_) and were excluded from further analyses related to BLA-TUS. The remaining n = 20 participants were considered across statistical analyses related to BLA-TUS effects (minimum volume stimulated > 2 W/cm^2^ per participant in mm^3^ in these n = 20 participants: *M* = 859, *SD* = 305, range 354–1,369). In these participants, on average, the maximum I_PA_ achieved in the BLA was 6.9 W/cm^2^ (*SD* = 1.5, range 2.9– 9.3) (**Fig 2h**).

### BLA-TUS changes resting-state connectivity

If TUS achieved effective neuromodulation in the BLA, we expected this should evoke differences in the amygdala’s resting-state connectivity fingerprint in a way analogous to previous non-human and human TUS studies targeting other regions(Folloni et al., 2019; Khalighinejad et al., 2020; Verhagen et al., 2019; Yaakub et al., 2023). Thus, as a direct proof of target engagement, in our final sample of n = 20 participants, we inspected the resting-state BOLD connectivity fingerprint of the BLA with its closely connected regions. Following the bilateral ultrasound stimulation delivered outside the scanner, participants immediately transitioned to a high-field 7T MRI scanner. Two resting-state runs of seven minutes each were acquired at the start and end of each scan session (**Fig 2e**). Across both runs, we found that BLA connectivity to monosynaptically connected regions summarized in a connectivity fingerprint(Verhagen et al., 2019; Yaakub et al., 2023) was reduced following BLA compared to sham stimulation sessions (*t*(19) = -3.552, *p* = .002; **Fig 3d-e**; see Methods for details on ROI selection; for comparisons with the active control TUS in mid-insula, see **Fig 6** and **S4**). Regions that generally showed a positive resting-state connectivity with the BLA (computed as the average across sham and BLA sessions) showed a reduced connectivity profile with the BLA following the sonication (**Fig 3b-e**; **Fig S4**). In a mixed-effects linear regression with the predictors TUS condition (amygdala vs. sham) and run (run1/run2), we found only a main effect of TUS condition (*b* = 0.050, 95% CI [0.032 0.078], *t*(76) = 3.640, *p* < .001; no effect of run and no interaction), suggesting that the TUS effect was comparable between the first resting-state run, which followed relatively soon after TUS (average time of first resting-state run: 19 minutes post-TUS), and the second run, which was up to 2h 12 min after TUS (average time of second resting-state run: 99 minutes post-TUS). Taken together, despite some heterogeneity in our targeting, post-hoc simulations and resting-state connectivity fingerprints consistently indicated that TUS successfully targeted the BLA in n = 20 participants, and that TUS-induced neural effects lasted well over an hour.

**Figure 3.**
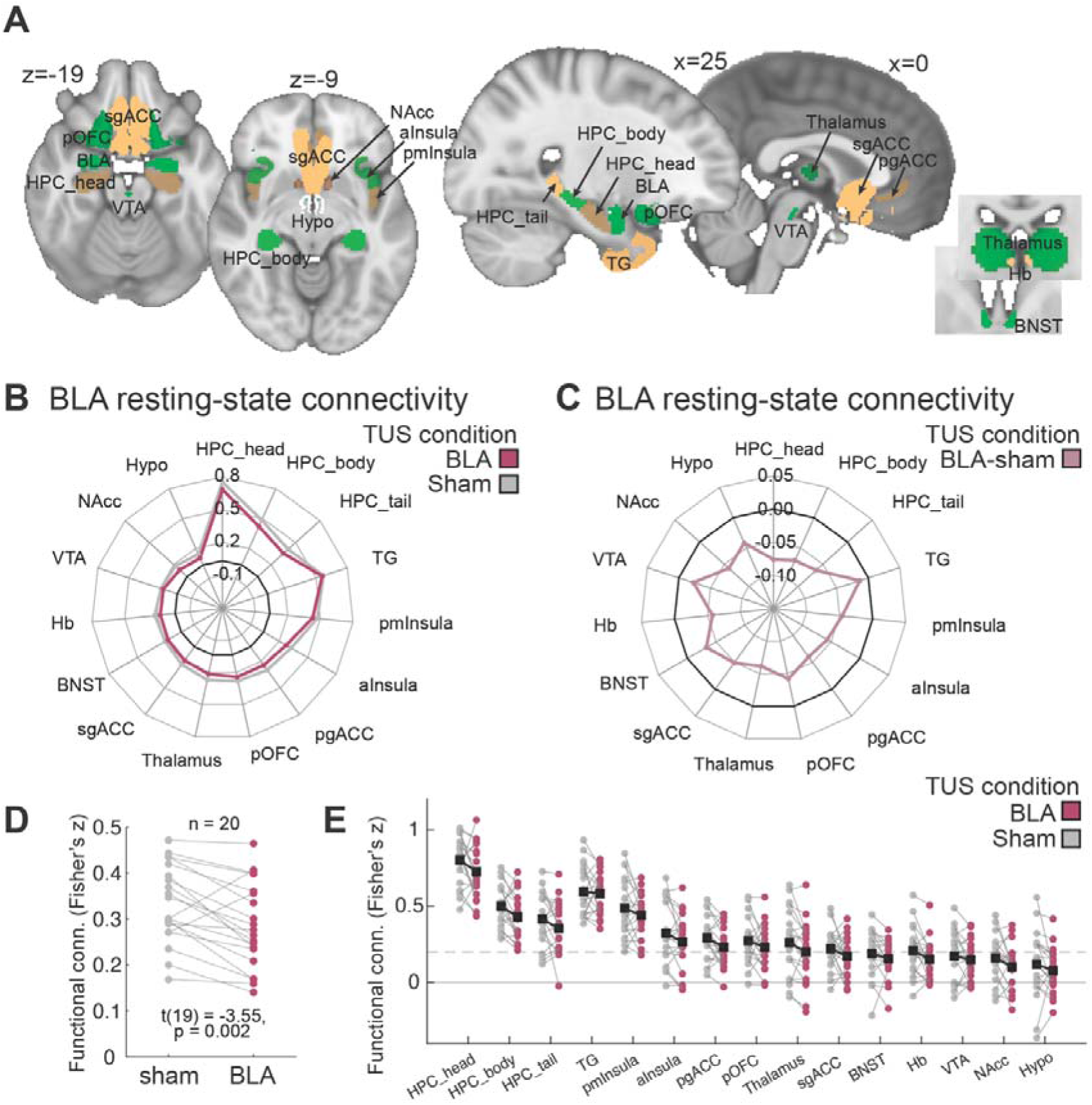
TUS target engagement probed using resting-state fMRI. **A**, ROIs used for extracting connectivity with BLA in panels b-e. **B,** Fingerprint of BLA functional connectivity with monosynaptically connected regions of interest (ROIs) following BLA (pink) and sham (grey) sonication shows a consistent reduction in connectivity following BLA-TUS. **C,** Direct comparison of sonication conditions (BLA stimulation minus sham: light pink) emphasizes the consistently reduced fingerprint (the 0 line indicating no difference is shown in black), suggesting BLA activity is driven less by its directly connected nodes following TUS to this region. **D,** Average connectivity across all connections in the fingerprint shown in A shows reductions are consistent across participants. **E**, Mean connectivity shown in A is shown including individual datapoints.

### BLA-TUS tends to decrease E/I balance measured with MRS

Resting-state data showed evidence for connectivity changes between the amygdala and its connected network. To additionally gain insights into local BLA processing and proxies of excitation/inhibition, we also measured metabolite concentrations using magnetic resonance spectroscopy (MRS). We acquired MRS data in two voxels of size 20 x 14 x 20 mm corresponding to our target locations, one placed in the BLA, and one in mIns. We note that studies measuring reliable MRS signals in the amygdala are rare(Nacewicz et al., 2012; Steinegger et al., 2024). The amygdala is difficult to image with MRS due to its large distance from the MR head coils and surrounding CSF and blood vessels, which can induce motion artifacts and susceptibility changes and limit the positioning of the MRS voxel. To compensate for low signal-to-noise ratio, we used a MEGA-edited semi-LASER sequence suited for measuring GABA concentration, measured at a high field strength of 7T, and applied stringent exclusion criteria. Among our n = 20 participants with successful BLA stimulation, n = 3 showed insufficient overlap between the *post-hoc* simulated ultrasound pressure beam and the MRS voxel placement (less than 200 mm^3^/5% overlap between ultrasound beam thresholded at > 2 W/cm^2^ and MRS voxel; **Fig 4a**). Of the remaining n = 17, n = 7 did not have sufficient MRS data quality in the BLA MRS voxel in at least one of the two sessions (BLA or sham, see Methods). However, we were able to include one additional participant who was initially excluded due to poor task performance but showed adequate TUS/MRS overlap and MRS quality. Thus, we included data from n = 11 participants and n = 22 sessions (BLA + sham) with sufficient data quality in this analysis. Their MRS voxel position and average MRS spectrum and fit is shown in **Fig 4b** and **Fig 4c**.

**Figure 4.**
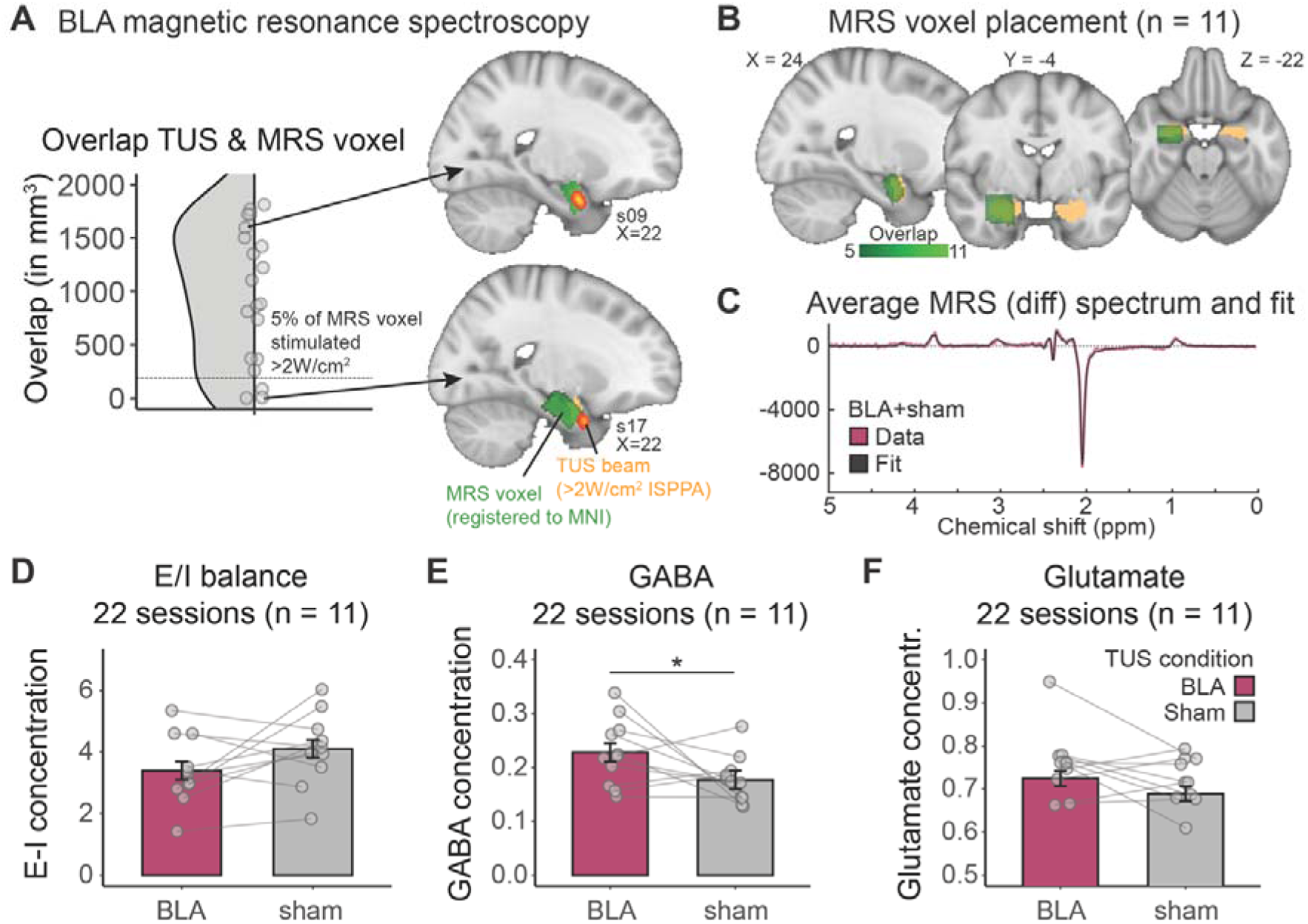
TUS metabolite changes probed using Magnetic Resonance Spectroscopy. **A**, Placing an MRS voxel such that it contains BLA is difficult given the amygdala’s proximity to major vessels and CSF boundaries. Inspection of overlap between MRS voxel placement and TUS post-hoc simulations (>2 W/cm^2^ I_PA_) revealed n = 17 participants with sufficient overlap (>200 mm^3^ = dashed line). Two example participants with sufficient (top) and insufficient (bottom) overlap are shown. **B,** Overlap in the MRS voxel placement across participants: shown is the sum over the n = 11 remaining participants following exclusion of datasets with insufficient MRS data quality (see Methods); MRS voxels were registered to MNI space, binarized, and summed over participants to show overlap. **C,** Average MRS difference spectrum for all n = 22 sessions (BLA-TUS and sham in each of n = 11 participants; for separate fits for BLA vs sham sessions, the fit of the off spectrum, and MRS quality metrics, see **Fig S5**). **D,** The E/I balance was reduced in 7 out of 11 people (*p* = .087). **E,** This was due to a significant increase in GABA in 8 out of 11 participants (*p* = .025). **F,** BLA-TUS did not lead to any changes in glutamate. Non-parametric tests yielded similar results (Wilcoxon signed-rank test: E/I: V = 17, p = 0.175; GABA: V = 54, p = 0.067; Glutamate: V = 46, p = 0.278).

In these datasets, we first inspected the ratio of glutamate and γ-aminobutyricacid (GABA) concentrations, often referred to as excitation/inhibition or E/I balance, as an index of neural excitability(Scholl et al., 2017; Stagg et al., 2011) before separately considering GABA and glutamate changes (**Fig 4d-f**). Compared to sham sessions, BLA-TUS decreased the E/I balance in 7 out of 11 participants **(Fig 4d)**, which was marginally significant at the group-level (χ^2^(1) = 2.924, *p* = .087, *d* = -0.516). Importantly, when broken down into the individual metabolites, we observed a significant increase in GABA, χ^2^(1) = 5.057, *p* = .025, *d* = 0.620 (increase in 8 out of 11 participants; **Fig 4e**), but no change in glutamate, χ^2^(1) = 2.054, *p* = .152, *d* = 0.432 (**Fig 4f**). Qualitatively similar results were obtained using non-parametric tests and when including gender, age, the time of day at data acquisition, and simulated pressure beam/MRS voxel overlap as covariates of no interest (**Table S26**; further MRS quality checks and separate BLA vs sham fits are shown in **Fig S5**). These results suggest that BLA-TUS might have had an inhibitory (rather than excitatory) effect on neural activity, which aligns with the BLA-TUS effect on resting-state connectivity.

### BLA-TUS changes approach bias to neutral facial expressions

Returning to the behavioural signatures recorded in our task, we next examined whether BLA-TUS had induced any changes in emotional approach/avoid biases or the ability to learn from feedback. As mentioned above, based on work in patients with brain lesions, we reasoned that behavioural effects of TUS might scale with the bilaterality of the stimulation(Li et al., 2016). Our continuous quantification of the bilaterality (“Bilaterality of BLA stimulation”) ranged between 354–1,369 mm^3^ (volume >2 W/cm^2^ I_PA_ of the less stimulated hemisphere) and was therefore included as a covariate in behavioural analyses (see **Fig S3c**).

We first tested our first hypothesis that BLA-TUS might alter emotional biases and thus participants’ tendency to approach/avoid faces based on the emotions they displayed. A mixed-effects logistic regression predicting participants’ responses (approach/avoid) using the stimulation condition (BLA/sham), the emotion of the face stimulus (happy/neutral/angry), and the bilaterality covariate showed a significant 3-way interaction between stimulation condition, emotion, and the bilaterality covariate (χ^2^(2) = 10.079, *p* = .006; **Fig 5a**), suggesting that BLA-TUS changed participants’ emotional bias in a way that scaled with the achieved bilaterality of the stimulation (for other main effects and interactions, see Supplementary Information). Post-hoc tests examining this result further revealed that TUS selectively increased the propensity for approaching neutral face stimuli in a way that scaled with the bilaterality of BLA stimulation (neutral stimuli only: 2-way interaction stimulation condition x covariate: *b* = 0.074, 95%-CI [0.023 0.124], χ^2^(1) = 8.145, *p* = .004; correlation TUS effect p(approach|TUS) – p(approach|sham) with covariate: *r*(18) = 0.669, *p* = .001; **Fig 5b**). There was no such change for happy face stimuli (*b* = -0.022, 95%-CI [-0.074 0.030], χ^2^(1) = 0.701, *p* = .403; correlation TUS effect with covariate: *r*(18) = - 0.178, *p* = .452) or angry face stimuli (*b* = 0.029, 95%-CI [-0.023 0.082], χ^2^(1) = 1.111, *p* = .292; correlation TUS effect with bilaterality covariate: *r*(18) = -0.220, *p* = .351; **Fig S6a,c**). Furthermore, the identified relationship of increased approaching of neutral faces following BLA-TUS was present independent of whether participants were asked to approach or avoid neutral faces (**Fig S6e,f**).

**Figure 5.**
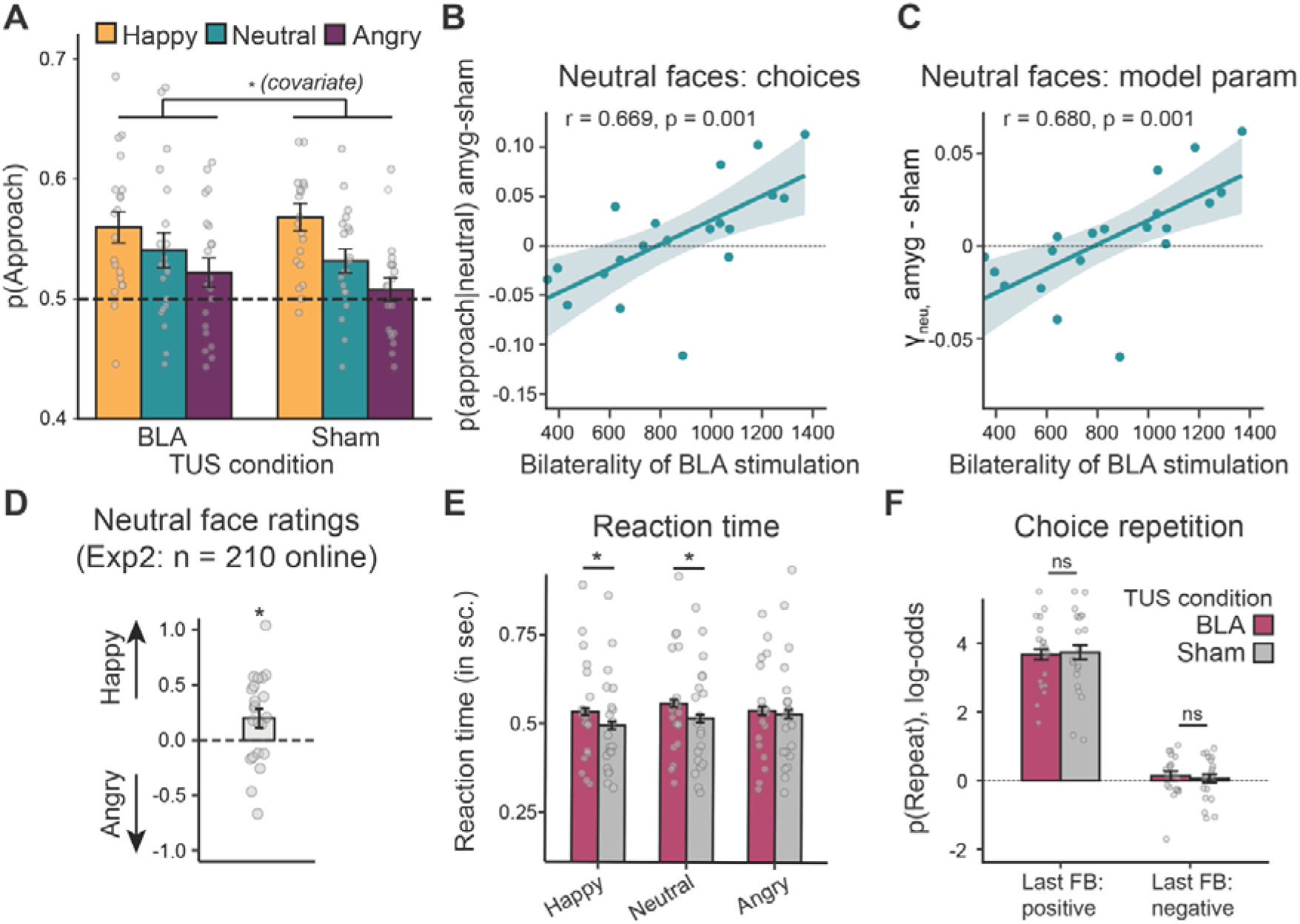
TUS to basolateral amygdala changes approach biases and RTs to ambiguous facial expressions. **A**, The proportion of trials when participants approached happy (yellow), neutral (turquoise), or angry (purple) faces was changed following BLA-TUS. **B,** The bilaterality of BLA stimulation (volume > 2 W/cm^2^ in the less stimulated hemisphere) predicted the increase in approaching neutral facial stimuli after BLA-TUS. The more volume of the BLA was stimulated bilaterally, the more likely participants became to approach neutral faces. There was no relationship between BLA stimulation and participant’s likelihood to approach happy or angry faces (**Fig S6a,c**). **C,** This finding was captured by the model-derived parameter reflecting the approach bias to neutral faces (but not happy or angry faces, see **Fig S6b,d**). **D,** The 22 neutral face identities used in our task were rated by an independent cohort of n = 210 online participants (Experiment 2) which showed that these neutral face stimuli were on average perceived as more happy than angry, suggesting neutral and happy faces were closer in perceptual similarity and could more easily be confused in our task. **E,** Reaction times (RTs) were slower following BLA-TUS compared to sham for happy and neutral, but not angry face stimuli, and thus the two perceptually closer emotion categories. **F,** The probability of repeating an action following positive or negative feedback was not changed following BLA-TUS (shown in log-odds space for improved visualization), suggesting basic learning from feedback was spared.

In line with this finding, in our winning computational reinforcement learning model, the difference in neutral approach parameter between amygdala and sham session was significantly correlated with the bilaterality covariate (*r*(18) = 0.680, *p* = .001), with no such relationship for the approach biases for happy (*r*(18) = -0.143, *p* = .547) or angry (*r*(18) = - 0.270, *p* = .248; **Fig 5c** **and Fig S6b,d**) face stimuli. These findings suggest that BLA-TUS changed the way participants responded to neutral face stimuli. This effect depended on how completely the BLA had been bilaterally stimulated in each participant. We confirmed the robustness of this finding in an analysis free of any assumptions about bilaterality. We used a data-driven approach inspired by voxel-based lesion-symptom mapping in patients(Bates et al., 2003). Binarized post-hoc acoustic field simulations for each participant (in situ I_PA_ > 2W/cm^2^ = 1; I_PA_ < 2 W/cm^2^ = 0) were correlated with p(approach|neutral) across subjects. The resulting map showed voxels where successful TUS stimulation was associated positively with p(approach|neutral) and confirmed that clusters were indeed located bilaterally in the BLA, but stronger in the right hemisphere (**Fig S6g** & Discussion).

### BLA-TUS slows responding to ambiguous facial expressions

One reason for why amygdala stimulation induced changes in choice behaviour selectively for neutral faces may have been because of the ambiguity in their emotional interpretation. In our task, neutral facial expressions are the least clear among the three emotion categories. Following, BLA-TUS, participants increased their rate of approaching neutral faces, effectively treating them more like happy faces. We reasoned that this might have been due to a blurring of the category boundaries between neutral and happy. In healthy populations, neutral faces are often subjectively perceived as slightly positive rather than objectively neutral. We thus examined whether the category boundary between happy and neutral faces was perceived closer for the facial identities used in the task. Subjective rating data collected in our online sample (n = 210) confirmed this for the set of neutral faces included here: neutral face identities were perceived on average more happy and less angry than the mid-point reference (happy and angry ratings given on a sliding scale from 1-4 where the neutral mid-point would correspond to 2.5: *M*_happy_ = 2.54, *M*_angry_ = 2.34, *t*(21) = 2.318, *p* = .031, *d* = 0.494; **Fig 5d**), consistent with our interpretation that happy and neutral emotion categories were closer and thus harder to distinguish.

We reasoned that, if the BLA-TUS effect on choices was driven by further blurring the distinction between happy and neutral emotion categories, participants might need longer to make choices on happy and neutral trials following BLA-TUS. Thus, we next tested for effects of BLA-TUS on RTs. In a mixed-effects linear regression with the predictors TUS condition (amygdala vs. sham), emotion of the displayed face (happy/neutral/angry), and feedback (positive/negative), we found a significant interaction between TUS condition and emotion (χ^2^(1) = 6.485, *p* = .039), with significant RT slowing under BLA-TUS for happy faces (*b* = 0.054, 95%-CI [0.005 0.104], χ^2^(1) = 4.585, *p* = .032) and neutral faces (*b* = 0.051, 95%-CI [0.004 0.097], χ^2^(1) = 4.555, *p* = .033), but not for angry faces (*b* = 0.003, 95%-CI [-0.048 0.053], χ^2^(1) = 0.012, *p* = .914; **Fig 5e**). Selective slowing for happy and neutral, but not angry faces was in line with participants responding more similarly to happy and neutral faces after BLA-TUS. There was no significant interaction between TUS condition and feedback (*b* = 0.003, 95%-CI [-0.014 0.034], χ^2^(1) = 0.117, *p* = .729) and none of the effects in the model were significantly modulated by the bilaterality covariate (all χ^2^ < 3.128, all *p* > .209). Hence, RT slowing for happy and neutral faces seemed to occur non-specifically in all participants who received some degree of BLA-TUS, irrespective of its bilaterality. RT slowing might thus reflect a “prodromal phase” of BLA-TUS effects that only translates into a change in responses when a sufficiently high dose is applied. In summary, BLA-TUS slowed RTs particularly when emotion recognition was challenging and emotions could be more easily mistaken for each other.

### Functional specificity: BLA-TUS leaves learning from feedback unaffected

Finally, we returned to our second hypothesis, that the BLA might also play a causal role in learning. In prior work, the amygdala has been associated with value updating, sensitivity to negative feedback and credit assignment(Costa et al., 2016; Izquierdo and Murray, 2007; Morrison and Salzman, 2010; Rudebeck et al., 2013; Taswell et al., 2021; Wassum and Izquierdo, 2015). For example, updating is slowed and learning from positive feedback reduced following amygdala lesions in macaques(Costa et al., 2016). Therefore, we next tested if amygdala stimulation caused changes in the way participants adapt their choices to the feedback they received. When regressing participants’ repeat/switch responses onto stimulation condition (BLA/sham), feedback on the last trial (positive/negative), mini-block half (first/second), bilaterally stimulated BLA covariate, and all possible interactions, none of the effects involving stimulation condition were significant (all χ^2^< 1.065, all *p* > .301; **Fig5f**). Similarly, beyond changes in the bias to approach neutral faces, there was no change in any other parameter of our computational learning model (Supplementary Results). These results might be due to the type of learning probed by our task. Our task does not require learning stimulus-outcome associations frequently studied in the context of amygdala function(Costa et al., 2016; Izquierdo and Murray, 2007; Morrison and Salzman, 2010; Rudebeck et al., 2013; Taswell et al., 2021; Wassum and Izquierdo, 2015). However, it suggests that BLA-TUS effects are functionally specific, changing responses to emotionally ambiguous stimuli while leaving learning from feedback unaffected.

### Spatial specificity: mIns-TUS reduces GABA, spares approach biases but causes perseveration

We also examined if our effects were specific to the site of ultrasound stimulation, and thus dependent on TUS targeting the BLA, as opposed to a control region. We chose to stimulate the mid-insula (mIns) as our active control region because the transducer stimulation trajectory through the temporal window is highly similar to the BLA trajectory and thus minimized any differences in the subjective experience for participants. Anterior parts of the insula (agranular and anterior aspects of dysgranular insula) show strong anatomical connectivity to the amygdala, while mIns connections to the amygdala are sparser(Aggleton et al., 1980; Amaral and Insausti, 1992; Carmichael and Price, 1995; Friedman et al., 1986; Stefanacci and Amaral, 2000; Turner et al., 1980; Evrard, 2019). Thus, the mIns can be considered part of a related anatomical and functional network, but not part of the same network as BLA.

The mIns was targeted using the same stimulation parameters as BLA (**Tables S4, S6**; I_SPPA_: *M* = 6.6, *SD* = 1.3, range 3.2 – 9.0; **Fig 6a-b**). Post-hoc simulations confirmed that stimulated tissue was located in the mid and ventral parts of the dysgranular and agranular insula (**Fig 6b**) corresponding mainly to cytoarchitectonic areas Ia3 and Id9 from a recent insula atlas(Quabs et al., 2022). Given that we did not miss the insula in any participant, we included all n = 29 in analyses pertaining to insula-TUS effects.

**Figure 6.**
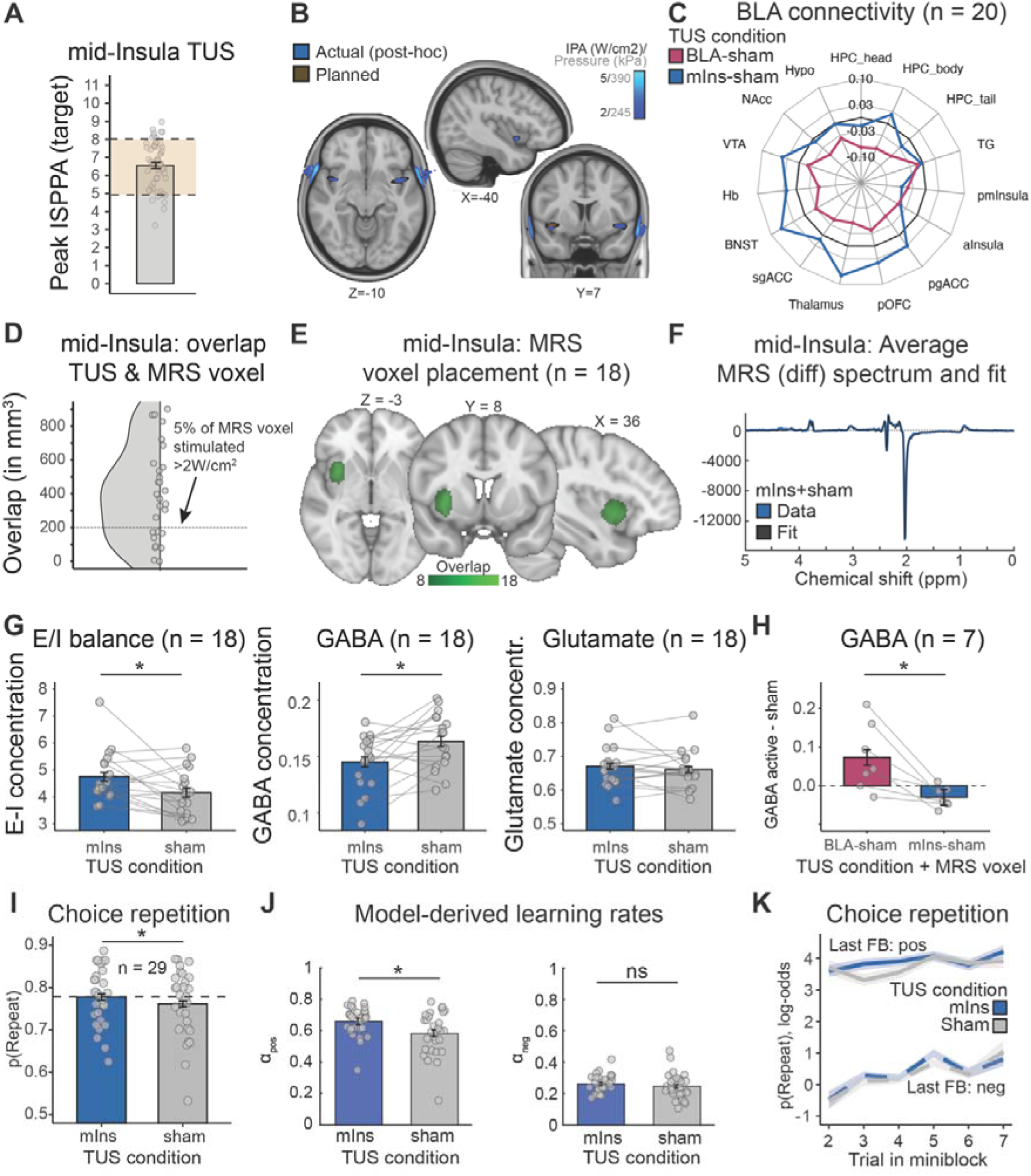
Behavioural and neural effects are specific to BLA-TUS. **A**, The peak achieved in situ I_SPPA_ in the insula in all n = 29 participants as inferred from *post-hoc* simulations (aim: values between 5-8 W/cm2 in the yellow range; achieved: mean of 6.6 W/cm^2^, range 3.2–9.0). **B,** I_PA_ map showing average *post-hoc* simulations of all mIns stimulations in MNI space (n = 29). **C,** Following mIns-TUS, participants’ BLA resting-state connectivity fingerprint was not changed (blue; connectivity for BLA-TUS sessions reported in Fig3b is shown for comparison in pink for the same n = 20 included in Fig3). **D,** MRS voxel placement was challenging due to large vessels. Inspection of overlap between MRS voxel placement and TUS *post-hoc* simulations (>2 W/cm^2^ I_PA_) revealed n = 18 out of all n = 29 participants with sufficient overlap (>5% = dashed line; nine participants had insufficient overlap and are below the cut-off; in one participant, no MRS data was recorded in the sham session; in another participant, the overlap could not be established because, due to a technical error, the transducer location during sonication was not recorded, which made it impossible to run acoustic simulations post-hoc). **E,** MRS voxel placement of n = 18 included participants (n = 36 sessions, sham + mIns) following exclusion of one additional dataset with insufficient MRS data quality and inclusion of one additional dataset originally excluded for poor behavioural performance. MRS voxels were registered to MNI space, binarized, and summed over participants to show overlap. **F,** Average MRS difference spectrum across mIns-TUS and sham sessions (n = 36, two sessions in each of n = 18 participants; for separate fits of mIns vs sham sessions, fits to the off spectrum and further MRS quality metrics, see **Fig S5**). **G,** The E/I balance of the insula was increased following mIns-TUS. This was due to a decrease in GABA, while glutamate was unchanged. Non-parametric tests yielded highly similar results (Wilcoxon signed-rank test: E/I: *V* = 140, *p* = .016; GABA: *V* = 31, *p* = .016; Glutamate: *V* = 93, *p* = .766). **H,** Direct comparison using the intersection of participants with good BLA and mIns MRS signal and sufficient MRS/TUS overlap across all sessions (n = 7, with three sessions each; total: 21 sessions) confirmed that the same TUS protocol caused plasticity effects with opposite polarity in mIns and BLA, increasing GABA in BLA following BLA-TUS but decreasing GABA in mIns following mIns-TUS. **I,** Following mIns-TUS, participants became more likely to repeat their previous response. **J,** This effect was captured by an increase in the model-derived positive, but not negative learning rate parameter. **K,** Illustration of the probability of repeating the previous response following negative (dashed) and positive (solid) feedback. Consistent with learning rate effects in J, this shows a higher probability of repeating choices particularly after positive feedback early in the mini-block (shown in log-odds space for easier visualization).

We first inspected whether the BLA resting-state connectivity fingerprint shown in **Fig 3** was changed following mIns-TUS in the n = 20 BLA participants (**Fig 6c**, **Fig S4**). BLA connectivity was not systematically changed following ultrasound to mIns (*t*(19) = 0.711; *p* = .486). In a mixed-effects linear regression with the predictors TUS condition (amygdala vs. insula vs. sham) and run (run1/run2), we found a significant difference between amygdala and insula (*b* = 0.061, *SE* = 0.017, *t*(114) = 3.61, *p* < .001), as well as between amygdala and sham (*b* = 0.050, *SE* = 0.014, *t*(114) = 3.64, *p* < .001), but no effects of, or interactions with, run.

We next tested whether metabolite changes detected with MRS after mIns-TUS followed a similar pattern to that observed following BLA-TUS, given the same TUS protocol was used in both regions. While MRS signals are easier to record in cortical regions like the insula compared to subcortical regions like BLA, the MRS voxel placement was difficult here too because the insula is innervated by many large blood vessels. Following the same exclusion criteria as for BLA MRS, starting with n = 29 participants, we achieved sufficient overlap between the simulated ultrasound pressure beam and the MRS voxel placement (at least 200 mm^3^/5% overlap between simulated beam thresholded at > 2 W/cm^2^ and MRS voxel) in n = 18 out of 29 participants (see Methods; **Fig 6d**). Out of these n = 18, one participant did not have sufficient MRS data quality. Again, it was possible to add one additional participant who was initially removed due to poor task performance but showed adequate TUS/MRS overlap and MRS quality. In our final sample of n = 18 participants (36 sessions – sham and mIns; overlap of voxel placement: **Fig 6e**, mean spectrogram and fit: **Fig 6f**), we found that mIns-TUS significantly increased the E/I balance, χ^2^(1) = 6.757, *p* = .009, *d* = 0.613 (**Fig 6g**). This was driven by a significant reduction in GABA, χ^2^(1) = 8.207, *p* = .004, *d* = -0.675, but no change in glutamate, χ^2^(1) = 0.678, *p* = .410, *d* = 0.194 (**Fig 6g**).

Qualitatively identical results were obtained using non-parametric tests and when including age, gender, time of day at data acquisition, and simulated pressure beam/MRS voxel overlap as covariates of no interest (**Table S27**; for further MRS quality controls, separate fits for off/diff spectra and mIns/sham sessions, see **Fig S5**). These results suggest that our ultrasound stimulation protocol appeared to be inhibitory (i.e., reducing E/I balance) when applied to the BLA, but consistently excitatory (increasing E/I balance) when applied to the mIns. Finally, we directly contrasted effects of BLA-TUS and mIns-TUS on recorded MRS signals in the intersection of participants included in both the BLA and mIns MRS analyses.

In these people, we required the post-hoc simulations for both mIns and BLA to fall into the respective MRS voxel (>5% overlap as before), and the MRS data quality to be sufficient in both regions and across both active and sham sessions. Only a small subset of seven participants (n = 7, 21 sessions) fulfilled these criteria. In this subset of people, the E/I balance was significantly higher after mIns-TUS than after BLA-TUS in every participant, χ^2^(1) = 30.317, *p* < .001, *d* = 2.081, driven by significantly lower GABA concentration after mIns-TUS compared to BLA-TUS, χ^2^(1) = 12.842, *p* < .001, *d* = -1.354 (**Fig 6h**; **Table S28**), with no significant difference in glutamate concentration, χ^2^(1) = 0.128, *p* = .720, *d* = -0.135 (**Fig S7a**). This direct comparison corroborates the suggestion that the same stimulation protocol had significantly different effects when applied to BLA and mIns, being inhibitory in the former and excitatory in the latter. Consistently across both regions, these metabolite changes were driven by the inhibitory neurotransmitter GABA.

We next analysed the effect of mIns-TUS on behaviour (n = 29). In contrast to the BLA-TUS results, we found no evidence for any effects of mIns-TUS on approach/avoid decisions in our task. To show this, we performed analogous analyses as those done for BLA-TUS, regressing participants’ approach/avoid responses onto the stimulation condition (mIns/sham) and the emotion of the face stimuli (happy/neutral/angry). Neither the main effect of stimulation condition on responses (*b* = 0.002, 95% CI [-0.025 0.029], χ^2^(1) = 0.020, *p* = .888) nor the interaction with emotion (χ^2^(2) = 0.172, *p* = .918) were significant (**Fig S7b**). Similarly, in our winning computational reinforcement learning model, neither the approach bias for happy (*t*(28) = 0.236, *p* = .815, *d* = 0.044), neutral (*t*(28) = -0.010, *p* = .992, *d* = -0.002), or angry face stimuli (*t*(28) = 0.443, *p* = .661, *d* = 0.082) differed between sonication conditions.

To directly contrast BLA and mIns TUS conditions in terms of their effects on approach/avoid decisions, we performed an additional analysis where we included both active sonication conditions, BLA-TUS and mIns-TUS, in the subset of participants with sufficient BLA simulation (n = 20). Consistent with the key effect reported for BLA-TUS vs. sham in **Fig 5**, we again found a significant interaction between stimulation condition (here BLA vs. mIns), emotion, and the bilaterally stimulated BLA volume covariate, χ^2^(2) = 7.499, *p* = .023, reflecting a bilaterality-dependent increase in approach behaviour to neutral faces under BLA-TUS compared to mIns-TUS. This interaction was also significant when including all three conditions (BLA vs. mIns vs. sham) in the model (χ^2^(2) = 11.786, *p* = .019).

Having examined the effect of mIns-TUS on approach/avoid choices, we next turned to effects on RTs. We again found that effects reported for BLA-TUS were region-specific and not present following mIns-TUS. When contrasted against sham, the interactions of mIns-TUS vs. sham with facial emotion (χ^2^(2) = 2.575, *p* = .276) and with feedback (*b* = 0.001, 95% CI [-0.011 0.013], χ^2^(1) = 0.023, *p* = .879) were not significant, providing no evidence for any effect of mIns-TUS on RTs, in line with the absence of mIns-TUS effects on responses. When directly contrasting BLA-TUS with mIns-TUS in the subset of participants with sufficient BLA simulation (n = 20), we found a trend-wise effect for a stimulation condition x emotion interaction on RTs, χ^2^(1) = 4.599, *p* = .100, which became marginally significant when also including the sham condition, χ^2^(1) = 7.940, *p* = .094. This effect again reflected RT slowing for responses to happy and neutral faces for BLA-TUS compared to mIns-TUS and sham. Taken together, we did not find evidence for mIns-TUS changing emotional approach/avoid tendencies or RTs in the way reported for BLA stimulation in **Fig 5**.

In a third and final behavioural analysis for the insula sessions, we tested for effects of mIns-TUS on learning. While BLA-TUS did not show any effects on learning (**Fig 5f**), here, by contrast, we found effects following insula stimulation (**Fig 6i-k**). We regressed participants’ repeat/switch responses on the stimulation condition, feedback received on the previous trial (positive/negative), and mini-block half. We observed a significant main effect of stimulation condition on repeat/switch responses, with more choice repetitions after mIns stimulation (*b* = 0.103, 95% CI [0.007 0.198], χ^2^(1) = 4.404, *p* = .036; **Fig 6i**) regardless of emotion. The interactions of stimulation condition with feedback (*b* = 0.046, 95% CI [-0.037 0.128], χ^2^(1) = 1.169, *p* = .280) and mini-block half (*b* = 0.004, 95% CI [-0.041 0.049], χ^2^(1) = 0.035, *p* = .852) were not significant. As with the amygdala stimulation sessions, we verified the voxel pattern underlying our results with an analysis akin to voxel-based lesion-symptom mappings in patients(Bates et al., 2003). Again, we used binarized *post-hoc* acoustic field simulations for each voxel and participant (in situ ISSPA > 2 W/cm^2^ = 1; I_PA_ < 2 W/cm^2^ = 0). The resulting map showed voxels where successful TUS stimulation positively related to p(repeat) and confirmed that these were indeed located bilaterally in the ventral mid-insula, with stronger relationships in the right hemisphere (**Fig S7d**). When directly contrasting mIns-TUS with BLA-TUS in all participants with sufficient mIns stimulation (n = 29), the main effect of stimulation on choice repetitions was not significant (χ^2^(1) = 2.141, *p* = .143) but became marginally significant when also including the sham condition (χ^2^(1) = 4.908, *p* = .086). This effect again reflected a higher propensity to repeat responses after mIns-TUS compared to both BLA-TUS and sham.

In line with the increase in response repetitions after mIns-TUS, in our winning computational reinforcement learning model, we found significantly larger positive learning rates after mIns sonication compared to sham (*t*(28) = 3.681, *p* = .001, *d* = 0.684; **Fig 6j**), with no significant difference in negative learning rates (*t*(28) = 1.402, *p* = .172, *d* = 0.260; **Fig 6j**). This can be visualised in the raw data by plotting p(repeat) separately per received feedback across trials in a mini-block, which showed a consistent effect that was particularly pronounced in early trials following positive feedback (**Fig 6k**). No such effect was seen when comparing behaviour following BLA-TUS and sham (**Fig 5f**, **Fig S7c**). Taken together, this finding suggests a double dissociation between BLA and mIns-TUS, with BLA-TUS increasing an approach bias towards neutral face stimuli, likely due to its role in resolving emotional ambiguity, and mIns-TUS increasing the propensity to repeat responses irrespective of the previous feedback, likely helping to decide whether to stick with a previous response.

### No side effects or change in mood following a single session of TUS to the amygdala

The amygdala has been consistently implicated in mood disorders. However, given the focus of our study on healthy participants and the brief nature of our TUS intervention, we did not expect any changes in mood in our population. In line with our hypotheses, analyses evaluating mood state showed no consistent changes in mood following BLA-TUS in the n = 20 participants with successful BLA stimulation (for details, see methods: STAI: *t*(19) = 0.164, *p* = .870; PANAS: *t*(19) = -0.173, *p* = .863; and MASQ: *t*(19) = 0.340, *p* = .736; **Fig S8a**) or following insula TUS in the n = 29 participants (STAI: *t*(28) = -1.159, *p* = .255; PANAS: *t*(28) = -0. 634, *p* = .530; and MASQ: *t*(28) = -0.786, *p* = .438; **Fig S8b**).

Similarly, there was no evidence for any difference in reported side effects after sham vs mIns vs BLA stimulation (all False Discovery Rate FDR-corrected p > .05 across the 26 symptoms in the debrief questionnaire; Friedman tests, **Fig S9; Table S29**). Upon more detailed investigation, questions related to scalp sensations showed differences between sham and active (mIns or BLA), but importantly not between BLA and mIns (see Supplementary Results).

## Discussion

The need to better describe the causal role of human subcortical regions such as the amygdala has long been recognized, both in health and in mental illness. Here, we used TUS to test whether the BLA causally contributes to emotional biases or learning from feedback in complex and emotionally ambiguous situations. We found that BLA-TUS increased approach tendencies to neutral faces, the perceptually most ambiguous condition in our task, in a manner that depended on the bilaterality of our stimulation. This effect was captured by a computational reinforcement learning model, with separate bias parameters for each emotion category, which showed an effect only in the parameter that captures approach tendencies of neutral faces. In general, after BLA-TUS, participants treated happy and neutral faces more similarly, corroborated by an overall slowing of reaction times for faces from those two emotion categories after TUS. BLA-TUS also affected measures of neural activity, reducing overall resting-state connectivity of the BLA with monosynaptically connected regions, and decreasing its excitation/inhibition balance (glutamate/GABA ratio) as measured with MRS, suggesting an inhibitory effect of our protocol on BLA activity. These effects were functionally and regionally specific: the identical stimulation protocol applied to mid-insula did not change participants’ propensity to approach the faces, but instead increased their tendency to repeat previous responses, captured by an increase in the positive learning rate. Furthermore, mIns-TUS significantly increased the excitation/inhibition balance in this region, suggesting an excitatory (instead of inhibitory) effect of the same protocol when applied to a different region. Overall, our study provides causal evidence for the role of the BLA in interpreting emotionally ambiguous faces, with TUS leading to an over-generalization of the emotional action bias from happy to neutral faces.

Our emotional learning task posed two computational challenges to participants: they needed to infer the correct response for each mini-block based on positive/negative feedback, and they needed to monitor the emotion of face stimuli presented on the current trial to detect the onset of a new mini-block. In line with previous literature(Volman et al., 2011), participants showed an emotional bias, with more (and faster) approach responses to happy faces than to angry faces. In sham sessions, behaviour towards neutral faces was close to the midpoint between happy and angry faces. However, under BLA-TUS, participants showed an increase in the propensity to approach neutral faces, leading to happy and neutral faces being treated more similarly. We speculate about two possible explanations for this selective effect: the BLA might be particularly needed for processing perceptually and emotionally ambiguous input, e.g., neutral faces, or BLA-TUS might have reduced perception of negative cues contained in ambiguous neutral faces.

Perceiving faces and detecting their emotional expression involves a distributed network of regions, including the fusiform face area and the medial temporal lobe(Carlin and Kriegeskorte, 2017; Kanwisher and Yovel, 2006; Loffler et al., 2005), but also the amygdala(Adolphs et al., 1999; Fried et al., 1997; Wang et al., 2014). The amygdala is similarly involved in detecting emotional expressions in voices(Scott et al., 1997), suggesting it as a cross-modal hub for emotion detection(Janak and Tye, 2015; LeDoux, 2000; Morrison and Salzman, 2010; Pessoa and Adolphs, 2010). The amygdala responds most strongly to fearful, but also to happy and neutral faces(Costafreda et al., 2008; Fitzgerald et al., 2006; Mende-Siedlecki et al., 2013) and groups faces into subjective categories based on perceptual features(Wang et al., 2014) rather than encoding low-level single identities or high-level abstract semantic categories(Cao et al., 2025). The amygdala projects back to early visual cortices and can bias further visual processing(Freese and Amaral, 2005). Hence, interference with amygdala activity might impair the classification of stimuli into broad emotional categories, especially for perceptually ambiguous input(Naaz et al., 2019; Wang et al., 2017). In our task, neutral faces were the most ambiguous category of faces. Online participants rated these faces as slightly more happy than angry, suggesting that happy and neutral faces could be more easily confused which may have led to more similar approach patterns to happy and neutral faces after BLA-TUS. Furthermore, reaction times to both happy and neutral faces were slowed following BLA-TUS, corroborating the interpretation that these two emotions became harder to distinguish. Hence, TUS might interfere with the BLA’s role in resolving ambiguity, a role critical for perceptual decisions such as the classification of facial emotions, but likely extending also to value-based decisions(Hsu et al., 2005; Madarasz et al., 2016; Pauli et al., 2012; Sharot et al., 2007).

An additional, not mutually exclusive explanation is that increased approach behaviour to neutral faces was caused by diminished perception of negative cues contained in ambiguous neutral faces. Previous literature has suggested a functional asymmetry between the left and right amygdala, with the left hemisphere predominantly processing positive information (such as happy faces), and the right hemisphere predominantly processing negative information (such as angry faces)(Ocklenburg et al., 2022; Palomero-Gallagher and Amunts, 2022). TUS to (particularly the right) BLA might have impaired participants’ ability to perceive the negative aspects of neutral (i.e., ambiguous) faces, treating them more like happy faces and showing an emotional approach bias towards them. This finding is in line with a case report of increased positivity bias when electrically stimulating the right BLA(Bijanki et al., 2014) and with our exploratory data-driven analyses. In this analysis akin to lesion symptom mapping, approach tendencies to neutral faces were associated with TUS in bilateral BLA, but the association was stronger in the right.

In sum, we found that bilateral BLA-TUS led to an increase in approach behaviour to neutral, i.e., emotionally ambiguous faces, and a RT slowing for neutral and happy faces, with these emotional categories treated more similarly after BLA-TUS. Our study shows that BLA-TUS, when used within currently established guidelines(Aubry et al., 2024), is an efficient and safe non-invasive method to perturb neural activity in a temporally and spatially precise way, with effects on resting-state connectivity, metabolites, and behaviour. We thus extend recent studies that established the feasibility of TUS to the human amygdala and found initial evidence for changes in subjective experience(Barksdale et al., 2025; Chou et al., 2024; Hoang-Dang et al., 2024; Mahdavi et al., 2023; Spivak et al., 2025), memory performance(Doss et al., 2025), and neural activity(Gorka et al., 2024; Kuhn et al., 2023). While most of these studies focused on targeting the amygdala unilaterally, here we show that bilaterality might be an important factor for modulating emotional processing and choice behaviour, and we provide direct proof of target engagement using metabolite and resting-state connectivity measures. We note that, while the *in situ* location of the pressure beam can be well predicted from acoustic simulations, the precise values of pressure derived from simulations have considerable uncertainty around them because the acoustic properties of bone are not well understood(Albelda Gimeno et al., 2019; Martin et al., 2024). For that reason, we supplemented our analyses with data-driven approaches inspired by voxel-based lesion-symptom mapping (VLSM) that merely relied on binarized simulated pressure fields, and which confirmed our key findings.

TUS applied to an active control region, the ventral mid-insula, led to markedly different changes in neural activity and behaviour. mIns-TUS did not change approach behaviour or RTs but led to an increase in the probability of repeating a previous response, captured by increased positive learning rates. Previous studies have suggested a role of the insula (though most notably the anterior insula) in processing negative feedback(Gueguen et al., 2021; Palminteri et al., 2012) and adjusting behaviour(McGuire and Kable, 2015; Trier et al., 2025). Importantly, we found an increase in response repetitions independently of the previous feedback, reflected in positive, but not negative learning rates. In the context of our task, not only negative feedback, but also surprising perceptual input suggesting the onset of a new mini-block can lead to changes in responses. We interpret our finding as mIns-TUS attenuating participants’ sensitivity to such ambiguous perceptual evidence, leading to a higher propensity to repeat responses and less false-positive switches to the other response option. While the insula has been associated with behavioural flexibility and adjustment to subtle changes in the environment(Sallet et al., 2020; Wittmann et al., 2020), mIns-TUS might make behaviour more rigid and less sensitive to such influences.

Intriguingly, MRS measurements suggested an opposite effect of BLA-TUS and mIns-TUS on E/I balance: while BLA-TUS tended to decrease E/I balance in the BLA, the very same stimulation protocol applied to the mIns significantly increased E/I balance in the mIns. While the possibility of different effects has been proposed for different parameter settings, e.g. different PRFs or duty cycles(Murphy et al., 2024), this finding suggests that the same TUS protocol can have both inhibitory and excitatory effects on different brain regions, implicating that TUS protocols cannot be described as uniformly “excitatory” or “inhibitory”. Instead, their effect might vary depending on the targeted brain region’s intrinsic connectivity, cell (including neuron) type distribution, receptor densities, or other structural or functional properties. For example, differences might arise from the recurrent nature of a region’s intrinsic connectivity. The BLA supports highly recurrent local circuitry, ideal for rapid associative learning and emotional memory encoding. In contrast, the midventral insula exhibits extensive, but more distributed, within-region connectivity that underpins integration of sensory, interoceptive, and affective signals rather than localized recurrent dynamics. Both regions have overall more excitatory than inhibitory cells e.g., the amygdala has ∼80% excitatory cells and only a smaller proportion of inhibitory neurons(Vereczki et al., 2021; Yu et al., 2023), which is similar for the insula(Carmichael and Price, 1994; Evrard, 2019; Quabs et al., 2022). Differences in cytoarchitecture and differential expression of ion channels might also play a role. For example, Piezo, TRP and K2P channels—key mechanosensitive ion channel families that likely mediate TUS effects(Prieto and Maduke, 2024)—are dense in the amygdala, but not well described for insula(Raha et al., 2023).

Consistent with the idea that many factors might contribute to whether excitation of inhibition is observed, MRS signals in other regions following TUS in humans have not led to consistent results. For example, TUS has been found to exhibit no effect on MRS measurements in anterior cingulate cortex, but to reduce GABA concentrations in posterior cingulate cortex, leading to net increased in E/I balance in line with our finding in the mid-insula(Yaakub et al., 2023). Similarly, effects on resting-state connectivity(Atkinson-Clement et al., 2025, 2024; Gorka et al., 2024; Sanguinetti et al., 2020; Yaakub et al., 2023) and M1 excitability(Zadeh et al., 2024; Zeng et al., 2022; Zhang et al., 2023) have been variable across studies. Notably, in our study, participants were playing the task while the MRS measurements were acquired, and different effects might have occurred if participants had been at rest. Thus, more systematic investigations into parameter settings, brain region properties, and functional state will be needed to fully understand the direction of effects on a physiological and behavioural level.

One weakness of our study is the considerable variability in the exact targeted area across participants. We chose to use an ultrasound transducer with a low fundamental frequency of 212 kHz, which came with a relatively short axial (depth) steering range. Lower fundamental frequencies lead to larger focal volumes for a fixed transducer aperture and, in the case of 212 kHz, matched the size of the BLA target region better than, e.g., a 500 kHz transducer. Also, 212 kHz allowed for better transmission of energy through the skull.

However, the short axial steering range meant that we could not accommodate a thicker and more flexible coupling medium such as a gel pad or water balloon, which would have reduced our axial depth in a way that would have moved the BLA out of range (see Methods). In future studies, one might use a transducer with a deeper steering range such that a more flexible coupling approach with full control over the angle can be added.

Emotion and learning processes are key functions affected in mood disorders(Admon and Pizzagalli, 2015; Browning et al., 2019; Eshel and Roiser, 2010; Harmer et al., 2009a; Korn et al., 2013). Understanding their causal underpinning in the human brain is therefore of vital importance for informing (neurally and functionally) targeted treatments given a patient’s specific impairments. Yet, learning and emotion processes are often associated with structures located deep inside the brain(Cardinal et al., 2002; Murray et al., 2011; Pizzagalli and Roberts, 2022). Thus far, causal investigations in this domain have largely been limited to animal work. Here we hoped to begin to bridge the gap between causal studies in animals and correlational work in humans to help us understand these key processes and the causal contribution of deep subcortical nuclei(Murphy and Fouragnan, 2024). Our insights may have relevance for work in patients with depression, anxiety, and other brain disorders in which affected circuits are located deep in the brain and can now be directly targeted for neuromodulation using low-intensity ultrasound(Bubrick et al., 2023, 2022; Kubanek, 2018; Riis et al., 2024b, 2024a). Specifically, these disorders are accompanied by neural changes in the amygdala(Drevets, 2003; Drevets et al., 1997; Murray et al., 2011) as well as changes in learning and emotional processing(Admon and Pizzagalli, 2015; Eshel and Roiser, 2010).Yet, until recently, there has been no method in humans to directly and causally link changes in amygdala neural processes to changes in emotional learning and behaviour. One interpretation of our findings could be that amygdala stimulation increases people’s optimism bias for neutral faces in a way that make them process these faces more similarly to happy faces (Hales et al., 2014; Harmer et al., 2009b; Korn et al., 2013; Willinger et al., 2022). Patients with altered amygdala processing might particularly struggle with ambiguous situations because they struggle to disambiguate the subtlety of emotions in such situations(Everaert, 2021; Everaert et al., 2017; Hirsch et al., 2016). While we cannot provide a direct link between functional impairments experienced daily by patients, we provide evidence that resolving ambiguity causally relies on the basolateral amygdala. This finding might have important implications for mood disorders, which might arise from a lack of disambiguation that attenuates or exacerbates emotional and learning biases. This should be directly explored in future work in patient populations.

## Methods

### Participants

A total of thirty-six healthy volunteers took part in this study. The study was approved by the University of Oxford CUREC (Ref: R85030/RE001). All participants provided written informed consent. Screening criteria included no history of neurological or psychiatric disorder, normal or corrected-to-normal vision and suitability for undergoing MRI scanning and ultrasound stimulation (see Supplementary Methods for full TUS screening form). Three participants withdrew from the study before data collection was complete, one due to discomfort during TUS (heating sensation on the skin underneath the transducer), two due to discomfort during 7T-MRI (nausea). An additional four participants were excluded from analyses of the behavioural data due to their inability to follow the task instructions in at least one session (flat learning or failure to perform incongruent actions). The final sample thus included n = 29 healthy volunteers (15 female), aged 19 to 40 years (M = 24.517, SD = 4.629). Participants were paid £200 including a £50 bonus for completing all four study visits.

### Study design and overall timeline

Participants attended four sessions (**Fig 2a**). During the first visit (45 minutes of which 30 minutes were MR scan time), we acquired T1-weighted and PETRA (pointwise encoding time reduction with radial acquisition; a variant of zero-echo time or ZTE) scans on a Siemens 3T Prisma scanner. These images were required for ultrasound planning (**Fig 2b**), which was performed between session 1 and 2 (for details on MR sequences and simulations, see separate sections below). In visits 2–4, participants read the task instructions, performed a practice version of the task, changed into MR-compatible scrubs, and received TUS targeting bilaterally either the basolateral amygdala (BLA), mid-insula (mIns, as an active control region), or sham (counterbalanced order). Following TUS, participants immediately transitioned into the ultra-high field (7T) MRI scanner for a duration of approximately 1.5– 2h. During this time, we acquired task-fMRI, resting-state fMRI, and magnetic resonance spectroscopy (MRS) data. The order in which the scans were acquired is shown in **Fig 2e**. Following the MRI, participants changed back into their clothes and filled in a TUS debrief and some shortened mood questionnaires (for full questionnaires, see Supplementary Methods). 24h after each TUS session, participants were again asked to fill in the TUS debrief in case of any changes (for full debrief, see Supplementary Methods and **Tables S29, S30**, and **Fig S9**). The total session duration for sessions 2–4 was 2.61–3.91h (*M* = 3.28h).

### MRI data acquisition

In visit 1, a T1-weighted magnetisation-prepared rapid gradient-echo (MPRAGE) sequence with GRAPPA acceleration factor 2 (repetition time (TR) = 2300 ms; echo time (TE) = 2.26 ms; flip angle: 8°; voxel size: 1 x 1 x 1 mm^3^) and a PETRA sequence (TR = 3.61 ms; TE = 0.07 ms; flip angle: 1°; slices per slab: 320; voxel size: 0.75 x 0.75 x 0.75 mm^3^) were acquired on a Siemens Magnetom 3T Prisma scanner. DIS3D bias correction was performed on the PETRA images.

For visits 2–4, the same scan sequence was followed in each session. Following a localizer, a T1-weighted structural scan as well as multiple functional scans (task-fMRI and resting-state-fMRI), and two magnetic resonance spectroscopy (MRS) voxels were acquired on a MAGNETOM 7T Plus MRI scanner (Siemens Healthineers, Erlangen Germany) using a 8Tx32Rx-channel head coil (Nova Medical Inc, Wilmington, MA, USA). The order and average post-TUS timing of the scans is shown in **Fig 2e**. The structural T1-weighted scan (3:46 minutes) was based on an MPRAGE sequence with GRAPPA acceleration factor 3 (TR = 2600 ms; TE = 3.18 ms; flip angle: 5°; voxel size: 1 x 1 x 1 mm^3^). Task-fMRI (∼25 minutes) and resting-state fMRI (two times 7:16 minutes) scans consisted of T2*-weighted multi-band echo planar imaging (EPI) images (TR = 2140 ms; TE = 20 ms; flip angle 69°; voxel size: 1.5 x 1.5 x 1.5 mm^3^; T > C30.0 angulation; number of slices = 78; multiband acceleration factor = 2; PAT acceleration factor = 2; interleaved slice acquisition; anterior-to-posterior phase encoding) based on the multi-band accelerated EPI sequences from the CMRR package(Moeller et al., 2010). In addition, we collected a single volume of the same sequence with 128 slices (whole-brain reference scan, 30 seconds) that was used to aid the registration. To correct for distortions introduced by the inhomogeneous B0 field, we collected a gradient-echo B0 field map (GRE; TR = 620 ms; TE1 = 4.08 ms; TE2 = 5.1 ms; flip angle 39°; voxel size 2 x 2 x 2 mm; 74 slices; 2 minutes duration). For the single-voxel MRS scans, we used a MEGA-edited semi-LASER sequence, available from the CMRR Spectroscopy Package, suited for measuring GABA concentration(Klomp et al., 2009; Mescher et al., 1998; Mullins et al., 2014). Sequence parameters were a voxel size of 20 x 14 x 20 mm^3^; TE = 78 ms; integrated VAPOR water suppression and OVS (VAPOR bandwidth = 150 Hz); 130 Hz bandwidth editing pulse, centred around 1.9 ppm (edit-on condition) and at 7.5 ppm (edit-off condition). We used different number of spectra for both voxels to compensate for lower signal in the amygdala (BLA: TR = 4 sec.; 192 averages, i.e. 96 each with edit-on and edit-off; total run time of 12 minutes 48 seconds; mIns: TR = 5 sec.; 128 averages, i.e. 64 each with edit-on and edit-off; total run time of 10 minutes 40 seconds). A water unsuppressed reference was collected for each voxel. Shimming was achieved using one-step (linear 6-projection) FASTESTMAP from the CMRR spectroscopy package(Gruetter and Tkáč, 2000). See **Table S9** for the MRSinMRS reporting checklist(Lin et al., 2021). The average total scan duration of an entire session was 100 minutes.

### Emotional approach-avoidance learning task

Both online and in-person participants (sessions 2–4) played the same newly designed emotional approach-avoidance learning task. The task incorporated two critical features: (1) participants were required to perform affect-congruent and affect-incongruent responses, testing their ability to override automatic emotional biases(Volman et al., 2011), and (2) it involved learning correct responses from feedback, with frequent updates in the required response reflecting the dynamic and changeable emotional contexts often encountered in real-life situations(Behrens et al., 2007; Chau et al., 2015; Costa et al., 2016).

On each trial, participants saw a face stimulus (from the Chicago Face Database (CFD)(Ma et al., 2015)) and had to decide to approach or avoid it by pressing one of two buttons on a button box (**Fig 1a, left**). Trials were structured into mini-blocks of 4–7 trials. Within a mini-block, all faces displayed the same emotion (either happy, neutral, or angry), and the correct response was either to approach or to avoid all these faces (**Fig 1a, right)**. Participants had to learn the correct response from trial-and-error using the positive/negative feedback presented at the end of each trial. Feedback was probabilistic, with 80% valid and 20% invalid feedback. When participants pressed the button to approach a face, the face grew bigger on the screen (duration: 1.5 s), providing visual feedback matching the approach action. Vice versa, for avoid actions, the image shrank (duration: 1.5 s). If participants failed to respond within 2 s of the face presentation, they saw an exclamation mark, reminding them to respond. If they still had not responded within 4 s from the face presentation, the trial was counted as a missed trial and the next face was shown. The feedback (green tick or red cross) was shown on the screen for 1 s. For the in-person MRI task version (Exp1), an inter-trial interval which displayed a fixation cross was added before each face stimulus for a duration of 2 s + jitter (0.5 s on average, range 0.2–0.8 s) minus the RT on the previous trial. A total of 22 CFD models were included for each of the three emotion categories. There was no overlap between emotions in terms of the models used, meaning a total of 66 racially diverse faces were used in the task.

In each TUS session (for in-person participants of Experiment 1) and the online session (Experiment 2), participants completed 528 trials of the task, split into four blocks of 132 trials each. Each block comprised eight mini-blocks for each of the three emotions (four with approach as correct response, four with avoid as correct response). Consecutive mini-blocks never featured the same emotion. At the start of the session, participants were given written task instructions and performed a short practice block (64 trials) of the task. They also answered a short series of multiple-choice questions on the task and were only allowed to proceed upon successful completion of these questions. In the main part of the experiment, in-person participants in Experiment 1, following the ultrasound stimulation, performed the first two blocks of the task during task-fMRI recordings and the latter two blocks during MRS image acquisition. Online participants in Experiment 2 played four blocks of the task online.

### Acoustic simulations of ultrasound beam trajectories

The T1-weighted structural and PETRA images acquired from each in-person participant in the first session were used for individualised ultrasound planning. Participant-specific target, skin and transducer coordinates were planned in T1 space in Brainsight and then brought into PETRA space. To build a model for energy absorption through the skull in our simulations, we followed a previously established procedure(Miscouridou et al., 2022) of creating a “pseudo-CT” (https://github.com/ucl-bug/petra-to-ct). We debiased the PETRA using the N4ITK MRI bias correction implemented in the 3D Slicer tool(Fedorov et al., 2012), histogram-normalised the image (to set the mode of the soft-tissue distribution to 1), and segmented it into background, soft-tissue, and skull-tissue using SPM12(Penny et al., 2011). We then built a pseudo-CT by setting the image background to values of -1000 (Hounsfield units), setting voxels classified as soft tissue to a value of 42, and mapping the values of voxels classified as skull linearly to a range of 42–3275. This mapping was established as part of another study performed at Oxford using data from three participants for whom a CT scan and a PETRA scan from the same Oxford 3T MR scanner was available. Finally, we linearly converted Hounsfield units into mass density (kg/m^3^) based on a CT scan of a CIRS Model 062M Electron Density Phantom with known mass densities that was scanned with the same CT scanner. An example pseudo-CT image is shown in **Fig2b**.

We performed simulations of the acoustic pressure field and the temperature field in the k-Plan software, which is a wrapper around the kWave toolbox(Treeby and Cox, 2010). Participant-specific coordinates were defined for the basolateral amygdala (BLA) and mid-insula (mIns) in Brainsight based on a combination of each participants’ brain anatomy and a BLA mask in MNI space that was projected back into subject space. The grid spacing was 1.21 mm, which corresponded to 6 points per wavelength in soft tissue (total grid size 239 x 239 x 239 mm). We used 15,974 time steps of 33.70 ns (total duration: 538 μs), which corresponds to a CFL of 0.04 (reference sound speed: 1509 m/s). Each participants’ TUS stimulation parameters were individually tailored to optimise the energy delivery to the intended target while minimizing skull-induced attenuation and staying within established safety guidelines (transcranial mechanical index (MItc) < 1.9, thermal dose CEM43 < 0.25)(Aubry et al., 2024). The free-field I_SPPA_ (setting on the transducer power output) was adjusted for each participant to reach an intensity of 5–8 W/cm^2^ at the focus in the target region (free-field I_SPPA_: M = 30.5, range 15.2 – 48.0 W/cm2). Acoustic planning was repeated iteratively with slightly modified target coordinates until all safety conditions were met, and the beam was centred on the desired target and reached the intended I_SPPA_. In all our plans, for all participants, we achieved an MItc < 1.8 and all CEM43 < 0.08. On average, the stimulation depth was set to 52.5 mm for the BLA (range 45.7–58.4) and 34 mm (range 32– 40) for the mIns. The average skin location and target location are shown in **Fig 2c**.

After data collection, we repeated acoustic simulations based on the actual transducer and target coordinates recorded via Neuronavigation during TUS (stream2file functionality of Brainsight). The actual transducer position was taken to be the median x/y/z coordinates recorded during the full 80-second stimulation duration (for TUS protocol, see below). We report values for both *ex-ante* and *post-hoc* simulations in **Tables S3-S6**. In *post-hoc* simulations based on the empirical transducer and target coordinates recorded with neuronavigation during TUS, the MItc exceeded 1.9 in three cases (MItc = 1.97, 2.03 and 2.11, all mIns; Table S6). In all cases, the CEM43 was below 0.13. For one participant, due to technical difficulties, no coordinate was recorded for the BLA-TUS session; and for another participant, no coordinate was recorded for the mIns session. In analyses involving intensity estimated from acoustic simulations, these participants were thus not included.

### TUS stimulation procedure and protocol

During sessions 2–4, following consent, task instructions and a change of clothes into MR-compatible scrubs, we used neuronavigation software (Brainsight v2.5; Rogue Research Inc., Montréal, Québec, Canada) to register the participant’s head to their T1 image acquired in session 1 to determine the correct placement of the transducer relative to the participant’s head. The section of the hair underneath the transducer was prepared with ultrasound gel (centrifuged at 3000rpm for 3–5 minutes prior to use to remove air bubbles from the liquid gel) in a way that follows ITRUSST-recommended guidelines and reduces the risk of air trapped in the hair when the transducer is coupled to the participant’s head(Murphy et al., 2025a).

The bowl shape of the transducer was filled with a gelpad which was created using a custom-made 3D printed metallic mould of the transducer bowl. To create each gelpad, the mould was heated and a chunk of gelpad cut off from an Aquaflex gelpad (Parker Laboratories Inc., ∼21-22g) was placed in the heated mould until it slightly melted which allowed it to set into the correct shape. It was then left to cool down and stored for the session. During each session, a small amount of liquid Aquaflex ultrasound gel was applied to the bottom of the transducer bowl before the moulded gel pad was inserted. The liquid gel ensured that the gelpad stayed in place and that no air was trapped underneath the gelpad. A small amount of liquid Aquaflex ultrasound gel was also applied to the surface of the gelpad to ensure good coupling to the participant’s head. As above, all liquid ultrasound gel was centrifuged and the moulded gelpad was inspected for air bubbles before its use. The gelpads we used only filled the cavity of the transducer but did not extend beyond it. Thus, the transducer was filled, but then placed directly onto participants’ skin, without another medium in-between the skin and the transducer edge except ultrasound gel. Hence, the angle of incidence of the pressure beam onto the skull was largely determined by the shape of participants’ head. Despite our best efforts at the planning stage, in some participants, we had a choice to be at an angle of incidence away from 0°, or to not couple to the skin well. In such cases, we accepted imperfections in our targeting (which ended up meaning that n = 20 out of n = 29 participants could only be included). In future studies, we would advise using a transducer with a deeper steering range such that a more flexible coupling approach with full control over the angle of incidence (e.g. a thicker gelpad or water membrane) can be used.

In our sessions, for both BLA and mIns, the transducer filled with the gelpad was coupled to the skin around a temporal lobe window. During stimulation, we recorded the location of the transducer using Brainsight’s stream2file functionality. We used a 212 kHz NeuroFUS Pro transducer (CTX-212-4ch, diameter 64 mm, central frequency 212 kHz, steering range 32.4 – 58.4 mm; SonicConcepts, Seattle, USA) part of the NeuroFUS Pro system (Brainbox Ltd., Cardiff, UK). This transducer is identical to the standard 250 kHz NeuroFUS Pro transducer in terms of it basic features and geometry, but it has a slightly lower fundamental frequency because it was custom built at the time of purchase to achieve our desired steering range (range 32.4—58.4 mm; **Table S1**).

We employed a published 5 Hz patterned protocol(Yaakub et al., 2023; Zadeh et al., 2024; Zeng et al., 2022) which consisted of 212 kHz stimulation in pulses of 20 ms (rectangular shape; no ramping) within a pulse-repetition interval 200 ms (5 Hz pulse-repetition frequency; 10% duty cycle) and in total 400 cycles (80 seconds; **Table S2**). This protocol has previously been used to change motor cortical excitability and resting-state/MRS signals in humans for up to 60 minutes following stimulation(Yaakub et al., 2023; Zadeh et al., 2024; Zeng et al., 2022). Note, however, that the transducer’s fundamental frequency was lower (212 kHz instead of 500 kHz) and we likely achieved higher intensities than in the previous studies.

In sham sessions, we followed the same procedures except that the TPO output I_SPPA_ was set to 0 W/cm^2^, ensuring that no energy was transmitted. To replicate the auditory experience of active sessions during the 80-second stimulation interval, a bone-conducting earphone was placed next to the transducer onto the participant’s head, and a custom masking sound generated to mimic the audible part of the active ultrasound using a 5Hz PRF, as in (Yaakub et al., 2023) was played for 80 seconds during the sham stimulation. On average, participants were above chance at recognizing an active session as an active session: Bang Index, BI (Active): 0.638, 95%-CI [0.469, 0.807])(Bang et al., 2004), likely due to somatosensory confounds (such as tingling and warmth), which a sham sound cannot mimic, and which are stronger for lower fundamental frequencies(Kop et al., 2025). However, importantly, their ability tell that a sham session was a sham session was at chance: BI (Sham): 0.103, 95%-CI [-0.256, 0.463]). Also, they could not distinguish between the two types of active sessions— target (BLA) and active control (insula) sessions (Wilcoxon signed-rank test for reported certainty in identifying stimulation type for BLA vs insula TUS: W = 98, *z* = 0.12, *p* = .901). Also, it is worth noting that all behavioural and MR measurements of interest were not recorded during TUS, but in the minutes and hours that followed. They are therefore unlikely to be confounded by the experience.

### Covariates for quantifying stimulated tissue

Post-hoc simulations were used to quantify how well we had succeeded in stimulating our intended targets. To extract a quantitative continuous measure, we first calculated the volume (in mm or voxels) within our BLA target mask where we had exceeded a value of 2 W/cm^2^ I_PA_. While this is a somewhat arbitrary cut-off, any other choice of threshold naturally leads to highly correlated covariates. The mask used was the sum of the basal (B + AB/BM) and lateral (LaD, LaI, LaV) masks for the BLA(Klein-Flügge et al., 2022). From the stimulated volume in each hemisphere, we then constructed a covariate that captured how much tissue had been stimulated bilaterally: by computing the minimum volume across both hemispheres. The volume of the entire BLA was 3,616 mm^3^ (left BLA: 1,912 mm^3^; right BLA: 1,704 mm^3^). Thus, the volume achieved in the less stimulated hemisphere could range from 0 (both hemispheres missed) – 1,704 (100% hit in both hemispheres, but summed bilateral volume is twice this number). We achieved the following minimum volume stimulated > 2 W/cm^2^ in mm^3^ in our full cohort of n = 29 participants: *M* = 651, *SD* = 424, range 5–1,369. For one participant, due to technical difficulties, the coordinates were not recorded, and this participant was excluded from all analyses involving this coordinate.

### Behavioural regression analyses

Given that participants showed no response on some trials (per session: mean = 4.2, median 1, range 0–100), this may have prevented them from adequately learning the correct response in a given mini-block. We therefore excluded mini-blocks with two or more trials without a response (trials excluded per session: mean 7.86, median 0, range 0–144). We fit mixed-effects logistic regression models (approach/avoid responses) and linear regression models (to reaction times) using the lmer() and glmer() functions from the lme4 package(Bates et al., 2015) in R. We computed Type-3 sums-of-squares *p*-values from Wald chi-square tests using the Anova() function from the car package. All categorical variables were sum-to-zero coded to reduce nonessential multicollinearity between main and interaction effects. All continuous variables were z-standardized for numerical stability. All regression coefficients can thus be interpreted as standardized regression coefficients. To interpret results involving factors with more than two levels (i.e., emotion with three levels), we refit the respective models with each possible combination of two levels only. The full model results for all regressions reported in the manuscript, are shown in **Tables S9-S19**. In all plots showing within-subjects comparisons, we display standard-errors computed with the Cousineau-Morey method(Morey, 2008) adjusting for between-subjects variance.

### Computational reinforcement learning models

We fit a series of increasingly complex reinforcement learning models to participants’ approach/avoid responses. These models were based on a Q-learner that learned the value of performing approach and avoid actions for each mini-block (with mini-blocks treated as separate stimuli). The action value Q of action a for stimulus s on trial t was updated based on the difference between the outcome r (+1 for correct, -1 for incorrect) and the action value from the previous trial (i.e., the reward prediction error) using the delta learning rule:

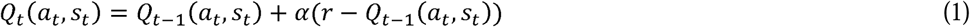

The learning rate α was a free parameter that was fitted for each participant. It determined how quickly action values are updated based on recent feedback. No such learning took place on trials on which participants forgot to make a response. Action values *Q* were then converted into action probabilities p using a softmax transform:

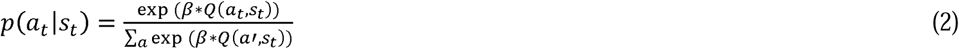

The inverse temperature β was a free parameter fitted for each participant that scaled how deterministic vs. stochastic participants’ choices followed from their learned action values.

The base model **M1** only featured a learning rate α and an inverse temperature β for each participant. In **M2**, we used separate learning rates α_POS_and α_NEG_ for positive and negative outcomes. In **M3**, we added a common general approach bias γ(Huys et al., 2011) that reflected individual differences in participants’ overall propensity to show approach vs. avoid responses to the action values. We refer to these “biased” action values as action weights *w*, which then formed the input to the softmax transform:

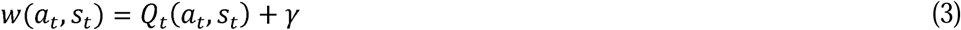

In **M4**, we replaced this general approach bias by emotion-specific biases γ_HAPPY_, γ_NEUTRAL_, and γ_ANGRY_ that reflected that fact that participants showed an emotion bias, with an increased tendency to approach happy (relative to neutral) faces and an increased tendency to avoid angry (relative to neutral) faces(Huys et al., 2011):

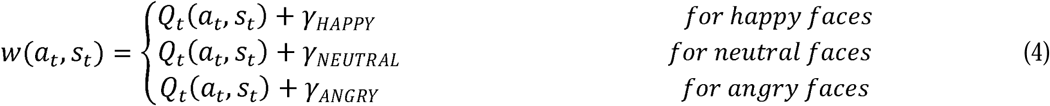

We constrained inverse temperatures to be positive using an exponential transform and learning rates to the range [0, 1] using the inverse-logit transform. In line with recent recommendations(Valton et al., 2020), we fitted each session of each participant separately. Models were fit using the CBM toolbox(Piray et al., 2019) in MATLAB. Fitting involved three steps: First, the data of each session of each participant was fit individually. The CBM toolbox uses an iterative expectation maximization algorithm that used the Laplacian approximation of the parameters’ posteriors. Second, we fitted all participants’ data (separately for BLA/mIns/sham sessions) using a hierarchical Bayesian inference procedure. In this approach, group-level parameters were iteratively fitted using mean-field variational Bayes. Group-level parameters then served as hyperpriors for individual participants’ parameters. Such a hierarchical estimation leads to more robust estimates and reduces the chance of overfitting. Third, we fitted all candidate models in a single step that iteratively determined the probability that the data of a given participant X was best fitted by a certain model Y, and then fitted the group-level parameters of model Y based on only those participants whose data was best characterized by this model (i.e., participants were weighted proportionately by their model responsibility scores). Such an approach allows for hierarchical Bayesian inference (with participants constraining each other’s parameter values) and random-effects model selection (allowing for the possibility that that different participants are best fitted by different models). For Bayesian model selection(Rigoux et al., 2014; Stephan et al., 2009), we then computed the model frequency and protected exceedance probability of each model. We selected the model with the highest exceedance probability as the winning model. We compared fitted parameters from the hierarchical Bayesian inference step using paired t-tests.

### Data-driven spatial mapping linking behaviour and TUS

As a proxy of the bilateral stimulation dose, we used the minimal number of voxels stimulated in either the left or the right BLA using an anatomical mask(Klein-Flügge et al., 2022). This principled approach is not able to detect whether stimulating certain subregions of the amygdala and/or one of both hemispheres was particularly predictive of behavioural effects. We thus adopted a data-driven approach inspired by voxel-based lesion-symptom mapping (VLSM)(Bates et al., 2003) and computed correlations between the binarized simulated intensity (I_PA_ > 2 W/cm^2^ = 1; I_PA_ <= 2 W/cm^2^ = 0) at each voxel, based on the *post-hoc* simulations and the difference in behaviour between active and sham sessions. We used Point biserial correlations as a measure of associations between TUS intensity (binary) and behavioural effects.

### Analyses of debriefing data

At the end of each of the three stimulation sessions, participants rated the severity (1 = Absent, 2 = Mild, 3 = Moderate, 4 = Severe) and the perceived relation to TUS (0 = N/A, 1 = Unrelated, 2 = Unlikely, 3 = Possible, 4 = Probable, 5 = Definite) for 26 symptoms (**Table S29**). Analyses included only symptoms rated as at least mild and perceived as related to the stimulation (relation > 2). For each symptom we ran non-parametric Friedman tests to compare severity across stimulation conditions. False discovery rate (FDR) correction was applied to control for multiple comparisons.

### fMRI resting-state preprocessing

Preprocessing of fMRI resting-state data was performed using FSL’s fMRI Expert Analysis Tool (FEAT v6.0). We performed brain-extraction on the T1 scans using Synthstrip(Hoopes et al., 2022) and performed B1-bias field correction on the T1 and all functional scans using FSL’s FAST tool. The other preprocessing steps were performed using FSL’s FEAT. These steps involved motion correction using MCFLIRT(Jenkinson et al., 2002), linear registration to structural and non-linear registration to standard (MNI152) space using FLIRT and FNIRT(Andersson et al., 2007; Jenkinson et al., 2002), unwarping using the field map to correct for B0 field distortion, spatial smoothing (Gaussian kernel of 5 mm FWHM), and temporal high-pass filtering (cutoff of 100 s). To aid the registration, we used the whole-brain functional reference scan as an intermediate high-resolution image. Furthermore, we used a brain mask based on the twice-dilated T1 brain mask (brought into functional space) for the brain extraction of the functional images.

We created 33 slice-specific nuisance regressors based on heart-rate and respiratory recordings using FSL’s Physiological Noise Model (PNM)(Brooks et al., 2008), comprising the cosine and sine of basic cardiac and respiratory regressors modelled with an order of 4, and thus 16 regressors; multiplicative cardiac and respiratory terms cos(c+r), sin(c+r), cos(c-r), sin(c-r), each modelled using an order of two, and thus again 16 regressors; plus respiration volume per time (RVT). In addition to physiological regressors, we constructed 24 motion regressors from the six motion regressors provided (the six original regressors, their derivatives, and the square of the resulting twelve regressors).

### ROI selection for resting-state analyses

We analysed resting-state connectivity between BLA and several *a priori* regions of interest (ROIs) which were informed by anatomical work using tracers in macaque monkeys, focusing on regions with mono-or disynaptic connectivity with BLA, as well as our own work looking at connectivity of the amygdala in humans in relation to mental health dimensions(Klein-Flügge et al., 2022). ROIs are illustrated in **Fig 3a** and ROI selection was performed prior to and independent of subsequent analyses focusing on TUS-induced changes.

BLA is strongly connected to medial and orbital prefrontal cortex (PFC). The strength of this connectivity gradually changes along an anterior-posterior axis, with strongest connectivity in more posterior parts that are closer to the corpus callosum(Aggleton et al., 1980; Amaral et al., 1992). Therefore, we included three posterior/agranular regions in the medial and orbital prefrontal cortex for which homologues in macaques have strong monosynaptic connectivity with the BLA. This included subgenual anterior cingulate cortex (**sgACC**: combined areas 14m and 25 from (Neubert et al., 2014)), pregenual anterior cingulate cortex (**pgACC**: area a24 from (Glasser et al., 2016)) and posterior orbitofrontal cortex (**pOFC**: area 13 from (Neubert et al., 2014)). In addition, BLA receives strong inputs from anterior temporal cortex/temporal pole. We thus included combined areas TGd + TGv from (Glasser et al., 2016) as **TG**. We also included two areas in the insula, anterior dorsal insula (**adInsula**: combined AVI + FOP5 + AAIC + MI from (Glasser et al., 2016)) and posterior/medial insula (**pmInsula**: combined FOP2 + FOP3 + PoI2 from (Glasser et al., 2016)). In addition to these six cortical regions, we also included nine subcortical regions. Here, our focus was, first, on the adjacent hippocampus, for which we included the head, body and tail from (Tian et al., 2020): **HPC_head, HPC_body, HPC_tail**. Second, we included the ventral tegmental area (**VTA**) and nucleus accumbens (**NAc**), both part of the dopaminergic system, taken from the AAN atlas(Edlow et al., 2012) and Harvard Subcortical Atlas, respectively. We also included the bed nucleus of the stria terminalis (**BNST;** mask obtained from (Neudorfer et al., 2020)) sometimes considered part of the extended amygdala, and the hypothalamus, to which amygdala projects via BNST (**hypo**: also from (Neudorfer et al., 2020)). Finally, we included the habenula (**Hb**) and thalamus, taken from the AAN atlas(Edlow et al., 2012) and from the Harvard-Oxford subcortical atlas, respectively. All masks will be shared upon publication and are shown in **Fig 3a**. In summary, we included a total of nine subcortical and six cortical regions.

### fMRI resting-state analyses

For analyses of group and individual resting-state connectivity fingerprints shown in **Fig3**, we first computed the average time course from each ROI, with masks back-projected into subject space. Thus, while a smaller ROI might have a less reliable average time course compared to a larger ROI, all ROIs were given equal weight in subsequent analyses. The functional connectivity between BLA and all other ROIs was computed as the Fisher’s *z*-transformed correlation coefficient. Since all selected ROIs had an average positive connectivity with the BLA seed ROI (averaged across all stimulation sessions), we initially computed the average connectivity of the BLA across the network as the mean across ROIs and both rs-fMRI runs and ran a paired *t*-test to compare this average between BLA and sham stimulation sessions. To explore differences between runs and include the active control stimulation session, we also fit mixed-effects linear regression models using the fitlme function in MATLAB with the factors ‘tus’ (model 1: BLA, sham; model 2: BLA, sham, mIns) and ‘run’ (run1, run2), including their interaction and all random slopes.

### MRS analyses

We first converted the MRS data from the vendor’s .dat (TWIX) format to the NIfTI-MRS format using the spec2nii tool v.0.8.6(Clarke et al., 2022). We then performed preprocessing and model fitting use FSL-MRS v.2.4.4(Clarke et al., 2021). Preprocessing steps included coil combination, windowed average phase- and frequency alignment between repeats, eddy current correction(Klose, 1990), truncation of the FID to remove two time-domain points before the echo centre and truncation of the total length to 1,024 points, removal of the residual water peak using Hankel Lanczos singular value decomposition (HLSVD) over 4.4–4.9 ppm, and finally phase and frequency alignment between averaged edit-on and edit-off spectra. For data sets with spectra in which initial water peak removal failed, we extended the removal range to 4.20–5.65. For data sets in which the initial phase correction failed, we used dynamic alignment in the range of 2.5–3.5 ppm. We also generated a phase-corrected, non-water-suppressed reference from the integrated reference scans. See **Table S9** for the MRSinMRS reporting checklist(Lin et al., 2021).

For each session of each participant, we fit spectral models to both the edit-off spectrum (i.e., the control editing condition) and the difference spectrum between the edit-off and edit-on spectrum. We used a linear combination model in which basis spectra were fitted to the complex-valued spectrum in the frequency domain by scaling, shifting, and broadening. A sequence-specific basis set was simulated using FSL-MRS. Macromolecule basis spectra were added to each basis set. For the edit-off case, four Voigt line-shape peaks were added at 0.8, 1.2, 1.3, and 1.6 ppm. For the difference spectrum, a single basis spectrum comprising eight Lorentzian peaks were added as parameterised and described in Table 1 of(Davies-Jenkins et al., 2024). The 3.0 ppm peak was allowed to vary in concentration relative to the other macromolecular peaks. The macromolecular basis peaks were scaled such that the 3 ppm grouping had an amplitude equivalent to two protons. When fitting the edit-off spectrum, the following metabolites were included in the basis set: γ-aminobutyricacid (GABA), glutamine (Gln), glutamate (Glu), glutathione (GSH), N-acetylaspartate (NAA), N-acetylaspartylglutamate (NAAG) and macromolecules (MM). The OFF spectrum comprised, in addition, ascorbic acid (Asc), aspartic acid (Asp), creatine (Cr), glycerophosphocholine (GPC), myo-inositol (Ins), phosphocholine (PCh), phosphocreatine (PCr), phosphorylethanolamine (PE), scyllo-Inositol (Scyllo), taurine (Tau), and the four MM peaks.

For fitting, we used a Monte-Carlo Markov chain (MCMC) optimiser based on the Metropolis-Hasting algorithm using 5,000 iterations. A low-order polynomial baseline (2^nd^-order for OFF, 0^th^-order for DIFF) was used in the fitting. Across all analyses, we used estimates internally re-referenced to the sum of the NAA and NAAG peaks (results were qualitatively identical when quantifying estimates as molarity or molality concentrations). We corrected estimates for relaxation and tissue type fraction (GM, WM, CSF) in the MRS voxel based on a segmentation of the T1-weighted scan using FSL’s FAST tool. To ensure data quality, we visually inspected all spectra for spurious echo artefacts inside the fitting range (0.2 to 4.2 ppm), excluding those with artefact present. We checked for and removed outlier cases (values more than 3 times the inter-quartile range above the median) on the linewidth based on the full-width at half-maximum (FWHM) of the estimated NAA concentration.

For the amygdala voxel, we had data available from n = 20 participants in whom the BLA had been successfully stimulated, and n = 17 in whom the simulated beam (thresholded at 2 W/cm^2^ I_PA_) overlapped with at least 200 mm^3^/∼5% with the MRS voxel. We excluded two of these participants due to being outliers on the NAA FWHM, and five more for the presence of spurious echo artefacts. Our final analysis contained data sets from n = 11 participants (one additional participant with poor behavioural performance not included in the original n = 20 fulfilled all quality checks for MRS) and thus 22 sessions (BLA-TUS vs sham).

For the mIns voxel, we had data available from n = 18 participants in whom the simulated beam (thresholded at 2 W/cm^2^ I_PA_) overlapped with at least 5% (200 mm^3^) with the MRS voxel (nine participants of n = 29 original participants had insufficient overlap as shown in **Fig 6d**; in one participant, no MRS data was collected in the sham session; in another participant, the overlap could not be established because, due to a technical error, the transducer location during sonication was not recorded, which made it impossible to run acoustic simulations post-hoc). We excluded one more participant for being an outlier on the NAA FWHM of the NAA concentration. As above, we added one additional participant with poor behavioural performance not included in the original n = 29 but who fulfilled all quality checks for MRS, leading to a total of n = 18 participants with n = 36 MRS measurements across the two sessions (mIns-TUS + sham).

We used the GABA+MM concentration estimate from fitting the difference spectrum, given that the difference spectrum allows for a more confident modelling of GABA concentration. We estimated glutamate concentration from the fit of the edit-off spectrum given the glutamate peak in the difference spectrum is closer to the water peak and associated artifacts(van Veenendaal et al., 2018). Results were qualitatively identical when using both metrics from the difference spectrum. The 3 ppm GABA resonance of the difference spectrum contains significant contribution from co-edited macromolecules, typically denoted as “GABA+”. In this work, when we refer to GABA concentration ([GABA]), we use the combined concentration of GABA plus the 3.0 ppm MM peak(Mikkelsen et al., 2018). We quantified E/I balance as the ratio of both metrics [Glu]/[GABA]. To test for an effect of TUS on E/I balance, we performed mixed-effects linear regression with the metabolite estimate as dependent variable and condition (BLA/mIns-TUS vs. sham) as independent variable of interest. We included a random intercept per participant. For control analyses, we added age, gender, time of day at data acquisition, and the overlap between the simulated pressure beam (thresholded at > 2 W/cm^2^ I_PA_) and the MRS voxel as covariates of no interest(Yaakub et al., 2023). For ease of comparison, we report Cohen’s *d* as a metric of effect size. As additional non-parametric control analyses, we performed Wilcoxon signed-ranked tests using the MASS package in R.

## Data availability

Data will be made available upon publication.

## Code availability statement

Code will be made available upon publication.

## Acknowledgements

MCKF was funded by a Wellcome Trust Henry Dale Fellowship (223263/Z/21/Z), a UKRI-converted ERC Starting Grant (EP/X021815/1) and a Leverhulme Award in Psychology. MFSR was funded by an BBSRC grant (BB/W003392/1) and a Wellcome Investigator Award (WT100973AIA, 225924/Z/22/Z). EM is funded by a UKRI Future Leaders Fellowship (MR/T019166/1). WTC is funded by a Wellcome fellowship (225924/Z/22/Z). MKW is funded by an MRC CDA (MR/Y010477/1). EF is funded by a UKRI FLF (MR/Y034368/1), a BBSRC (BB/Y001494/1), a Neuromod+ grant (EP/W035057/1) and an ARIA grant (SCNI-PR01-P15). The funders had no role in study design, data collection and analysis, decision to publish or preparation of the manuscript. The study was also supported by the NIHR Oxford Health Biomedical Research Centre (NIHR203316). The views expressed are those of the author(s) and not necessarily those of the NIHR or the Department of Health and Social Care. The Centre for Integrative Neuroimaging was supported by core funding from the Wellcome Trust (203139/Z/16/Z and 203139/A/16/Z). We thank the UK BBSRC (grant number BB/W019582/1) for support. We thank Rong Bi, Eleonora Carpino, Maja Friedemann, James Hong, Naomi Kingston, Deng Peng, Pranav Sankhe for assistance with data collection. We thank Xinghao Cheng for help with setting up the water tank measurements for validating the calibration of the ultrasound transducer. We thank Ioana Grigoras and Charlotte Stagg for advice with blinding and neuronavigation, Urs Schüffelgen for practical help with gelpad moulding, and the radiographers and Aaron Hess at OxCIN for their help in setting up the scan sequences. This research was funded in whole, or in part, by the Wellcome Trust (203139/Z/16/Z, 203139/A/16/Z, 223263/Z/21/Z, 221794/Z/20/Z, 225924/Z/22/Z and WT100973AIA). For the purpose of open access, the author has applied a CC BY public copyright licence to any Author Accepted Manuscript version arising from this submission.

## Competing interests statement

Bradley E. Treeby and Tim Den Boer report a relationship with NeuroHarmonics LTD that includes: board membership, employment, and equity or stocks. Elsa Fouragnan reports a relationship with Attune Neurosciences that includes: board membership and consulting or advisory. Eleanor Martin reports a relationship with Brainbox that includes: consulting or advisory.

## Author contributions

JA, MR, LW and MCKF conceived the study, designed the task, collected and analysed the data, and wrote the manuscript. TDB helped with data collection. EM and BT advised on ultrasound planning. MG and RC helped with water tank measurements. WTC advised on MRS sequences and data analyses. MW, EF and MFSR gave advice on methodology and data analysis. JA and MCKF wrote the first draft of the manuscript, and all authors edited the manuscript.

