## Supplementary Files for "Low-intensity focused ultrasound to human amygdala reveals a causal role in ambiguous emotion processing and alters local and network-level activity"

Supplementary Materials

Supplementary Figures and Figure Legends
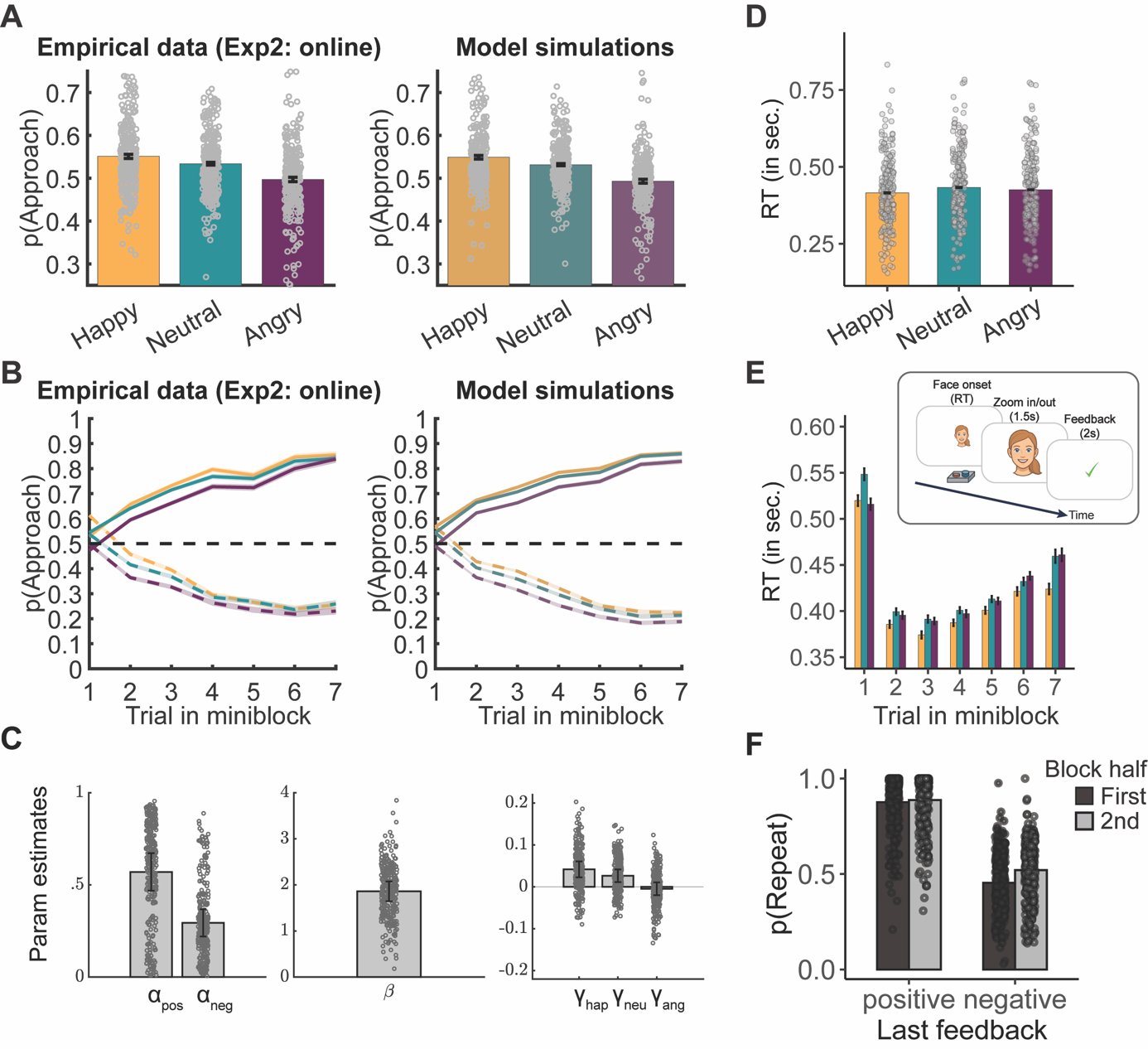

**Figure S1. Regression results for online participants (Experiment 2)**. The key results obtained in the in-person data (n = 29 x 3 sessions) were all replicated in the online data (n = 210). **A, Left:** Participants showed an emotional bias in their responding. Due to the fully balanced design, the expected p(approach) is 50% for each emotion. However, participants were more likely to approach happy over neutral and neutral over angry faces. **Right**: A reinforcement learning (RL) model with separate learning rates for positive and negative feedback and separate approach biases for each emotion could mimic participants’ behaviour. **B, Left**: Over trials in a mini-block, participants correctly learned when to approach (solid) or avoid (dashed) the faces (happy = yellow; neutral = turquoise; angry = purple). **Right**: Again, the model was able to capture this behaviour. **C,** Parameter estimates from model M4 showed faster learning from positive than negative feedback (left) and decreasing approach biases from happy to neutral and neutral to angry faces (right). **D,** Reaction times (RTs) mirrored the emotional bias seen in choices in B, with faster RTs in happy than neutral and angry trials. **E,** Reaction times were fastest on trials 2–4 and slowed down with the likelihood of a new mini-block starting (highest on trial 1 and increasing in likelihood from trial 5 to 7). Different from the in-person data, participants were selectively slowed for neutral faces compared to happy and angry faces on trial #1 of a new block. The difference in RTs between neutral and angry faces disappeared after trial #1. **F.** Participants were more likely to repeat their response following positive compared to negative feedback and late compared to early in the mini-block, showing that feedback was taken into account appropriately.

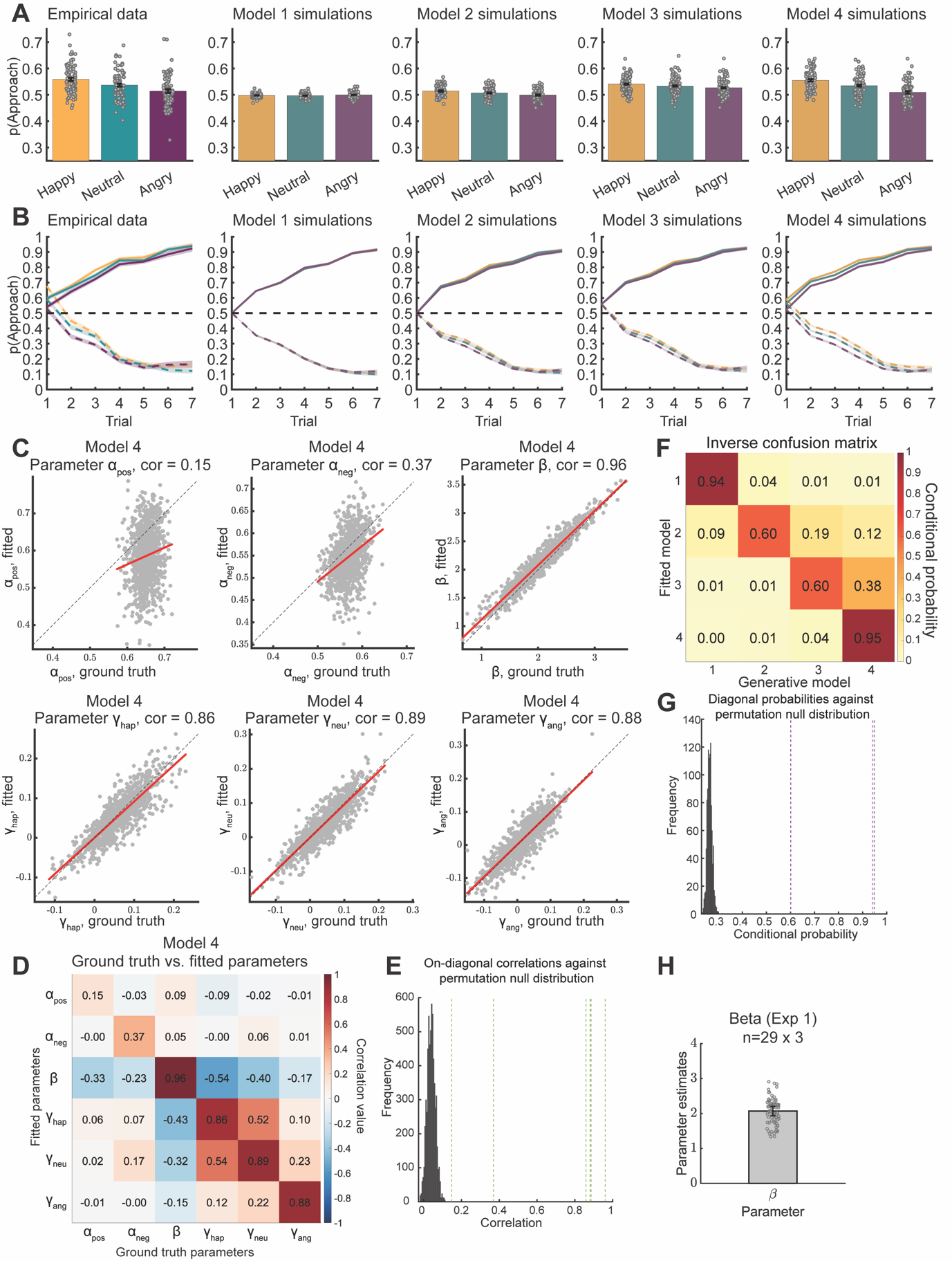
**Figure S2. Qualitative and quantitative validation of computational reinforcement learning models**. **A**, Approach/avoid choices from the empirical in-person data (n = 29 x 3 sessions; leftmost column, as in Fig1B) compared side by side with simulated from increasingly complex computational reinforcement learning models (M1–M4) as a function of the facial emotion displayed (happy = yellow; neutral = turquoise; angry = purple) averaged over all trials in a mini-block. Only M4 was able to qualitatively reproduce the pattern observed in the empirical data. **B,** Approach/avoid choices per facial emotion over trials in a mini-block. Participants correctly learned when to approach (solid) or avoid (dashed) the faces, which was captured by all models. However, only M4, featuring separate approach biases for all three emotions, was able to qualitatively reproduce the emotional biases (i.e., higher approach propensity towards happy than neutral and towards neutral than angry faces) observed in the empirical data. **C,** Parameter recovery for model M4. We fitted a multivariate normal distribution to the empirical parameter values from the in-person data (n = 87 sessions), sampled 1,000 new parameter value combinations, simulated new choices for these parameter values, and refitted M4 to the simulated data (simple Laplace approximation separately per data set, no hierarchical fit). Scatterplots depict the correlation between ground-truth parameter values used for simulating data with the respective fitted parameter values. **D,** Correlations between all ground-truth and fitted parameter values. On-diagonal correlations of ground-truth parameters with their fitted counterparts are generally higher than off-diagonal correlations. There are some negative trade-offs between the inverse temperature parameter β and the learning rates α and the bias parameters γ. **E**, Comparison of the on-diagonal correlations from D (vertical dashed lines) against a permutation null-distribution of correlation values obtained by 1,000 times permuting the assignment of fitted to ground-truth parameter combinations and saving the highest on-diagonal correlation. The 95^th^ percentile of this distribution was r = 0.076, which implies that all empirically observed on-diagonal correlations were significantly higher than expectable under chance. **F,** Model recovery validates that the best fitting model M4 could be reliably distinguished from simpler models. For each of the four models under consideration, we fitted a multivariate normal distribution to the empirical parameter values from the in-person data (n = 87 sessions), sampled 1,000 new parameter value combinations (parameter constraints: α > 0.01, β < 400, |γ| > 0.02), simulated new choices for these parameter values, and refitted each of the four models to each simulated choice data set (simple Laplace approximation separately per data set, no hierarchical fit). We then sorted data sets according to the ground-truth model used for simulating it (x-axis) and the best-fitting model (according to log-model evidence; y-axis) and divided by the row sums. We display the inverse confusion matrix with the conditional probabilities of how likely a data set best fitted by model X (x-axis) was in fact generated by model Y (y-axis). For the best-fitting model M4, the chance that M4 was the true underlying model when it was the best-fitting model was 95%, suggesting that we were well able to identify this model. **G**, Comparison of the on-diagonal conditional probabilities from F (vertical dashed lines) against a permutation null-distribution of conditional probabilities obtained by 1,000 times permuting the assignment of fitted log-model evidences to ground-truth parameter combinations (for each generative model separately) and again sorting data sets by ground-truth and best-fitting model identity. The 95^th^ percentile of this distribution was 0.282, which implies that all empirically observed on-diagonal conditional probabilities were significantly higher than expectable under chance. **H**, distribution of values for the inverse temperature parameter β in the in-person data (n = 29 participants, n = 87 sessions).

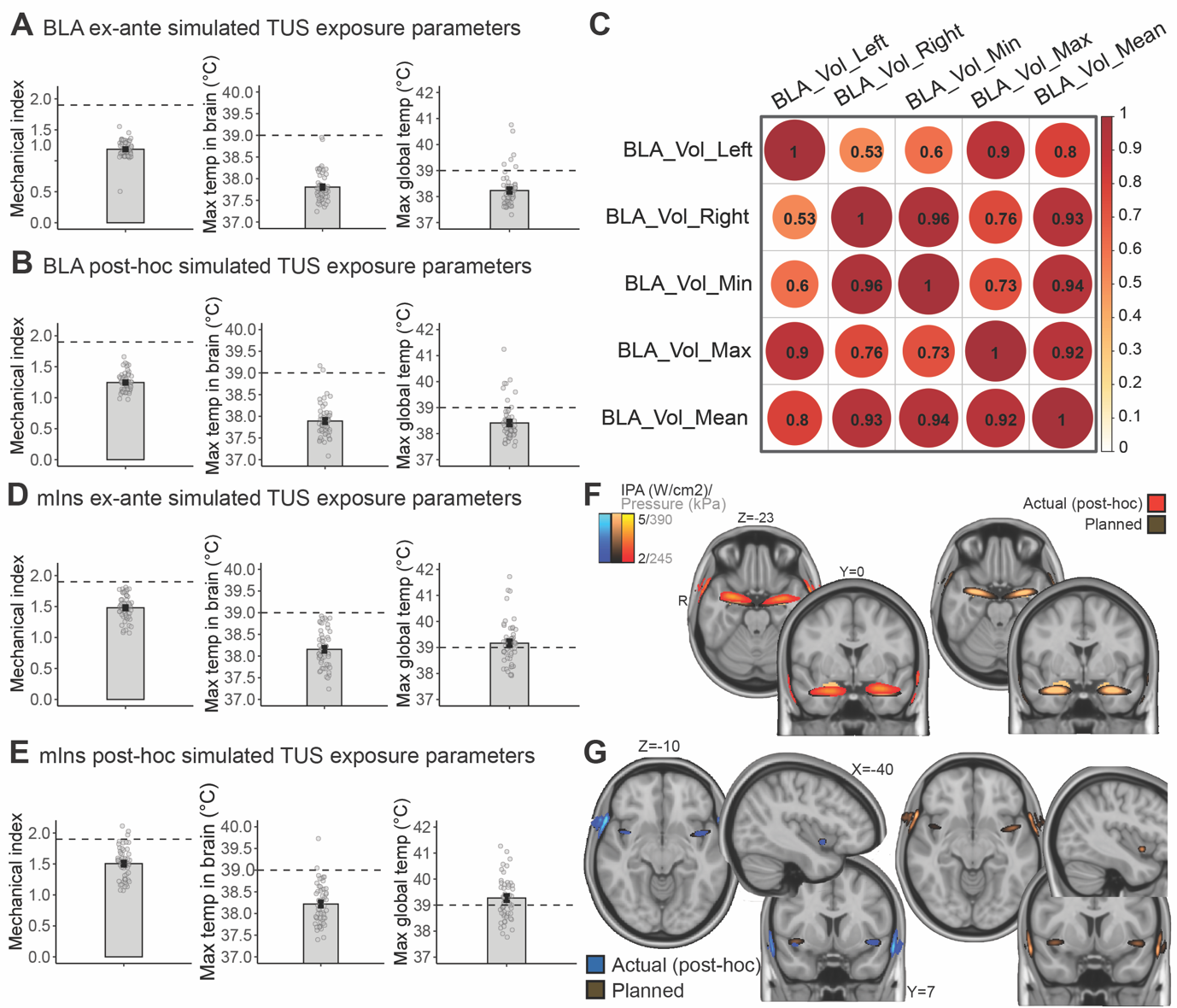

**Figure S3. Safety parameters of ex-ante and post-hoc simulations of the ultrasound pressure field.** **A**, Summary statistics of transcranial mechanical index (MItc), temperature in the brain (the temperature rise can be read off as the values shown minus 37°; to derive values in the brain, we used a mask made up of the conjunction of the gray-matter, white-matter, and CSF masks obtained with SPM) and global temperature rise for *ex-ante* simulations of the BLA stimulations (performed before the BLA-TUS sessions; n = 20 participants, left and right BLA, thus n = 40 data points). The MItc was always < 1.9. in two cases, the temperature rise in the brain exceeded the threshold of 2°C (here shown as >39°); however, note that the thermal dose did not exceed 0.25 CEM43 and was thus within ITRUSST guidelines (see **Tables S3-S6**). **B,** Summary statistics for *post-hoc* simulations of the BLA stimulations (based on coordinates recorded with neuronavigation during the BLA-TUS sessions). The MItc was always < 1.9 and the temperature rise in the brain always < 2°C. **C**, Intercorrelation between possible covariates for stimulation volume (for n = 20 participants), including the stimulated tissue in the left and right BLA (quantified as the volume in mm^3^ with I_PA_ values > 2 W/cm^2^ within an anatomical BLA mask) as well the minimum of either (used in the main results), maximum of both, and mean of both. **D,** Summary statistics for *ex-ante* simulations of the mIns stimulations (n = 29 participants, coordinate for one session missing, thus n = 28 included; left and right mIns, thus n = 56 data points). The MItc was always < 1.9 and the temperature rise in the brain always < 2°C. **E**, summary statistics for *post-hoc* simulations of the mIns stimulations. The MItc exceeded the threshold of 1.9 in three cases. Temperature rise in the brain exceeded the threshold of 2°C in two cases; however, as for BLA-TUS, note that again the thermal dose did not exceed 0.25 CEM43 in these cases and was thus within ITRUSST guidelines. Please refer to **tables S3-S6** for a conservative estimate of the thermal dose. **F,G:** comparison of ex ante and post-hoc (planned vs actual) simulations for BLA (F, top) and mIns (G, bottom), in red vs brown or blue vs brown, respectively. Left shows both planned and actual overlaid on top of each other like in Fig 2; right shows just planned locations.

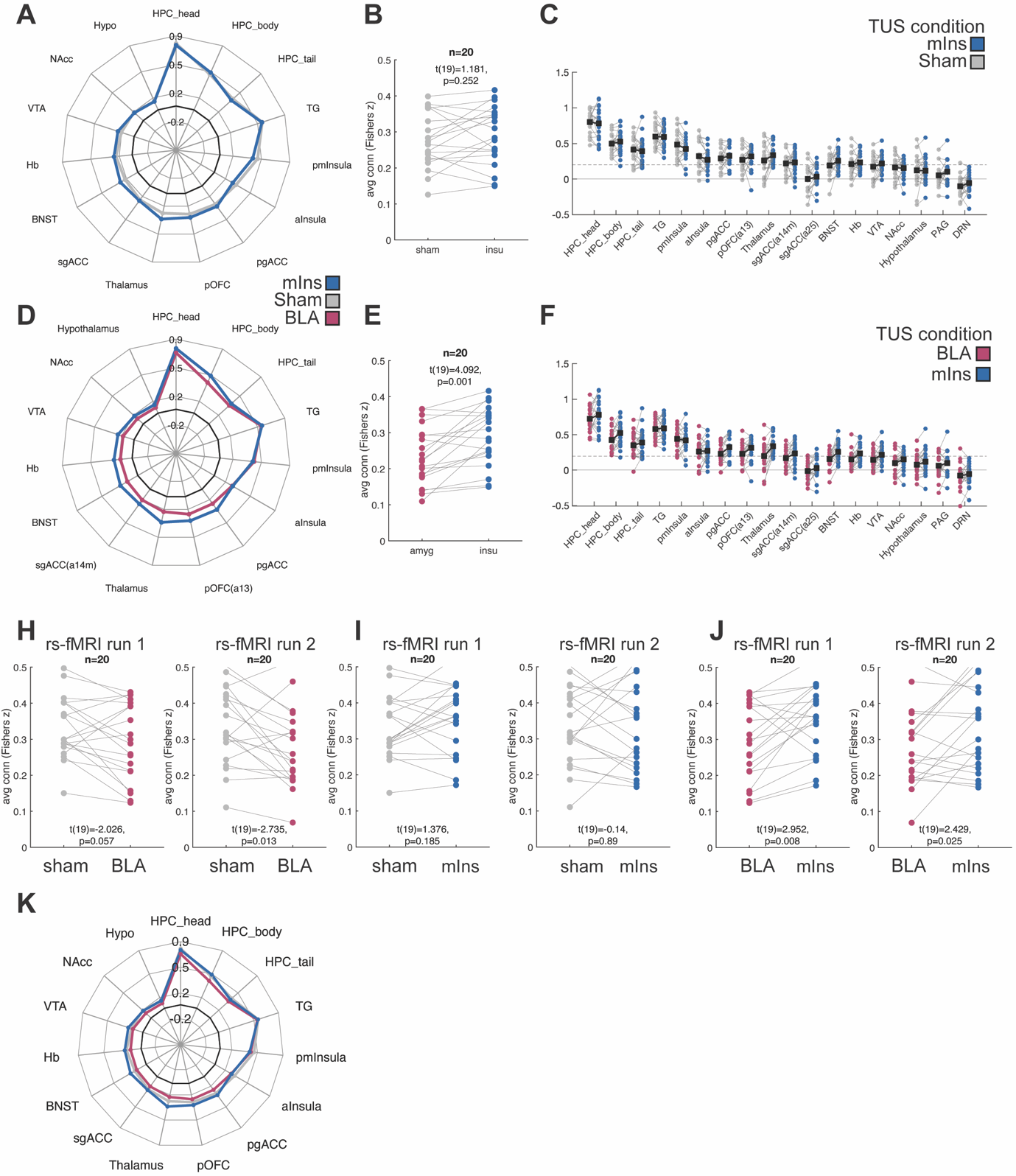

**Figure S4, BLA resting-state connectivity for all TUS conditions and split across runs. A-C**, Following mIns TUS, there was no change in the BLA’s resting-state connectivity fingerprint (A), the average connectivity across the fingerprint (B), or consistent changes across connections as seen in individual data. points (C). **D-F**, However, when comparing mIns and BLA TUS sessions directly, BLA connectivity was reduced following BLA TUS compared to mIns TUS as seen in the fingerprint (D), average connectivity across all regions in the fingerprint (E) and when including individual datapoints (F). **H-J** show comparisons between all three TUS conditions: BLA vs sham (H), mIns vs sham (I) and BLA vs mIns (J) for run 1 (left) and run2 (right), respectively. This showed no difference between runs and similar trends in both run 1 and 2. **K**, Overlay of the BLA fingerprint for all three TUS session types. The change was specific to BLA stimulation.

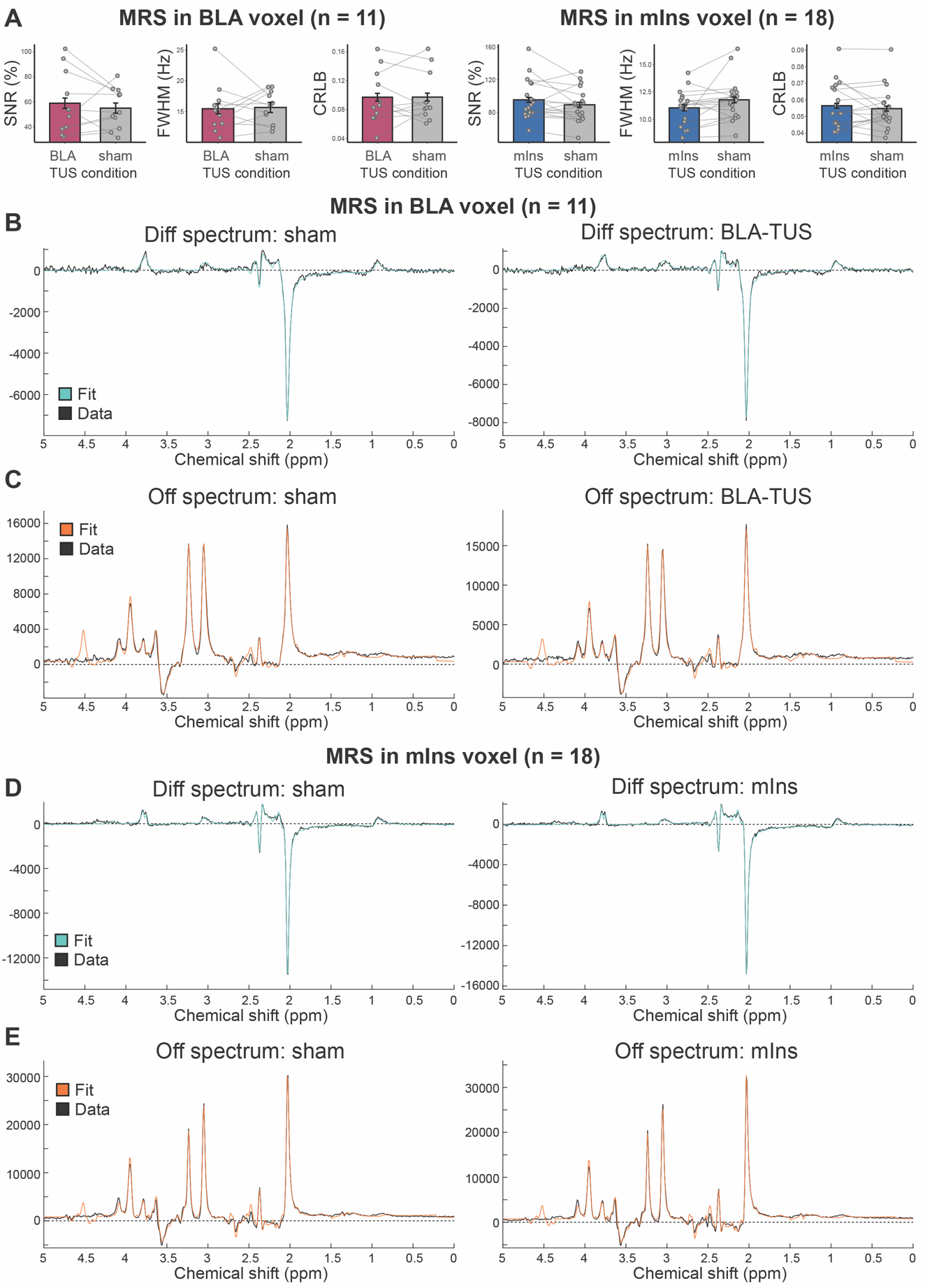

**Figure S5, MRS quality controls and MRS group spectra. A**, MRS quality control metrics show signal-to-noise ratio (SNR), full-width half-maximum (FWHM) and Cramér-Rao Lower Bound (CRLB) for both MRS voxels, the BLA voxel (left) and the mIns voxel (right), and for both active (pink/blue) and sham (grey) conditions. **B-E**, Group mean spectra (black) and fits are shown for all MRS data. Data for the BLA voxel are shown in B-C and for the mIns voxel in D-E; data from sham sessions are displayed on the left and the active TUS sessions on the right. B+D show the diff spectra used to estimate GABA (fit: turquoise) and C+E show off spectra used to estimate glutamate (fit: orange). Note that the range considered during fitting corresponds to x axis limits of (0.2, 4.2) ppm. GABA differences quantified and reported as part of the key results in the manuscript can be appreciated visually at e.g. ~3ppm in the Diff spectra.

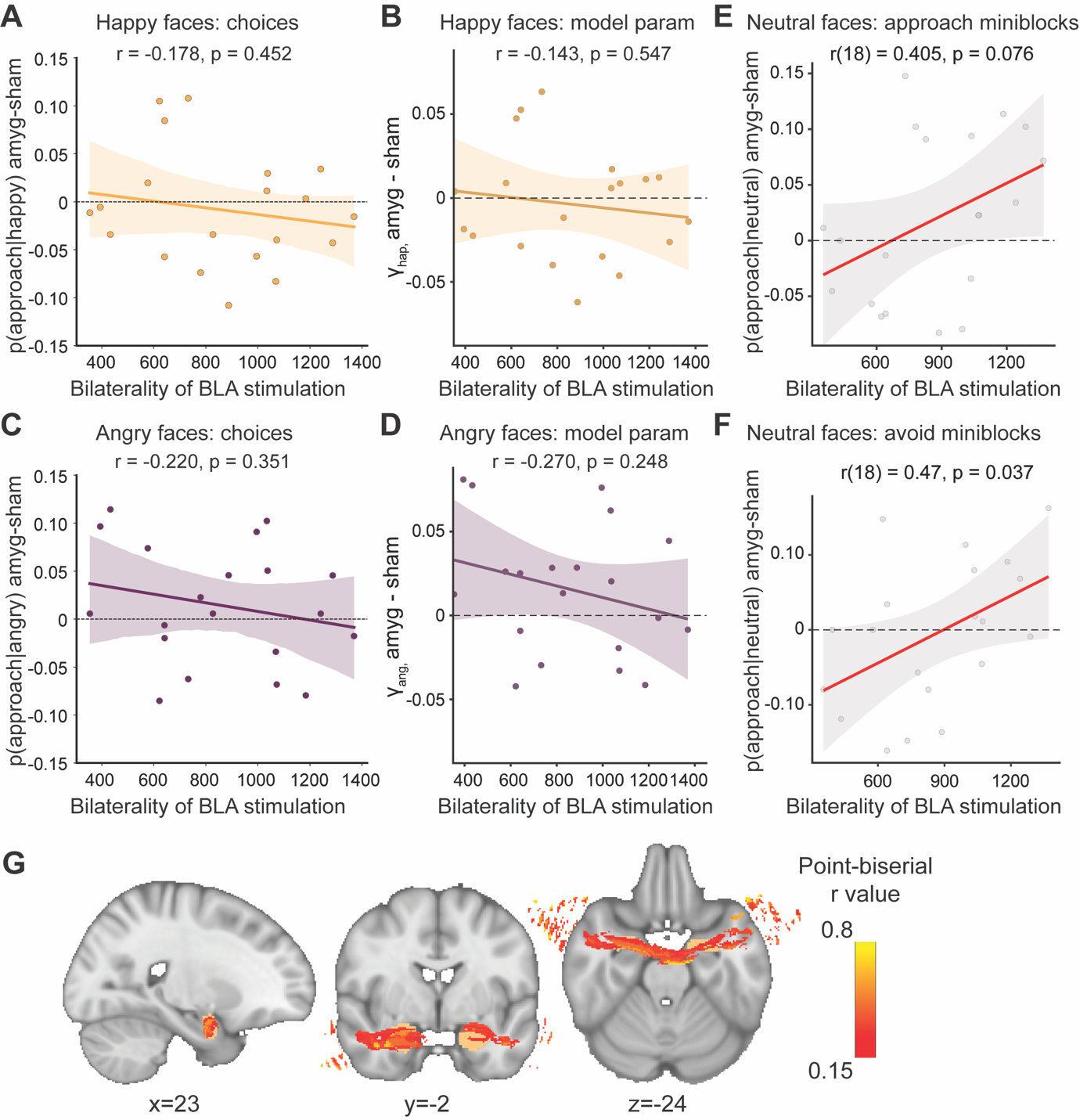

**Figure S6. Control analyses for BLA-TUS effects on behaviour.** **A,** The bilaterality of BLA stimulation (volume > 2 W/cm^2^ in the less stimulated hemisphere) did not significantly correlate with any chance in the propensity to approach happy facial stimuli after BLA-TUS. **B,** There was also no significant correlation between the bilaterality of BLA stimulation and the model-derived parameter reflecting the approach bias to happy faces. **C,** The bilaterality of BLA stimulation (volume > 2 W/cm^2^ in the less stimulated hemisphere) did not significantly correlate with any chance in the propensity to approach angry facial stimuli after BLA-TUS. **D,** There was also no significant correlation between the bilaterality of BLA stimulation and the model-derived parameter reflecting the approach bias to angry faces. The increase in the probability of approaching of neutral faces following BLA-TUS was present both in miniblocks where participants were required to approach (**E**) as well as avoid (**F**) neutral faces. **G,** All simulated pressure fields for BLA post-hoc simulations were brought into MNI standard space and thresholded at 2 W/cm^2^, assigning values of 1 to voxel with values above this threshold and values of 0 to all other voxels. We then computed the point-biserial correlation between the voxel-specific TUS stimulation and the behavioral effect size p(approach|neutral & BLA-TUS) – p(approach|neutral & sham) across participants (n = 20), akin to voxel-based lesion-symptom mapping approaches. There were positive correlations between the change in approach behaviour to neutral faces and stimulation of both the left and right BLA, but these correlations were higher and more extended for the right (compared to the left) BLA.

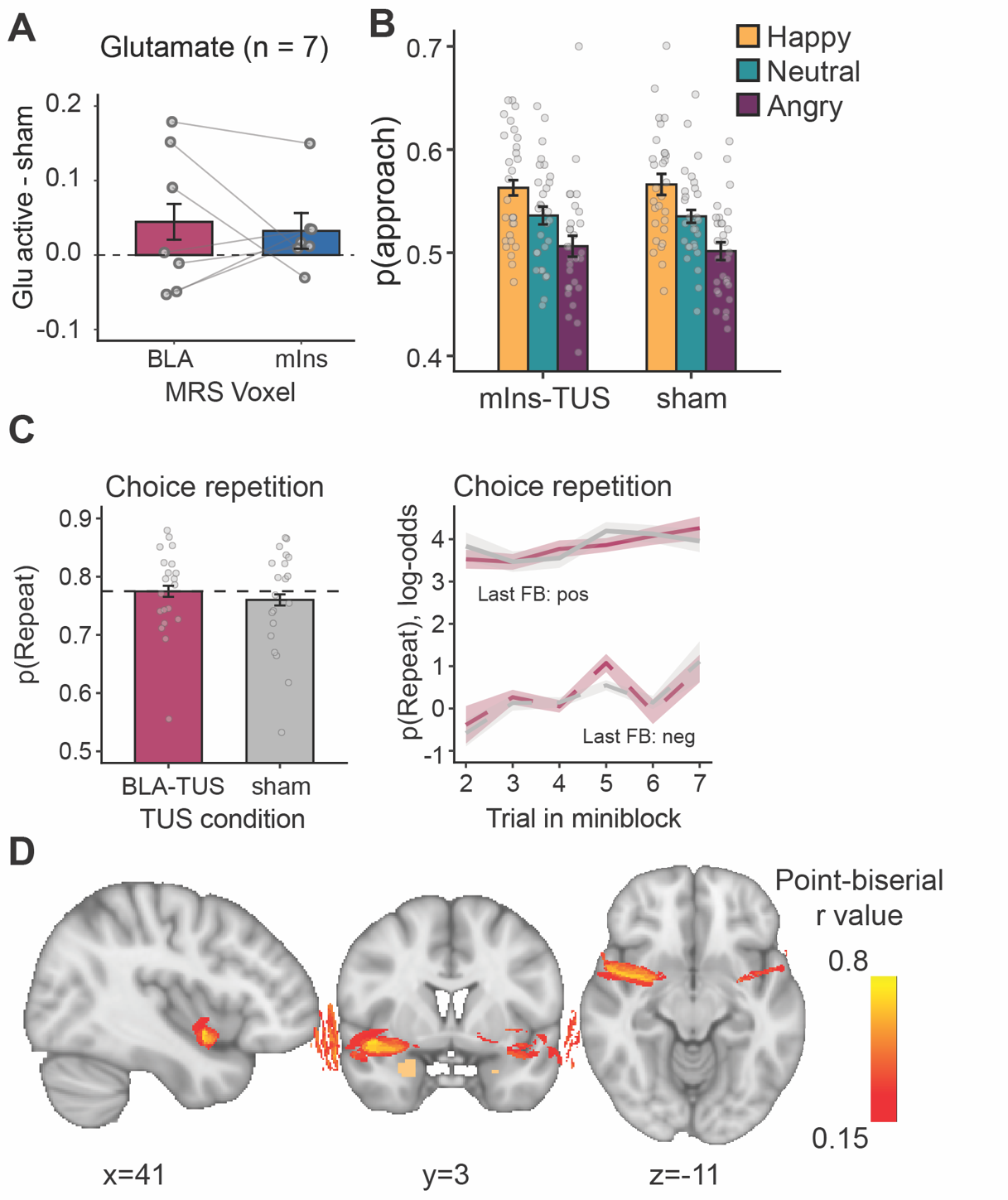

**Figure S7. Control analyses for mIns-TUS.** **A,** Following BLA-TUS compared to mIns-TUS, when comparing metabolite changes in the BLA and mIns voxel, there was no significant change in glutamate concentration, suggesting the same TUS protocol had opposing effects on GABA in the two regions, but not on glutamate (n = 7, 21 sessions). **B,** The proportion of trials when participants approached happy (yellow), neutral (turquoise), or angry (purple) faces was not significantly changed following mIns-TUS (n = 29). **C**, There was no significant change in choice repetitions after BLA-TUS. Hence, this behavioural effect was specifically seen only after mIns TUS, both on average across all trials (left) and when illustrating the probability of repeating the previous response following negative (dashed) and positive (solid) feedback after BLA-TUS (red lines) and sham (grey lines) for each trial position in a mini-block (right). **D,** All simulated pressure fields for mIns post-hoc simulations were brought into MNI standard space and thresholded at 2 W/cm^2^, assigning values of 1 to voxel with values above this threshold and values of 0 to all other voxels. We then computed the point-biserial correlation between the voxel-specific and the behavioral effect size p(repeat|mIns-TUS) – p(repeat|sham) across participants (n = 29). There were positive correlations between the change in choice repetitions and stimulation of bilateral ventral mid-insula. These correlations were higher and more extended for the right (compared to the left) mid-insula.

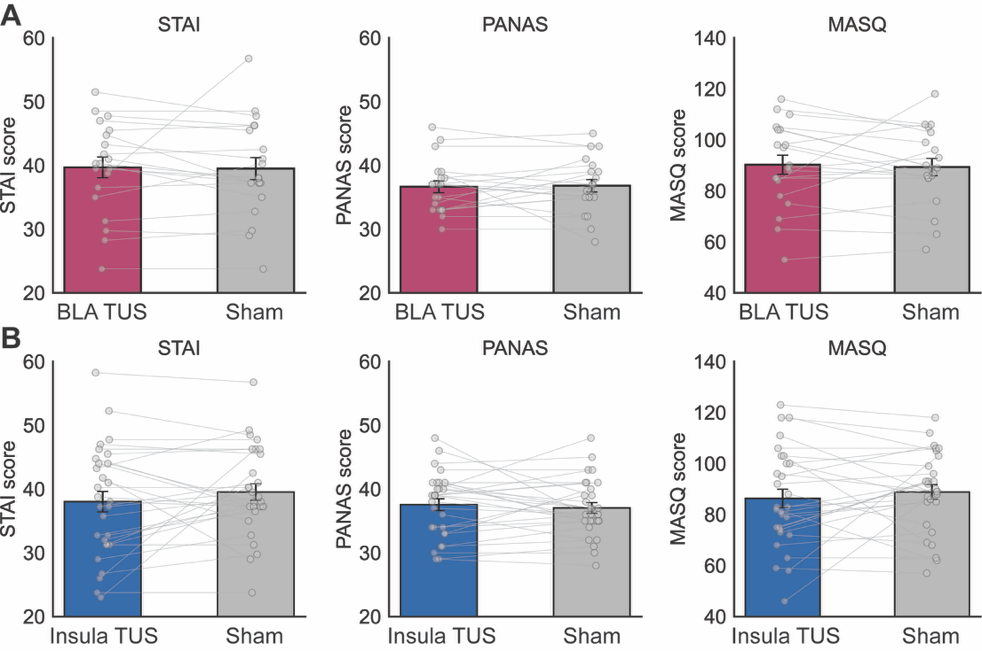

**Figure S8. Self-reported mood**, Participants rated 43 items selected from the STAI-State (20 items), PANAS (10 items) and MASQ (37 items) on a 5-point scale (1 = Not at all, 2 = Slightly, 3 = Moderately, 4 = Quite a bit, 5 = Extremely). Shown are self-report measures (STAI, subsets of PANAS and MASQ) following BLA-TUS (**A**) and mIns-TUS (**B**) relative to sham. As expected for a healthy cohort and given the brevity of the TUS intervention, there was no change in any scale or for any TUS condition.

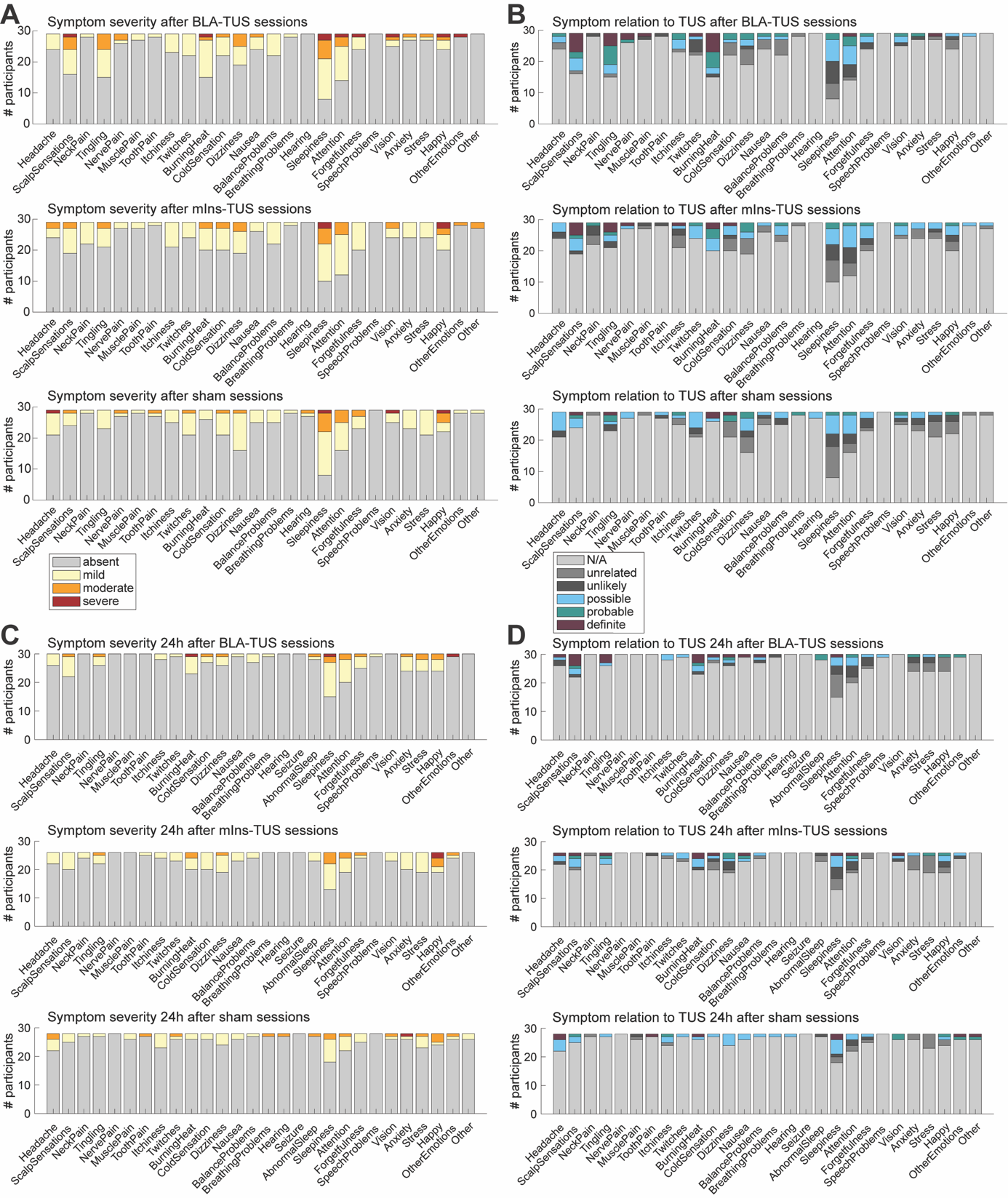

**Figure S9. Self-reported side-effects after BLA-TUS, mIns-TUS, and sham sessions.** At the end of each MRI-TUS session, as well as 24-hours post-TUS, participants were asked to rate the severity (1 = Absent, 2 = Mild, 3 = Moderate, 4 = Severe) and the perceived relation to TUS (0 = N/A, 1 = Unrelated, 2 = Unlikely, 3 = Possible, 4 = Probable, 5 = Definite) of 26 and 28 symptoms, respectively. The 24-hour post TUS survey had two extra questions related to the sleep quality and occurrence of seizures. **A,** Post TUS session severity of reported side effects for each symptom following BLA-TUS (upper panel), mIns-TUS (middle panel) and sham (lower panel). **B,** Post TUS session perceived relation to TUS for each symptom. **C,** 24-hour post TUS severity of reported side effects for each symptom following BLA-TUS (upper panel), mIns-TUS (middle panel) and sham (lower panel). **D,** 24-hour post TUS perceived relation to TUS for each symptom.

### Supplementary Methods

#### Standardized reporting for transcranial ultrasound

| **Transducer and Drive System Description** | |
| --- | --- |
| Transducer manufacturer and model number | NeuroFUS PRO CTX-212-4CH S/N: 021 transducer; Sonic Concepts; Brainbox Ltd |
| Transducer centre frequency | 212 kHz |
| Active Diameter | 64.0 mm |
| Geometric Focus | 63.2 mm also referred to as the Radius of Curvature (ROC). |
| Focal Depth | 52.4 mm measured from exit plane of transducer housing rim to geometric focus. |
| # of Elements | 1 central disc and 3 annular rings |
| Drive system components, including manufacturer and model number | NeuroFUS TPO-105-19; Sonic Concepts; Brainbox Ltd |
| Matching Network | 4 each – MR safe fundamental mode RF impedance matching networks are supplied inside a single RF shielded external enclosure |
| **Drive System Settings** | |
| Operating frequency | 212 kHz |
| Output level settings | Free field I_SPPA_: mean = 30.5 W/cm^2^, range 15.2-48 W/cm^2^ |
| Focal position settings | Amygdala TUS: 45.7-58.4 mm  Insula TUS: 32.4-40 mm |
| Description of transducer coupling method | Moulded gelpad (Aquaflex, Parker Laboratories Inc.), moulded to be 10.8mm at the transducer centre, exactly filling the transducer cavity, but to become thinner towards the edge of the transducer and not extend beyond the transducer edge (i.e., adding 0mm between participant’s head and transducer). |

**Table S1: Methodological details describing our transducer and drive system.**

**Free Field Acoustic Parameters**

The TPO maintains a constant I_SPPA_ along the steering range 32.4 – 58.4 mm

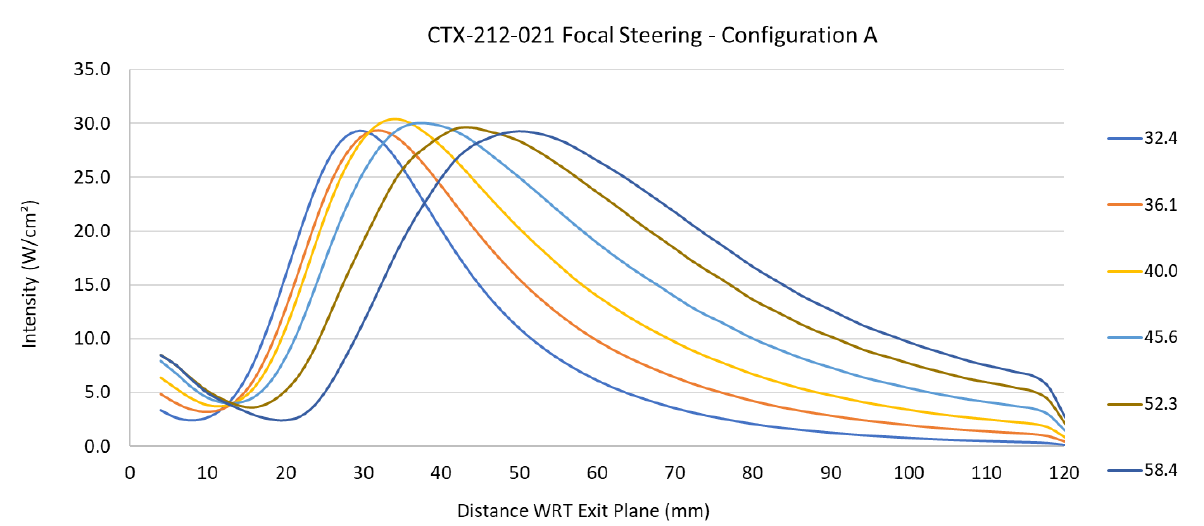

□ Reference position for measurements

- Transducer exit plane

□ Spatial-peak pressure amplitude

- Free-field value used during calibration: 950kPa
- Free-field values used in experiment: Mean 956 kPa, range: 675 – 1200 kPa.

□ Position of centre of -6dB focal region (relative to transducer exit plane)

- 32.4-58.4 mm

□ Size of focal volume (-3 dB and -6 dB axial lengths and lateral widths) and position of centre of focal volume (centre of -3 dB relative to reference position)

| Position (mm) | Axial -6dB | Axial -3dB |
| --- | --- | --- |
| 32.4 | 25.7 | 17.2 |
| 36.1 | 30.2 | 19.8 |
| 40.0 | 35.5 | 23.1 |
| 45.6 | 43.6 | 28.8 |
| 52.3 | 50.1 | 33.6 |
| 58.4 | 52.6 | 35.6 |

| Position (mm) | Lateral -6dB | Lateral -3dB |
| --- | --- | --- |
| 40 | 8 | 5.9 |
| 45 | 8.6 | 6.3 |
| 50 | 9.3 | 6.7 |
| 55 | 10 | 7.3 |

□ Description of how free field parameters were obtained

- Onda HNR-0500 calibrated hydrophone (SN: 2196) and calibrated oscilloscope, in an automated scanning tank filled with degassed and deionised water.

**Pulse Timing Parameters**

|  | Duration | Ramp Duration | Ramp Shape | Repetition Interval/ Frequency |
| --- | --- | --- | --- | --- |
| Pulse | 20 ms | None | Rectangular | 200 ms/ 5 Hz |
| Pulse Train | 80 s | None | Rectangular |  |

**Table S2: Pulse timing parameters used in all active sessions (mIns and BLA).**

**In Situ Estimates of Exposure Parameters estimated using acoustic simulations in k-Plan (see Methods)**

|  | **Left BLA** | | | | | **Right BLA** | | | | |
| --- | --- | --- | --- | --- | --- | --- | --- | --- | --- | --- |
| **ID** | **Free-field I_sppa_**  **(W/cm2)** | **Focal Depth (mm)** | **Target I_sppa_ (W/cm2)** | **MI_tc_** | **CEM43** | **Free-field I_sppa_**  **(W/cm2)** | **Focal Depth (mm)** | **Target I_sppa_ (W/cm2)** | **MI_tc_** | **CEM43** |
| 1 | 30 | 48 | 1.59 | 0.51 | 0.0008 | 30 | 46 | 6.44 | 1.01 | 0.0013 |
| 2 | 30 | 49 | 7.62 | 1.24 | 0.0019 | 25 | 52 | 8.2 | 1.15 | 0.0018 |
| 3 | 26 | 50 | 7.23 | 1.13 | 0.0019 | 25 | 52 | 8.48 | 1.15 | 0.0018 |
| 4 | 27 | 47 | 8.56 | 1.17 | 0.0013 | 27 | 48 | 8.02 | 1.08 | 0.0009 |
| 5 | 30 | 52 | 8.55 | 1.33 | 0.0021 | 25 | 50 | 7.91 | 1.08 | 0.0009 |
| 6 | 27.5 | 51 | 8.24 | 1.17 | 0.0011 | 23 | 51 | 7.98 | 1.18 | 0.001 |
| 7 | 30 | 52 | 7.37 | 1.14 | 0.0025 | 27 | 51 | 7.47 | 1.31 | 0.0011 |
| 8 | 25 | 55 | 7.97 | 1.33 | 0.0013 | 20 | 52 | 7.77 | 1.11 | 0.0008 |
| 9 | 26.5 | 53 | 8.16 | 1.32 | 0.0022 | 31 | 53 | 7.64 | 1.29 | 0.0035 |
| 10 | 30 | 52 | 8.13 | 1.35 | 0.0019 | 25 | 52 | 7.81 | 1.22 | 0.0005 |
| 11 | 25 | 51 | 7.71 | 1.13 | 0.001 | 25 | 50 | 7.95 | 1.32 | 0.0013 |
| 12 | 30 | 48 | 8 | 1.24 | 0.0014 | 29 | 50 | 7.65 | 1.18 | 0.002 |
| 13 | 35 | 52 | 7.83 | 1.45 | 0.0048 | 28.5 | 52 | 7.36 | 1.29 | 0.0023 |
| 14 | 38 | 49 | 7.69 | 1.26 | 0.012 | 32 | 52 | 7.91 | 1.2 | 0.0073 |
| 15 | 23.5 | 45.7 | 8.04 | 1.12 | 0.0012 | 23.5 | 47.1 | 7.21 | 1.08 | 0.0012 |
| 16 | 226 | 57 | 7.59 | 1.35 | 0.0596 | 25.5 | 53 | 8.07 | 1.19 | 0.0034 |
| 17 | 15.2 | 54 | 7.89 | 1.07 | 0.0008 | 20 | 53 | 7.88 | 1.12 | 0.001 |
| 18 | 35 | 56.3 | 7.69 | 1.55 | 0.0094 | 25.5 | 54.9 | 7.29 | 1.36 | 0.0426 |
| 19 | 19.5 | 56 | 8.06 | 1.07 | 0.0007 | 17.5 | 53 | 8.11 | 1.08 | 0.0008 |
| 20 | 21.5 | 52.5 | 7.66 | 1.15 | 0.0012 | 25 | 53.3 | 7.52 | 1.07 | 0.0016 |
| 21 | 32 | 55.1 | 8.11 | 1.29 | 0.0024 | 31 | 49.9 | 8.2 | 1.34 | 0.0016 |
| 22 | 28.5 | 53.8 | 7.46 | 1.27 | 0.0024 | 24.7 | 58.4 | 8.05 | 1.29 | 0.0015 |
| 23 | 24.5 | 54.8 | 7.97 | 1.22 | 0.0009 | 28.5 | 53.5 | 8.06 | 1.33 | 0.0014 |
| 24 | 21.5 | 55.5 | 7.5 | 1.05 | 0.0009 | 28 | 56.7 | 7.58 | 1.29 | 0.0023 |
| 25 | 19 | 54 | 8.01 | 1.07 | 0.0009 | 16.5 | 52 | 8.18 | 1.08 | 0.0008 |
| 26 | 23.4 | 57.6 | 8.02 | 1.08 | 0.0064 | 25 | 57.5 | 8.08 | 1.16 | 0.0096 |
| 27 | 19.2 | 58.4 | 7.98 | 1.06 | 0.0007 | 19.2 | 58.4 | 8.05 | 1.11 | 0.0008 |
| 28 | 25.7 | 51 | 8.02 | 1.07 | 0.0015 | 21.5 | 49 | 8.04 | 1.07 | 0.0012 |
| 29 | 28.5 | 58 | 9.12 | 1.14 | 0.0021 | 30.6 | 58 | 8.54 | 1.15 | 0.0028 |

**Table S3:** For each participant (ID column), and the left and right hemisphere, from left to right columns, we report the free-field I_SPPA_ (TPO output), and *ex ante* simulation-derived estimated BLA target I_SPPA_, global maximum MI_tc_ and CEM43. Note that the CEM43 index reported here is a conservative proxy that assumes the global maximum temperature constantly for 80 seconds, so it is an overestimate of the true CEM43.

|  | **Left mIns** | | | | | **Right mIns** | | | | |
| --- | --- | --- | --- | --- | --- | --- | --- | --- | --- | --- |
| **ID** | **Free-field I_sppa_**  **(W/cm2)** | **Focal Depth (mm)** | **Target I_sppa_ (W/cm2)** | **MI_tc_** | **CEM43** | **Free-field I_sppa_**  **(W/cm2)** | **Focal Depth (mm)** | **Target I_sppa_ (W/cm2)** | **MI_tc_** | **CEM43** |
| 1 | 26 | 37 | 5.16 | 1.04 | 0.0036 | 26 | 37 | 5.35 | 1.16 | 0.0018 |
| 2 | 35 | 38 | 6.31 | 1.34 | 0.0214 | 32 | 40 | 8.26 | 1.08 | 0.0034 |
| 3 | 40 | 33 | 5.46 | 1.52 | 0.0075 | 32 | 34 | 6.15 | 1.53 | 0.0104 |
| 4 | 35 | 32 | 7.77 | 1.61 | 0.0018 | 35 | 32 | 7.07 | 1.55 | 0.0016 |
| 5 | 35 | 36 | 7.81 | 1.42 | 0.0049 | 38 | 37 | 8.05 | 1.19 | 0.0083 |
| 6 | 38 | 32.4 | 7.79 | 1.34 | 0.0019 | 27.5 | 33 | 7.59 | 1.3 | 0.0012 |
| 7 | 37.5 | 37 | 6.68 | 1.39 | 0.0088 | 37.5 | 37 | 6.63 | 1.68 | 0.0029 |
| 8 | 27 | 38 | 8.15 | 1.07 | 0.0012 | 25 | 37 | 8.08 | 1.2 | 0.0012 |
| 9 | 30 | 34 | 6.34 | 1.39 | 0.0144 | 37 | 33 | 6.71 | 1.43 | 0.0077 |
| 10 | 32 | 36 | 5.07 | 1.5 | 0.0049 | 40 | 34 | 7.84 | 1.49 | 0.0049 |
| 11 | 38 | 32 | 6.81 | 1.3 | 0.0041 | 38 | 32 | 6.61 | 1.52 | 0.0054 |
| 12 | 38.5 | 32 | 7.96 | 1.33 | 0.0029 | 38 | 32 | 7.41 | 1.45 | 0.0056 |
| 13 | 35 | 34 | 5.07 | 1.48 | 0.0177 | 40 | 35 | 6.78 | 1.42 | 0.0073 |
| 14 | 30 | 34 | 4.66 | 1.56 | 0.0169 | 37 | 33 | 5.75 | 1.73 | 0.0136 |
| 15 | 40 | 34.1 | 8.09 | 1.66 | 0.0084 | 30 | 33.9 | 6.51 | 1.55 | 0.0053 |
| 16 | 30 | 34 | 5.09 | 1.55 | 0.1101 | 40 | 33 | 6.45 | 1.76 | 0.0068 |
| 17 | 28.5 | 33 | 8.05 | 1.36 | 0.0026 | 29.5 | 33 | 7.94 | 1.55 | 0.0039 |
| 18 | 40 | 32.8 | 5.68 | 1.79 | 0.1052 | 25 | 34.1 | 3.61 | 1.72 | 0.2266 |
| 19 | 42 | 33.9 | 8.17 | 1.61 | 0.0041 | 43 | 33.8 | 7.07 | 1.56 | 0.0065 |
| 20 | 30 | 32.5 | 6.06 | 1.27 | 0.0042 | 28 | 33.1 | 5.17 | 1.44 | 0.024 |
| 21 | 38 | 33 | 7.43 | 1.46 | 0.0042 | 43 | 32.6 | 5.67 | 1.61 | 0.0146 |
| 22 | 42 | 34.1 | 8.08 | 1.54 | 0.0151 | 42 | 33 | 6.96 | 1.78 | 0.0182 |
| 23 | 31.5 | 33.4 | 7.75 | 1.17 | 0.0019 | 33 | 33.9 | 6.23 | 1.42 | 0.0021 |
| 24 | 38.5 | 33.1 | 6.88 | 1.79 | 0.0042 | 33 | 33 | 5.72 | 1.75 | 0.0061 |
| 25 | 31 | 33 | 7.27 | 1.12 | 0.0073 | 28.5 | 33 | 7.14 | 1.26 | 0.0016 |
| 26 | 38.5 | 33.3 | 7.49 | 1.81 | 0.0074 | 32 | 33.4 | 7.03 | 1.68 | 0.0709 |
| 27 | 28.6 | 34.3 | 7.99 | 1.11 | 0.0014 | 34.4 | 33.2 | 8.01 | 1.37 | 0.0038 |
| 28 | 46 | 32.4 | 7.92 | 1.39 | 0.0091 | 41 | 33 | 7.6 | 1.33 | 0.0071 |
| 29 | 42 | 32.4 | 7.65 | 1.74 | 0.0083 | 48 | 33 | 7.77 | 1.77 | 0.0066 |

**Table S4:** For each participant (ID column), and the left and right hemisphere, from left to right columns, we report the free-field I_SPPA_ (TPO output), and *ex ante* simulation-derived estimated mIns target I_SPPA_, global maximum MI_tc_ and CEM43. Note that the CEM43 index reported here is a conservative proxy that assumes the global maximum temperature constantly for 80 seconds, so it is an overestimate of the true CEM43.

**Post-hoc simulation of exposure parameters**

|  | **Left BLA** | | | | | **Right BLA** | | | | | **Both** |
| --- | --- | --- | --- | --- | --- | --- | --- | --- | --- | --- | --- |
| **ID** | **Free-field I_sppa_**  **(W/cm2)** | **Target I_sppa_ (W/cm2)** | **MI_tc_** | **CEM43** | **Vol (mm3)** | **Free-field I_sppa_**  **(W/cm2)** | **Target I_sppa_ (W/cm2)** | **MI_tc_** | **CEM43** | **Vol (mm3)** | **Min Vol** |
| 1 | 30 | 3.17 | 1.26 | 0.012 | 229.92 | 30 | 5.12 | 1.23 | 0.002 | 525.23 | 229.92 |
| 2 | 30 | 6.81 | 1.37 | 0.0041 | 1037.39 | 25 | 8.98 | 1.17 | 0.0027 | 1355.91 | 1037.39 |
| 3 | 26 | 4.93 | 1.19 | 0.0012 | 388.12 | 25 | 1.96 | 1.14 | 0.0017 | 179.72 | 179.72 |
| 4 | 27 | 0.81 | 1.34 | 0.0017 | 4.64 | 27 | 1.08 | 1.11 | 0.0017 | 108 | 4.64 |
| 5 | 30 | 1.18 | 1.32 | 0.0042 | 38.81 | 25 | 5.12 | 1.14 | 0.0013 | 530.72 | 38.81 |
| 6 | 27.5 | 7.17 | 1.14 | 0.0014 | 932.77 | 23 | 1.39 | 1.53 | 0.0017 | 132.47 | 132.47 |
| 7 | 30 | 7.66 | 1.25 | 0.0148 | 1213.31 | 27 | 6.74 | 1.55 | 0.002 | 1069.03 | 1069.03 |
| 8 | 25 | 4.7 | 1.39 | 0.0013 | 728.16 | 20 | 4.81 | 1.17 | 0.0011 | 621.84 | 621.84 |
| 9 | 26.5 | 3.19 | 1.26 | 0.0026 | 288.98 | 31 | 6.96 | 1.35 | 0.0027 | 855.98 | 288.98 |
| 10 | 30 | 7.29 | 1.66 | 0.0053 | 1188.84 | 25 | 3.85 | 1.34 | 0.0018 | 433.27 | 433.27 |
| 11 | 25 | 3.27 | 1.51 | 0.002 | 427.78 | 25 | 3.96 | 1.21 | 0.0016 | 394.45 | 394.45 |
| 12 | 30 | 7.35 | 1.22 | 0.0015 | 1039.08 | 29 | 5.53 | 1.39 | 0.0031 | 641.25 | 641.25 |
| 13 | 35 | 7.42 | 1.42 | 0.0007 | 827.3 | 28.5 | 7.72 | 1.27 | 0.0028 | 1084.22 | 827.3 |
| 14 | 38 | 7.59 | 1.31 | 0.0187 | 1085.06 | 32 | 2.92 | 1.37 | 0.019 | 354.38 | 354.38 |
| 15 | 23.5 | 7.74 | 1.14 | 0.0015 | 959.34 | 23.5 | 6.93 | 1.08 | 0.0014 | 732.38 | 732.38 |
| 16 | NA | NA | NA | NA | NA | NA | NA | NA | NA | NA | NA |
| 17 | 15.2 | 7.34 | 1.08 | 0.0009 | 903.23 | 20 | 5.67 | 1.1 | 0.0011 | 577.12 | 577.12 |
| 18 | 35 | 1.58 | 1.56 | 0.0228 | 121.08 | 25.5 | 3.22 | 1.53 | 0.1171 | 410.06 | 121.08 |
| 19 | 19.5 | 8.02 | 1.14 | 0.001 | 1034.44 | 17.5 | 7.57 | 1.12 | 0.0008 | 1301.91 | 1034.44 |
| 20 | 21.5 | 7.99 | 1.07 | 0.001 | 1287.56 | 25 | 7.75 | 1.09 | 0.0015 | 1334.39 | 1287.56 |
| 21 | 32 | 7.31 | 1.18 | 0.0045 | 837.84 | 31 | 6.41 | 1.06 | 0.0013 | 641.67 | 641.67 |
| 22 | 28.5 | 6.55 | 1.26 | 0.0023 | 1131.05 | 24.7 | 7.86 | 1.33 | 0.0015 | 1072.41 | 1072.41 |
| 23 | 24.5 | 7.31 | 1.45 | 0.0016 | 1218.38 | 28.5 | 7.25 | 1.25 | 0.0015 | 887.62 | 887.62 |
| 24 | 21.5 | 7.36 | 1.15 | 0.0012 | 1197.7 | 28 | 8.08 | 1.18 | 0.0016 | 1184.2 | 1184.2 |
| 25 | 19 | 6.41 | 0.97 | 0.0012 | 995.2 | 16.5 | 6.83 | 0.98 | 0.0009 | 1047.94 | 995.2 |
| 26 | 23.4 | 7.92 | 1.13 | 0.0077 | 1344.94 | 25 | 6.7 | 1.09 | 0.0032 | 780.05 | 780.05 |
| 27 | 19.2 | 8.46 | 1.09 | 0.0008 | 1369.41 | 19.2 | 8.47 | 1.14 | 0.0009 | 1657.97 | 1369.41 |
| 28 | 25.7 | 7.72 | 1.25 | 0.0017 | 1046.67 | 21.5 | 0.69 | 1.3 | 0.0021 | 60.33 | 60.33 |
| 29 | 28.5 | 8.31 | 1.08 | 0.0035 | 1241.58 | 30.6 | 9.29 | 1.29 | 0.0026 | 1440.7 | 1241.58 |

**Table S5:** For each participant (ID column), and the left and right hemisphere, from left to right columns, we report the free-field I_SPPA_ (TPO output, same as in Tables S3/S4), and *post-hoc* simulation-derived estimated target BLA I_SPPA_, global maximum MI_tc_, CEM43 and BLA volume. Note that the CEM43 index reported here is a conservative proxy that assumes the global maximum temperature constantly for 80 seconds, so it is an overestimate of the true CEM43. For s16, accidentally, no brainsight coordinates were recorded; hence, post-hoc simulations were not possible (note this participant is excluded from all amygdala results).

|  | **Left mIns** | | | | | **Right mIns** | | | | |
| --- | --- | --- | --- | --- | --- | --- | --- | --- | --- | --- |
| **ID** | **Free-field I_sppa_**  **(W/cm2)** | **Target I_sppa_ (W/cm2)** | **MI_tc_** | **CEM43** | **Vol (mm3)** | **Free-field I_sppa_**  **(W/cm2)** | **Target I_sppa_ (W/cm2)** | **MI_tc_** | **CEM43** | **Vol (mm3)** |
| 1 | 26 | 4.91 | 1.13 | 0.0045 | 486 | 26 | 5.11 | 1.21 | 0.0037 | 475.45 |
| 2 | 35 | 5.39 | 1.7 | 0.015 | 506.67 | 32 | 5.84 | 1.16 | 0.0052 | 51.05 |
| 3 | 40 | 4.9 | 1.59 | 0.0134 | 533.25 | 32 | 6.26 | 1.51 | 0.0058 | 456.89 |
| 4 | 35 | 7.56 | 1.66 | 0.002 | 605.81 | 35 | 6.66 | 1.74 | 0.003 | 536.2 |
| 5 | 35 | 7.21 | 1.55 | 0.0034 | 467.44 | 38 | 8.41 | 1.25 | 0.0084 | 727.73 |
| 6 | 38 | 6.47 | 1.57 | 0.0048 | 718.45 | 27.5 | 7.44 | 1.33 | 0.0017 | 420.19 |
| 7 | 37.5 | 7.76 | 1.48 | 0.0459 | 708.33 | 37.5 | 6.85 | 1.66 | 0.0029 | 447.19 |
| 8 | 27 | 6.9 | 1.07 | 0.0011 | 336.23 | 25 | 7.19 | 1.57 | 0.0021 | 393.19 |
| 9 | 30 | 6.47 | 1.53 | 0.0608 | 94.08 | 37 | 5.58 | 1.87 | 0.017 | 444.23 |
| 10 | NA | NA | NA | NA | NA | NA | NA | NA | NA | NA |
| 11 | 38 | 5.85 | 1.49 | 0.0024 | 478.41 | 38 | 6.81 | 1.56 | 0.008 | 391.5 |
| 12 | 38.5 | 8.12 | 1.24 | 0.0034 | 434.53 | 38 | 6.9 | 1.72 | 0.0085 | 437.91 |
| 13 | 35 | 4.5 | 1.59 | 0.0123 | 407.11 | 40 | 6.57 | 1.47 | 0.0095 | 751.36 |
| 14 | 30 | 5.27 | 1.17 | 0.005 | 132.89 | 37 | 4.62 | 1.82 | 0.0193 | 387.28 |
| 15 | 40 | 8.54 | 1.31 | 0.0032 | 712.97 | 30 | 6.61 | 1.57 | 0.0041 | 306.28 |
| 16 | 30 | 5.18 | 1.84 | 0.5765 | 543.8 | 40 | 5.06 | 2.11 | 0.0901 | 577.12 |
| 17 | 28.5 | 8.42 | 1.22 | 0.0028 | 527.34 | 29.5 | 7.79 | 1.54 | 0.0057 | 423.56 |
| 18 | 40 | 5.86 | 1.58 | 0.0302 | 502.03 | 25 | 3.23 | 1.59 | 0.1207 | 120.66 |
| 19 | 42 | 8.97 | 1.44 | 0.0044 | 603.7 | 43 | 6.13 | 1.97 | 0.0348 | 646.31 |
| 20 | 30 | 6.4 | 1.21 | 0.0034 | 622.27 | 28 | 4.14 | 1.67 | 0.0171 | 177.61 |
| 21 | 38 | 7.53 | 1.48 | 0.0046 | 707.91 | 43 | 5.19 | 1.47 | 0.0159 | 499.08 |
| 22 | 42 | 8.09 | 1.51 | 0.0133 | 742.5 | 42 | 6.33 | 1.76 | 0.0174 | 561.52 |
| 23 | 31.5 | 7.57 | 1.17 | 0.0015 | 413.86 | 33 | 6.12 | 1.45 | 0.0022 | 502.03 |
| 24 | 38.5 | 6.64 | 1.85 | 0.0076 | 394.03 | 33 | 4.97 | 1.85 | 0.0089 | 385.17 |
| 25 | 31 | 6.77 | 1.44 | 0.0423 | 274.22 | 28.5 | 6.35 | 1.13 | 0.0014 | 635.77 |
| 26 | 37.5 | 7.66 | 1.56 | 0.0049 | 677.11 | 32 | 4.84 | 1.26 | 0.0128 | 510.47 |
| 27 | 28.6 | 8.2 | 1.08 | 0.0009 | 511.31 | 34.4 | 8.34 | 1.21 | 0.0025 | 790.17 |
| 28 | 46 | 7.8 | 1.36 | 0.0077 | 688.08 | 41 | 7.82 | 1.25 | 0.0059 | 749.67 |
| 29 | 42 | 7.64 | 1.75 | 0.0085 | 664.88 | 48 | 7.34 | 2.03 | 0.0152 | 677.53 |

**Table S6:** For each participant (ID column), and the left and right hemisphere, from left to right columns, we report the free-field I_SPPA_ (TPO output, same as in Tables S3/S4), and *post-hoc* simulation-derived estimated target mIns I_SPPA_, global maximum MI_tc_, CEM43 and mIns volume. Note that the CEM43 index reported here is a conservative proxy that assumes the global maximum temperature constantly for 80 seconds, so it is an overestimate of the true CEM43. For s10, accidentally, no brainsight coordinates were recorded; hence, post-hoc simulations were not possible.

#### Average TUS coordinates in MNI space

##### Ex-ante coordinates

| Target | Side | Coordinate | X | Y | Z |
| --- | --- | --- | --- | --- | --- |
| BLA | Left | Transducer | -86 | 14 | -24 |
| (x = 20) |  | Skin | -75 | 11 | -23 |
|  |  | Target | -21 | -3 | -23 |
|  | Right | Transducer | 88 | 15 | -25 |
|  |  | Skin | 77 | 12 | -24 |
|  |  | Target | 23 | -3 | -23 |
| mIns | Left | Transducer | -87 | 15 | -10 |
| (x = 29) |  | Skin | -76 | 12 | -10 |
|  |  | Target | -41 | 5 | -7 |
|  | Right | Transducer | 88 | 19 | -11 |
|  |  | Skin | 77 | 16 | -10 |
|  |  | Target | 43 | 7 | -7 |
| Average planned (ex-ante) coordinates of transducer (back of emitting surface), skin (front of emitting surface), and target in MNI space for both BLA and mIns sonications. | | | | | |

**Table S7:** The average BLA and mIns planned (*ex ante*) coordinates of the transducer, skin and target are reported for both hemispheres in MNI coordinates.

##### Post-hoc coordinates

| Target | Side | Coordinate | X | Y | Z |
| --- | --- | --- | --- | --- | --- |
| BLA | Left | Transducer | -87 | 13 | -24 |
| (n = 20) |  | Skin | -76 | 11 | -23 |
|  |  | Target | -22 | -2 | -20 |
|  | Right | Transducer | 89 | 15 | -25 |
|  |  | Skin | 78 | 13 | -24 |
|  |  | Target | 24 | 0 | -23 |
| mIns | Left | Transducer | -87 | 15 | -10 |
| (n = 29) |  | Skin | -76 | 12 | -9 |
|  |  | Target | -41 | 5 | -8 |
|  | Right | Transducer | 89 | 19 | -11 |
|  |  | Skin | 78 | 16 | -11 |
|  |  | Target | 43 | 9 | -10 |
| Average measured (post-hoc) coordinates of transducer (back of emitting surface), skin (front of emitting surface), and target in MNI space for both BLA and mIns sonications. | | | | | |

**Table S8:** The average BLA and mIns planned (*post-hoc*) coordinates of the transducer, skin and target are reported for both hemispheres in MNI coordinates.

#### Participants and Experimental Procedure for Experiment 2 (online)

Ethical approval for this study was obtained through the Medical Sciences Interdivisional Research Ethics Committee (MS-IDREC; ref: R73912/RE001). Informed consent was obtained from each participant before they began the experiment. Behavioural and questionnaire data was collected using Prolific (prolific.co). We collected data from 344 participants. We excluded 130 participants who either showed (a) a mean accuracy < 0.60 during trials 4–7 of the mini-blocks (i.e., when learning should asymptote) in any of the six task conditions or (b) a strong response bias (> 80% of the same response) for any of the three emotion conditions. Afterwards, we excluded another four participants with mean RTs larger than three times the inter-quartile range above the median RT across participants. All analyses include data from the remaining 210 participants (mean age = 30.83, males = 121, females = 88, other = 1).

Participants took part in a single session online. They were given task instructions (for task details, see below), and their task understanding was probed using 5 multiple choice questions. Upon correct completion of these questions, they were admitted to the main part of the study and asked to fill in some questionnaires, play four blocks of the task (M = 41 minutes, SD = 10.5), rate 22 neutral images subjectively on two sliders corresponding to happy and angry on a Likert scale with 4 levels, and fill in some more questionnaires. The total session duration was on average 77 minutes (SD = 24.78).

#### Transcranial Ultrasound Stimulation Safety Screening Form

1. Are you currently diagnosed, or have you ever been diagnosed with any neurological or psychiatric conditions? YES NO

2. Do you have a personal or family history of epilepsy or seizures? YES NO

3. Did you ever suffer from recurring (i.e., more than one) fainting fits or febrile convulsions? YES NO

4. Are you currently taking any prescribed or unprescribed medications (or herbal remedies)? YES NO

5. Are you currently undergoing anti-malarial treatment? YES NO

6. Are you currently taking psychoactive medication or recreational drugs? YES NO

7. Do you suffer from migraines? YES NO

8. Are you currently suffering, or have you ever suffered from alcohol or drug dependency (defined as excessive and regular intake of alcohol or drugs over several weeks which would typically involve a loss of control) YES NO

9. Have you had more than 3 units of alcohol in the last 24 hours? YES NO

10. Are you taking part in other brain stimulation or pharmacological studies within the time scale of this study (i.e., with less than a week gap from any of the visits for this study, or in the time between the first and the last visit of this study)? YES NO

11. Do you have a pacemaker or pacing wires? YES NO

12. Have you had any surgery to your heart, head (including eyes), neck or spine? YES NO

13. Do you have any implanted devices? YES NO

14. Do you have cochlear implants? YES NO

15. Do you have an implanted neurostimulator (e.g. DBS, epidural/subdural, VNS)? YES NO

16. Do you have a medication infusion device? YES NO

17. Do you have metal in the brain, skull or elsewhere in your body (e.g., splinters, fragments, clips, etc.)? YES NO

18. Have you had any surgical procedures or endoscopies in the last 6 weeks? YES NO

19. Have you had any other surgical procedures of any kind? YES NO

20. Have you had any injuries involving metal to the eyes or other part of the body? YES NO

21. Have you had any operations involving metallic pins/plates/screws/wires? YES NO

22. Have you had any serious accident or injury (e.g., involving impact to the head)? YES NO

23. Have you ever experienced a fit or blackout, or suffers from epilepsy or diabetes? YES NO

24. Do you have the following: body piercings, coloured contact lenses, eye makeup (unless removable for MRI scan); hearing aid, wearable medical device; tattoos; dentures, dental braces, dental implants, dental bridge; medicated skin patches; artificial limb, protheses, splint, brace or support, IUD (coil) YES NO

25. Is there any chance that you could be pregnant? YES NO

26. Have you ever had an MRI scan before? YES NO

27. Have you ever participated in neurostimulation studies before? YES NO

28. How many hours sleep did you have last night?

#### Mood questionnaire

We reproduce here the exact questionnaire that participants answered at the end of each TUS visit.

The following list shows 43 words that describe different feelings, sensations, and experiences. Please use the scale below to indicate for each word to what extent it describes how you are feeling right now.

1 = Not at all, 2 = Slightly, 3 = Moderately, 4 = Quite a bit, 5 = Extremely

1 Content

2 Full of energy

3 Tense
4 Sociable

5 Tired

6 Focused

7 Stressed

8 Optimistic

9 Satisfied

10 Cheerful

11 Afraid

12 Confused

13 Sad

14 Alert

15 Nauseous

16 Happy

17 Nervous

18 Irritable

19 Bored

20 Clearheaded

21 Confident

22 Worried

23 Fatigued

24 Hopeful

25 Decisive

26 Light (free of burdens/worries)

27 Relaxed

28 Attentive

29 Excited

30 Interested

31 Jittery

32 Upset

33 Aroused

34 Calm

35 Sleepy

36 Sluggish

37 Secure

38 Strained

39 At Ease

40 Frightened

41 Comfortable

42 Steady

43 Pleasant

#### TUS debrief questionnaire

We reproduce here the debrief questionnaire that participants answered at the end of each TUS visit.

1. Do you think that you received **real or placebo stimulation** today (placebo meaning ineffective stimulation)?

Real/Placebo

2. What did the ultrasound stimulation **feel** like?

3. What did the ultrasound stimulation **sound** like?

4. If you felt or heard anything: did it feel/sound the **same on both sides** of your head? (If no, how did it feel different)

5. Did the ultrasound make you feel different in terms of your **emotions** (e.g., stressed, happy, worried, lighter)? If yes, can you please describe the emotion and how long approximately this change lasted?

6. Did the ultrasound make you feel any different in terms of your **body/sensation** (e.g., tingling, headache, …)? If yes, can you describe how, whether the sensation was local to the stimulation site or elsewhere in your body and how long approximately did this change last?

7. Did you experience anything unpleasant or painful during the study not mentioned above?

YES/NO

Please provide two values per row to indicate if you experienced any of the following:

|  | Value  1 = Absent  2 = Mild  3 = Moderate  4 = Severe | Relation  1= Unrelated  2 = Unlikely  3 = Possible  4 = Probable  5 = Definite |
| --- | --- | --- |
| *Bodily changes/sensations* |  |  |
| Headache |  |  |
| Scalp sensations (i.e., feelings on the skin of your head) |  |  |
| Neck Pain |  |  |
| Tingling |  |  |
| Nerve pain (e.g., down your facial nerves) |  |  |
| Muscle pain |  |  |
| Tooth Pain |  |  |
| Itchiness |  |  |
| Unusual twitches or muscle movements |  |  |
| Burning or heat |  |  |
| Cold sensation |  |  |
| Dizziness |  |  |
| Nausea (stomach sickness) |  |  |
| Balance problems |  |  |
| Breathing problems |  |  |
| Change in hearing |  |  |
| *Cognitive changes* |  |  |
| Sleepiness |  |  |
| Difficulty paying attention |  |  |
| Forgetfulness |  |  |
| Speech problems |  |  |
| Vision problems (e.g., double vision) |  |  |
| *Emotions/feelings* |  |  |
| Anxious/Worried |  |  |
| Stressed |  |  |
| Happy/Feeling lighter |  |  |
| Other emotion/feeling (specify): |  |  |
| *Other* |  |  |
| Other  Please explain: |  |  |

#### MRSinMRS reporting checklist

| 1. Hardware | |
| --- | --- |
| a. Field strength | 7 T |
| b. Manufacturer | Siemens Healthineers, Erlangen, Germany |
| c. Model (software version) | MAGNETOM 7T Plus (VE12U-AP01) |
| d. RF coils: nuclei (transmit/receive), number of channels, type, body part | Nova 8Tx32Rx Head Coil (Nova Medical, Wilmington, MA, USA) |
| e. Additional hardware |  |
| 2. Acquisition | |
| a. Pulse sequence | MEGA-edited semi-LASER sequence from the CMRR Spectroscopy package |
| b. Volume of interest (VOI) locations | Two single voxels (consecutive acquisitions) placed over the basolateral amygdala (BLA) and mid-insula (mIns) |
| c. Nominal VOI size (cm^3^, mm^3^) | 20 x 14 x 20 mm^3^ |
| d. Repetition time (T_R_), echo time (T_E_) [ms, s] | BLA: TR = 4 sec., TE = 78 ms  mIns: TR = 5 sec., TE = 78 ms |
| e. Total number of excitations or acquisitions per spectrum  In time series for kinetic studies  i. Number of average spectra (NA) per time point  ii. Averaging method (eg block-wise or moving average)  iii. Total number of spectra (acquired/in time series) | BLA: 192 averages (96 edit-on and 96 edit-off)  mIns: 128 averages (64 edit-on and 64 edit-off) |
| f. Additional sequence parameters (spectral width in Hz, number of spectral points, frequency offsets)  If STEAM: mixing time (TM) If MRSI: 2D or 3D, FOV in all directions, matrix size, acceleration factors, sampling method | 130 Hz bandwidth editing pulse, centred around 1.9 ppm (edit-on condition) and at 7.5 ppm (edit-off condition) |
| g. Water suppression method | Integrated VAPOR water suppression and OVS (VAPOR bandwidth = 150 Hz) |
| h. Shimming method, reference peak, and thresholds for “acceptance of shim” chosen | One-step (linear 6-projection) FASTESTMAP from the CMRR spectroscopy package (Gruetter and Tkáč, 2000) |
| i. Triggering or motion correction method (respiratory, peripheral, cardiac triggering, incl. device used and delays) | N/A |
| 3. Data analysis methods and outputs | |
| a. Analysis software | Both processing and fitting was done in FSL-MRS version 2.4.4 (Clarke et al., 2021). |
| b. Processing steps deviating from quoted reference or product | Standard pre-processing:   1. coil combination, 2. windowed average phase- and frequency alignment between repeats, 3. eddy current correction (Klose, 1990), 4. truncation of the FID to remove two time-domain points before the echo centre and truncation to 1,024 data points in total 5. removal of the residual water peak using Hankel Lanczos singular value decomposition (HLSVD) over 4.4–4.9 ppm, 6. phase and frequency alignment between averaged edit-on and edit-off spectra   Deviations:   1. For data sets with spectra in which initial water peak removal failed, we extended the removal range to 4.20–5.65 2. For data sets in which the initial phase correction failed, we used dynamic alignment in the range of 2.5–3.5 ppm   Standard fitting:   1. Common to all fits: Fit set of basis sets with MCMC sampling using the Metropolis-Hasting algorithm with 5,000 samples (with priors disabled); reference to internal reference of sum of NAA + NAAG peak; also compute outcome in metrics of molality and molarity using the water reference and the tissue fractions in the voxel based on a segmentation with fsl_anat 2. Specific to difference fit: flat (polynomial of order 0) baseline; basis set for difference fit 3. Specific to edit-off fit: polynomial baseline of order 2; basis set for edit-off fit |
| c. Output measure (eg absolute concentration, institutional units, ratio) | Glutamate and GABA concentration relative to the sum of the NAA + NAAG peaks (internal reference)  E/I balance: ratio of glutamate concentration (from the fit to the edit-off spectrum) and the GABA concentration (from the fit to the difference spectrum) |
| d. Quantification references and assumptions, fitting model assumptions | NAA + NAAG peak as reference for both glutamate and GABA concentrations; we confirmed results also hold when instead expressed in molality and molarity units using the water reference and tissue fractions from the voxel segmentation |
| 4. Data quality | |
| a. Reported variables (SNR, linewidth (with reference peaks)) | See .csv files with SNR, FWHM, and CRLB values for all participants/all sessions/all metabolites in the data sharing collection. |
| b. Data exclusion criteria | BLA voxel:   1. Select n = 20 in whom the BLA had been successfully stimulated with TUS 2. Select n = 17 in whom the simulated pressure beam (thresholded at 2 W/cm2 ISPPA) overlapped with at least 200 mm^3^/~5% with the MRS voxel 3. Select n = 10 clean data sets while excluding 7 outliers: 2 outliers (> 3 times the interquartile range above the median) on the NAA FWHM and 5 outliers due to the presence of spurious echo artefacts. 4. Add n = 1 in whom all MRS quality and overlap criteria hold but who were not part of n = 20 due to poor task performance   mIns voxel:   5. Select N = 29 in whom the mIns has been successfully stimulated with TUS 6. Select n = 18 in whom the simulated pressure beam (thresholded at 2 W/cm2 ISPPA) overlapped with at least 200 mm^3^/5% with the MRS voxel 7. Select n = 17 clean data sets by excluding 1 outlier (> 3 times the interquartile range above the median) on the NAA FWHM. 8. Add n = 1 in whom all MRS quality and overlap criteria hold but who were not part of n = 20 due to poor task performance |
| c. Quality measures of postprocessing model fitting (eg CRLB, goodness of fit, SD of residual) | FWHM of the NAA peak |
| d. Sample spectrum | Fig. 4C; Fig. 6F; Fig S5; see single-session fits in the report.html files of each session in the data sharing collection |

**Table S9:** MRSinMRS reporting checklist of the MRS acquisition parameters and analysis methods(Lin et al., 2021).

### Supplementary Results

#### Behavioural results across all online data, N = 210 (Experiment 2)

Here we report the identical analyses reported for the in-person cohort in the main part of the manuscript for the larger dataset acquired online in the absence of any TUS. First, we analysed participants approach/avoid responses as a function of the required response, the emotion of the displayed face, and mini-block half (trials 1–4 vs. 5–7). Online participants exhibited an emotional bias (Fig S1; main effect of emotion; χ^2^(2) = 80.473, *p* < .001), with overall more approach responses to happy than neutral faces (*b* = 0.050, 95%-CI [0.027 0.074], χ^2^(1) = 17.207, *p* < .001) and more approach responses to neutral than angry faces (*b* = 0.095, 95%-CI [0.068 0.123], χ^2^(1) = 45.496, *p* < .001; Fig S1). Participants learned the correct response of each mini-block (Fig S1; main effect of required response, *b* = 1.101, 95%-CI [1.047, 1.155], χ^2^(1) = 1607.341, *p* < .001; mean/median accuracy per participant 0.71, range 0.55–0.81), with better performance in the 2^nd^ compared to the 1^st^ half of mini-blocks (2-way interaction required response x mini-block half; b = 0.663, 95%-CI [0.620 0.707], χ^2^(1) = 893.337, *p* < .001).

Again, a simple reinforcement learning model reproduced the behavioural findings. Again, only M4 was qualitatively able to produce the kind of emotional biases observed in the online data. The fitted parameter values suggested again that participants learned faster from positive than negative feedback (α_POS_: *M* = 0.665, *SD* = 0.145, range 0.119–0.877; α_NEG_: *M* = 0.321, *SD* = 0.152, range 0.044–0.829; paired-samples *t*-test: *t*(209) = 26.214, *p* < .001, *d* = 1.809) and showed greater approach tendencies for happy over neutral and neutral over angry face stimuli (γ_HAPPY_: *M* = 0.031, *SD* = 0.033, range -0.066–0.153; γ_NEUTRAL_: *M* = 0.016, *SD* = 0.028, range -0.054–0.111; γ_ANGRY_: *M* = -0.009, *SD* = 0.022, range -0.093–0.057; paired-samples *t*-test happy – neutral: *t*(209) = 6.870, *p* < .001, *d* = 0.474; paired-samples *t*-test neutral – angry: *t*(209) = 11.208, *p* < .001, *d* = 0.773).

Second, mimicking the analysis performed in the in-person cohort, we analysed reaction times of the online cohort as a function of the emotion of the displayed face (happy/neutral/angry), feedback on the previous trial (positive/negative), and mini-block half (first/second) using mixed-effects linear regression. Again, participants displayed an emotional bias also in their RTs (main effect of emotion; χ^2^(2) = 36.457, *p* < .001), with faster responses to happy faces than to neutral faces (b = -0.025, 95% CI [-0.034 -0.016], χ^2^(1) = 30.267, *p* < .001) and angry faces (b = -0.022, 95% CI [-0.031 -0.014], χ^2^(1) = 26.771, *p* < .001), but no difference between neutral and angry faces (*b* = 0.003, 95% CI [-0.005 0.011], χ^2^(1) = 0.599, *p* = .439; Fig S1). Regarding the RT evolution across trials in a mini-block (Fig S1), on trial #1 of mini-blocks, RTs were in fact slowest for neutral faces compared to both happy faces (*b* = -0.039, 95% CI [-0.057 -0.022], χ^2^(1) = 19.372, *p* < .001) and angry faces (*b* = 0.045, 95% CI [0.029 0.060], χ^2^(1) = 32.182, *p* < .001), potentially reflecting that those faces were most ambiguous and made the decision whether a new mini-block had started or not harder than happy or angry faces did. From trial #2 onwards, the normal emotional bias emerged, with faster response to happy than neutral and angry faces. Overall, reaction times were fastest on trial #2 of a block, slowed down towards the end of a mini-block, and were slowest on trial #1 of a new mini-block (Fig S1).

Third, we analysed participants’ choices to repeat or switch their responses as a function of the feedback they had received on the previous trial (positive/negative) and the mini-block half (first/second). Participants showed an overall tendency to repeat responses (intercept: *b* = 1.846, 95% CI [1.725 1.966], χ^2^(1) = 902.930, *p* < .001), repeated their responses more often after positive than negative feedback (main effect of feedback: *b* = 1.844, 95% CI [1.757 1.931], χ^2^(1) = 1721.827, *p* < .001) and more in the 2^nd^ than in the 1^st^ half of mini-blocks (main effect of block-half: *b* = 0.181, 95% CI [0.144 0.217], χ^2^(1) = 94.152, *p* < .001; Fig XXX).

Taken together, we replicated the key effects shown by participants in the in-person sessions in the online participants. Most importantly, online participants showed an emotional bias in responses and RTs, learned the task, and adjusted their behaviour to feedback. The only notable deviations from the in-person findings was an altered pattern of RTs on the first trial of a new mini-block, on which online participants were slowest for neutral faces (which were likely most ambiguous and made it hard to conclusively detect the start of a new mini-block) compared to happy and angry faces.

#### Additional model fitting results for in-person cohort (Exp1)

Average model parameters looked as follows: parameter estimates from M4 (**Fig1f**) confirmed faster learning from positive over negative feedback (α_POS_: *M* = 0.616, *SD* = 0.121, range 0.166 – 0.821; α_NEG_: *M* = 0.254, *SD* = 0.074, range 0.120 – 0.514; paired-samples *t*-test: *t*(86) = 25.105, *p* < .001, *d* = 2.692) and greater approach tendencies for happy over neutral and neutral over angry face stimuli (γ_HAPPY_: *M* = 0.046, *SD* = 0.026, range -0.008 – 0.112; γ_NEUTRAL_: *M* = 0.032, *SD* = 0.025, range -0.032 – 0.094; γ_ANGRY_: *M* = 0.011, *SD* = 0.027, range -0.034 – 0.092; paired-samples *t*-test happy – neutral: *t*(86) = 5.181, *p* < .001, *d* = 0.555; paired-samples *t*-test neutral – angry: *t*(89) = 5.990, *p* < .001, *d* = 0.642).

Evidence from model parameters supported our claim made in the main part of the manuscript that BLA-TUS did not affect learning from feedback: There was neither a change in the positive learning rate (*t*(19) = 0.354, *p* = 0.728, *d* = 0.079) nor the negative learning rate (*t*(19) = 0.026, *p* = 0.980, *d* = 0.006), and for neither of these parameters was the difference between amygdala and sham sessions correlated with the amount of bilaterally stimulated BLA volume (positive learning rate: *r*(18) = -0.092, *p* = 0.898; negative learning rate: *r*(18) = 0.219, *p* = 0.354).

#### Full results for all regression results reported in the manuscript

##### Overview tables behavioural results collapsed over TUS conditions, n = 29

| Effect | *b* | 95% CI | | *df* | χ^2^ | *p* |
| --- | --- | --- | --- | --- | --- | --- |
| DV: approach/avoid response |  |  |  |  |  |  |
| Intercept | 0.164 | 0.140 | 0.188 | 1 | 180.652 | < .001 |
| Required response | 1.169 | 1.074 | 1.263 | 1 | 586.616 | < .001 |
| Emotion | 0.125 |  |  | 2 | 23.863 | < .001 |
| *Happy – neutral* | 0.059 | 0.018 | 0.010 | 1 | 7.792 | .005 |
| *Happy – angry* | 0.132 | 0.078 | 0.186 | 1 | 23.111 | < .001 |
| *Neutral – angry* | 0.073 | 0.033 | 0.112 | 1 | 13.074 | < .001 |
| Mini-block half | -0.062 | -0.103 | -0.021 | 1 | 8.693 | .003 |
| Required response x emotion |  |  |  | 2 | 1.298 | .523 |
| *Required response x happy – neutral* | -0.019 | -0.53 | 0.015 | 1 | 1.230 | .267 |
| *Required response x happy – angry* | -0.006 | -0.042 | 0.029 | 1 | 0.113 | .737 |
| *Required response x neutral – angry* | 0.012 | -0.027 | 0.051 | 1 | 0.361 | .548 |
| Required response x mini-block half | 0.706 | 0.627 | 0.785 | 1 | 306.010 | < .001 |
| Emotion x mini-block half |  |  |  | 2 | 6.304 | .043 |
| *Happy – neutral x mini-block half* | -0.008 | -0.042 | 0.026 | 1 | 0.197 | .657 |
| *Happy – angry x mini-block half* | -0.040 | -0.073 | 0.007 | 1 | 5.766 | .016 |
| *Neutral – angry x mini-block half* | -0.033 | -0.064 | 0.001 | 1 | 4.189 | .041 |
| Required response x emotion x mini-block half |  |  |  | 2 | 6.210 | .045 |
| *Required response x happy – neutral x mini-block half* | 0.012 | -0.021 | 0.045 | 1 | 0.515 | .473 |
| *Required response x happy – angry x mini-block half* | 0.045 | 0.013 | 0.077 | 1 | 7.593 | .006 |
| *Required response x neutral – angry x mini-block half* | 0.032 | 0.004 | 0.067 | 1 | 3.059 | .080 |
| *Overview results from mixed-effects logistic regression models*. Mixed-effects linear regression models were fit to participants’ approach/avoid choices with the glmer() function from lme4 package in R. The dependent variable was coded as approach = 1, avoid = 0. All factors were coded with sum-to-zero coding (required response: approach = 1, avoid = -1; emotion: happy/neutral/angry: 1/0/-1 and 0/1/-1; mini-block half: early = -1, late = 1) such that regression coefficients can be interpreted as standardized regression coefficients. Significant effects involving emotion were followed up by eliminating one factor level refitting the model with only the two remaining levels, respectively (indented; in italics). | | | | | | |

**Table S10:** Regression on choices to approach/avoid collapsed across all 3 sessions (n = 87 sessions in n = 29 participants) for the in-person cohort of Experiment 1.

| Effect | *b* | 95% CI | | *df* | χ^2^ | *p* |
| --- | --- | --- | --- | --- | --- | --- |
| DV: RTs |  |  |  |  |  |  |
| Intercept | -0.064 | -0.193 | .066 | 1 | 0.927 | 0.336 |
| Emotion |  |  |  | 2 | 12.841 | .002 |
| *Happy – neutral* | -0.019 | -0.032 | -0.006 | 1 | 8.038 | .005 |
| *Happy – angry* | -0.021 | -0.033 | -0.009 | 1 | 11.542 | < .001 |
| *Neutral – angry* | -0.001 | -0.014 | 0.012 | 1 | 0.021 | .884 |
| Feedback last trial | 0.063 | 0.045 | 0.081 | 1 | 47.311 | < .001 |
| Mini-block half | 0.033 | 0.018 | 0.048 | 1 | 18.233 | < .001 |
| Emotion x feedback last trial |  |  |  | 2 | 6.326 | .042 |
| *Happy – neutral x feedback last trial* | -0.005 | -0.019 | 0.009 | 1 | 0.451 | .502 |
| *Happy – angry x feedback last trial* | 0.011 | -0.001 | 0.024 | 1 | 3.072 | .080 |
| *Neutral – angry x feedback last trial* | 0.016 | 0.003 | 0.029 | 1 | 5.632 | .018 |
| Emotion x mini-block half |  |  |  | 2 | 1.925 | .382 |
| *Happy – neutral x mini-block half* | -0.002 | -0.014 | 0.009 | 1 | 0.153 | .695 |
| *Happy – angry x mini-block half* | -0.009 | -0.021 | 0.004 | 1 | 1.891 | .169 |
| *Neutral – angry x mini-block half* | -0.006 | -0.018 | 0.006 | 1 | 1.013 | .314 |
| Feedback last trial x mini-block half | 0.005 | -0.005 | 0.015 | 1 | 0.887 | .346 |
| Emotion x feedback last trial x mini-block half |  |  |  | 2 | 2.651 | .266 |
| *Happy – neutral x feedback last trial x mini-block half* | -0.012 | -0.027 | 0.003 | 1 | 2.542 | .111 |
| *Happy – angry x feedback last trial x mini-block half* | -0.003 | -0.018 | 0.012 | 1 | 0.182 | .670 |
| *Neutral – angry x feedback last trial x mini-block half* | 0.009 | -0.005 | 0.023 | 1 | 1.461 | .227 |
| *Overview results from mixed-effects linear regression models*. Mixed-effects linear regression models were fit to participants’ reaction times (RTs) with the lmer() function from lme4 package in R. The dependent variable was z-standardized. All factors were coded with sum-to-zero coding (required response: approach = 1, avoid = -1; emotion: happy/neutral/angry: 1/0/-1 and 0/1/-1; mini-block half: early = -1, late = 1) such that regression coefficients can be interpreted as standardized regression coefficients. Significant effects involving emotion were followed up by eliminating one factor level refitting the model with only the two remaining levels, respectively (indented; in italics). | | | | | | |

**Table S11:** Regression on RTs collapsed across all 3 sessions (n = 87 sessions in n = 29 participants) for the in-person cohort of Experiment 1.

| Effect | *b* | 95% CI | | *df* | χ^2^ | *p* |
| --- | --- | --- | --- | --- | --- | --- |
| DV: repeat/switch response |  |  |  |  |  |  |
| Intercept | 1.870 | 1.635 | 2.105 | 1 | 244.115 | < .001 |
| Feedback last trial | 1.743 | 1.554 | 1.932 | 1 | 328.319 | < .001 |
| Mini-block half | 0.217 | 0.125 | 0.308 | 1 | 21.373 | < .001 |
| Feedback last trial x mini-block half | -0.020 | -0.099 | 0.058 | 1 | 0.261 | .609 |
| *Overview results from mixed-effects logistic regression models*. Mixed-effects linear regression models were fit to participants’ repeat/switch choices (with respect to the previous response) with the glmer() function from lme4 package in R. Note that the first trial of each mini-block was omitted from these analyses. The dependent variable was coded as repeat = 1, switch = 0. All factors were coded with sum-to-zero coding (feedback last trial: positive = 1, negative = -1; mini-block half: early = -1, late = 1) such that regression coefficients can be interpreted as standardized regression coefficients. | | | | | | |

**Table S12:** Regression on choices to repeat/switch the previous response collapsed across all 3 sessions (n = 87 sessions in n = 29 participants) for the in-person cohort of Experiment 1.

##### Overview tables behavioural results online sample, n = 210

| Effect | *b* | 95% CI | | *df* | χ^2^ | *p* |
| --- | --- | --- | --- | --- | --- | --- |
| DV: approach/avoid response |  |  |  |  |  |  |
| Intercept | 0.056 | 0.026 | 0.087 | 1 | 12.918 | < .001 |
| Required response | 1.101 | 1.047 | 1.155 | 1 | 1607.341 | < .001 |
| Emotion |  |  |  | 2 | 80.473 | < .001 |
| *Happy – neutral* | 0.050 | 0.027 | 0.074 | 1 | 17.207 | < .001 |
| *Happy – angry* | 0.145 | 0.113 | 0.177 | 1 | 79.459 | < .001 |
| *Neutral – angry* | 0.095 | 0.068 | 0.123 | 1 | 45.496 | < .001 |
| Mini-block half | -0.037 | -0.055 | -0.019 | 1 | 16.710 | < .001 |
| Required response x emotion |  |  |  | 2 | 4.389 | .111 |
| *Required response x happy – neutral* | -0.004 | -0.025 | 0.018 | 1 | 0.103 | .748 |
| *Required response x happy – angry* | 0.016 | -0.006 | 0.038 | 1 | 2.099 | .147 |
| *Required response x neutral – angry* | 0.019 | 0.001 | 0.039 | 1 | 3.872 | .049 |
| Required response x mini-block half | 0.663 | 0.620 | 0.707 | 1 | 893.337 | < .001 |
| Emotion x mini-block half |  |  |  | 2 | 5.619 | .060 |
| *Happy – neutral x mini-block half* | -0.019 | -0.038 | -0.001 | 1 | 3.993 | .046 |
| *Happy – angry x mini-block half* | -0.019 | -0.038 | -0.001 | 1 | 4.025 | .045 |
| *Neutral – angry x mini-block half* | 0.002 | -0.017 | 0.022 | 1 | 0.029 | .866 |
| Required response x emotion x mini-block half |  |  |  | 2 | 29.846 | < .001 |
| *Required response x happy – neutral x mini-block half* | 0.047 | 0.027 | 0.066 | 1 | 22.246 | < .001 |
| *Required response x happy – angry x mini-block half* | 0.050 | 0.031 | 0.069 | 1 | 26.215 | < .001 |
| *Required response x neutral – angry x mini-block half* | 0.003 | -0.015 | 0.022 | 1 | 0.133 | .716 |
| *Overview results from mixed-effects logistic regression models*. Mixed-effects linear regression models were fit to participants’ approach/avoid choices with the glmer() function from lme4 package in R. The dependent variable was coded as approach = 1, avoid = 0. All factors were coded with sum-to-zero coding (required response: approach = 1, avoid = -1; emotion: happy/neutral/angry: 1/0/-1 and 0/1/-1; mini-block half: early = -1, late = 1) such that regression coefficients can be interpreted as standardized regression coefficients. Significant effects involving emotion were followed up by eliminating one factor level refitting the model with only the two remaining levels, respectively (indented; in italics). | | | | | | |

**Table S13:** Regression on choices to approach/avoid for the online cohort (n = 210) of Experiment 2.

| Effect | *b* | 95% CI | | *df* | χ^2^ | *p* |
| --- | --- | --- | --- | --- | --- | --- |
| DV: RTs |  |  |  |  |  |  |
| Intercept | -0.085 | -0.137 | -0.033 | 1 | 10.113 | .001 |
| Emotion |  |  |  | 2 | 36.457 | < .001 |
| *Happy – neutral* | -0.025 | -0.034 | -0.016 | 1 | 30.267 | < .001 |
| *Happy – angry* | -0.022 | -0.031 | -0.014 | 1 | 26.771 | < .001 |
| *Neutral – angry* | 0.003 | -0.005 | 0.011 | 1 | 0.599 | .439 |
| Feedback last trial | 0.043 | 0.034 | 0.052 | 1 | 82.652 | < .001 |
| Mini-block half | 0.033 | 0.025 | 0.040 | 1 | 69.760 | < .001 |
| Emotion x feedback last trial |  |  |  | 2 | 0.806 | .668 |
| *Happy – neutral x feedback last trial* | -0.001 | -0.008 | 0.007 | 1 | 0.010 | .919 |
| *Happy – angry x feedback last trial* | -0.003 | -0.010 | 0.004 | 1 | 0.705 | .401 |
| *Neutral – angry x feedback last trial* | -0.002 | -0.010 | 0.005 | 1 | 0.404 | .525 |
| Emotion x mini-block half |  |  |  | 2 | 0.360 | .835 |
| *Happy – neutral x mini-block half* | 0.001 | -0.007 | 0.007 | 1 | 0.001 | .993 |
| *Happy – angry x mini-block half* | -0.002 | -0.009 | 0.005 | 1 | 0.283 | .595 |
| *Neutral – angry x mini-block half* | -0.002 | -0.010 | 0.005 | 1 | 0.321 | .571 |
| Feedback last trial x mini-block half | 0.010 | 0.004 | 0.016 | 1 | 11.173 | < .001 |
| Emotion x feedback last trial x mini-block half |  |  |  | 2 | 0.103 | .950 |
| *Happy – neutral x feedback last trial x mini-block half* | 0.001 | -0.007 | 0.008 | 1 | 0.029 | .865 |
| *Happy – angry x feedback last trial x mini-block half* | -0.001 | -0.008 | 0.006 | 1 | 0.054 | .817 |
| *Neutral – angry x feedback last trial x mini-block half* | -0.001 | -0.008 | 0.006 | 1 | 0.112 | .738 |
| *Overview results from mixed-effects linear regression models*. Mixed-effects linear regression models were fit to participants’ reaction times (RTs) with the lmer() function from lme4 package in R. The dependent variable was z-standardized. All factors were coded with sum-to-zero coding (required response: approach = 1, avoid = -1; emotion: happy/neutral/angry: 1/0/-1 and 0/1/-1; mini-block half: early = -1, late = 1) such that regression coefficients can be interpreted as standardized regression coefficients. Significant effects involving emotion were followed up by eliminating one factor level refitting the model with only the two remaining ones, respectively (indented; in italics). | | | | | | |

**Table S14:** Regression on RTs for the online cohort (n = 210) of Experiment 2.

| Effect | *b* | 95% CI | | *df* | χ^2^ | *p* |
| --- | --- | --- | --- | --- | --- | --- |
| DV: repeat/switch response |  |  |  |  |  |  |
| Intercept | 1.846 | 1.725 | 1.966 | 1 | 902.930 | < .001 |
| Feedback last trial | 1.844 | 1.757 | 1.931 | 1 | 1721.827 | < .001 |
| Mini-block half | 0.181 | 0.144 | 0.217 | 1 | 94.152 | < .001 |
| Feedback last trial x mini-block half | -0.010 | -0.040 | 0.021 | 1 | 0.391 | .532 |
| *Overview results from mixed-effects logistic regression models*. Mixed-effects linear regression models were fit to participants’ repeat/switch choices (with respect to the previous response) with the glmer() function from lme4 package in R. Note that the first trial of each mini-block was omitted from these analyses. The dependent variable was coded as repeat = 1, switch = 0. All factors were coded with sum-to-zero coding (feedback last trial: positive = 1, negative = -1; mini-block half: early = -1, late = 1) such that regression coefficients can be interpreted as standardized regression coefficients. | | | | | | |

**Table S15:** Regression on choices to repeat/switch the previous response for the online cohort (n = 210) of Experiment 2.

##### Overview tables behavioural results with amygdala vs. sham TUS, n = 20

| Effect | *b* | 95% CI | | *df* | χ^2^ | *p* |
| --- | --- | --- | --- | --- | --- | --- |
| DV: approach/avoid response |  |  |  |  |  |  |
| Intercept | 0.155 | 0.100 | 0.210 | 1 | 30.301 | < .001 |
| Amygdala vs. sham | 0.010 | -0.020 | 0.040 | 1 | 0.443 | .506 |
| Emotion |  |  |  | 2 | 13.204 | .001 |
| *Happy – neutral* | 0.056 | 0.018 | 0.094 | 1 | 8.398 | .004 |
| *Happy – angry* | 0.098 | 0.045 | 0.151 | 1 | 12.978 | <.001 |
| *Neutral – angry* | 0.042 | 0.001 | 0.083 | 1 | 3.940 | .047 |
| Bilaterally stimulated tissue | -0.013 | -0.068 | 0.042 | 1 | 0.221 | .638 |
| Amygdala vs. sham x emotion |  |  |  | 2 | 1.359 | .507 |
| *Amygdala vs. sham x happy – neutral* | -0.018 | -0.051 | 0.016 | 1 | 1.025 | .311 |
| *Amygdala vs. sham x happy – angry* | -0.023 | -0.067 | 0.020 | 1 | 1.111 | .292 |
| *Amygdala vs. sham x neutral – angry* | -0.005 | -0.043 | 0.033 | 1 | 0.069 | .793 |
| *Amygdala vs. sham for happy* | -0.017 | -0.068 | 0.034 | 1 | 0.420 | .517 |
| *Amygdala vs. sham for neutral* | 0.019 | -0.032 | 0.069 | 1 | 0.534 | .465 |
| *Amygdala vs. sham for angry* | 0.029 | -0.023 | 0.082 | 1 | 1.212 | .271 |
| Amygdala vs. sham x bilaterally stimulated tissue | 0.008 | -0.022 | 0.038 | 1 | 0.270 | .603 |
| Emotion x bilaterally stimulated tissue |  |  |  | 2 | 2.201 | .333 |
| *Happy – neutral* *x bilaterally stimulated tissue* | -0.009 | -0.048 | 0.029 | 1 | 0.235 | .628 |
| *Happy – angry x bilaterally stimulated tissue* | 0.023 | -0.030 | 0.076 | 1 | 0.704 | .401 |
| *Neutral – angry x bilaterally stimulated tissue* | 0.033 | -0.008 | 0.075 | 1 | 2.449 | .118 |
| *Bilaterally stimulated tissue for happy* | -0.004 | -0.085 | 0.077 | 1 | 0.011 | .917 |
| *Bilaterally stimulated tissue for neutral* | 0.015 | -0.068 | 0.098 | 1 | 0.215 | .724 |
| *Bilaterally stimulated tissue for angry* | -0.052 | -0.099 | -0.005 | 1 | 4.739 | .029 |
| Amygdala vs. sham x emotion x bilaterally stimulated tissue |  |  |  | 2 | 10.079 | .006 |
| *Amygdala vs. sham x happy – neutral x bilaterally stimulated tissue* | -0.048 | -0.082 | -0.014 | 1 | 7.495 | .006 |
| *Amygdala vs. sham x happy – angry x bilaterally stimulated tissue* | 0.003 | -0.041 | 0.046 | 1 | 0.017 | .897 |
| *Amygdala vs. sham x neutral – angry x bilaterally stimulated tissue* | 0.051 | 0.013 | 0.089 | 1 | 6.971 | .008 |
| *Amygdala vs. sham x bilaterally stimulated tissue for happy* | -0.022 | -0.074 | 0.030 | 1 | 0.701 | .403 |
| *Amygdala vs. sham x bilaterally stimulated tissue for neutral* | 0.074 | 0.023 | 0.124 | 1 | 8.145 | .004 |
| *Amygdala vs. sham x bilaterally stimulated tissue for angry* | -0.028 | -0.081 | 0.024 | 1 | 1.111 | .292 |
| *Overview results from mixed-effects logistic regression models*. Mixed-effects linear regression models were fit to participants’ approach/avoid choices with the glmer() function from lme4 package in R. The dependent variable was coded as approach = 1, avoid = 0. All factors were coded with sum-to-zero coding (required response: approach = 1, avoid = -1; emotion: happy/neutral/angry: 1/0/-1 and 0/1/-1; mini-block half: early = -1, late = 1) such that regression coefficients can be interpreted as standardized regression coefficients. Significant effects involving emotion were followed up by eliminating one factor level refitting the model with only the two remaining levels, respectively (indented; in italics). | | | | | | |

**Table S16:** Regression on choices to approach/avoid for the in-person cohort considered for BLA sessions (n = 20).

| Effect | *b* | 95% CI | | *df* | χ^2^ | *p* |
| --- | --- | --- | --- | --- | --- | --- |
| DV: RTs |  |  |  |  |  |  |
| Intercept | -0.051 | -0.216 | 0.114 | 1 | 0.363 | .547 |
| Amygdala vs. sham | 0.036 | -0.009 | 0.081 | 1 | 2.482 | .115 |
| Emotion |  |  |  | 2 | 11.105 | .004 |
| *Happy – neutral* | -0.031 | -0.051 | -0.011 | 1 | 8.906 | .003 |
| *Happy – angry* | -0.024 | -0.042 | -0.006 | 1 | 7.081 | .008 |
| *Neutral – angry* | 0.007 | -0.013 | 0.028 | 1 | 0.504 | .478 |
| Feedback last trial | 0.060 | 0.042 | 0.079 | 1 | 39.757 | < .001 |
| Bilaterally stimulated tissue | 0.004 | -0.162 | 0.169 | 1 | 0.002 | .965 |
| Amygdala vs. sham x emotion |  |  |  | 2 | 6.485 | .039 |
| *Amygdala vs. sham x happy – neutral* | -0.001 | -0.019 | 0.018 | 1 | 0.001 | .985 |
| *Amygdala vs. sham x happy – angry* | 0.025 | 0.003 | 0.048 | 1 | 4.766 | .029 |
| *Amygdala vs. sham x neutral – angry* | 0.025 | 0.004 | 0.045 | 1 | 5.743 | .017 |
| *Amygdala vs. sham for happy* | 0.054 | 0.005 | 0.104 | 1 | 4.585 | .032 |
| *Amygdala vs. sham for neutral* | 0.051 | 0.004 | 0.097 | 1 | 4.555 | .033 |
| *Amygdala vs. sham for angry* | 0.003 | -0.048 | 0.053 | 1 | 0.012 | .914 |
| Amygdala vs. sham x feedback last trial | 0.003 | -0.014 | 0.034 | 1 | 0.117 | .733 |
| Amygdala vs. sham x bilaterally stimulated tissue | 0.005 | -0.040 | 0.050 | 1 | 0.055 | .815 |
| Emotion x feedback last trial |  |  |  | 2 | 2.372 | .306 |
| *Happy – neutral x feedback last trial* | -0.001 | -0.020 | 0.017 | 1 | 0.016 | .898 |
| *Happy – angry x feedback last trial* | 0.013 | -0.005 | 0.031 | 1 | 2.040 | .153 |
| *Neutral – angry x feedback last trial* | 0.013 | -0.007 | 0.034 | 1 | 1.634 | .201 |
| Emotion x bilaterally stimulated tissue |  |  |  | 2 | 0.603 | .740 |
| *Happy – neutral* *x bilaterally stimulated tissue* | -0.007 | -0.028 | 0.013 | 1 | 0.508 | .476 |
| *Happy – angry x bilaterally stimulated tissue* | -0.001 | -0.019 | 0.017 | 1 | 0.009 | .926 |
| *Neutral – angry x bilaterally stimulated tissue* | 0.007 | -0.014 | 0.028 | 1 | 0.430 | .512 |
| Feedback last trial x bilaterally stimulated tissue | 0.007 | -0.011 | 0.026 | 1 | 0.597 | .440 |
| Amygdala vs. sham x emotion x feedback last trial |  |  |  | 2 | 1.489 | .475 |
| *Amygdala vs. sham x happy – neutral x feedback last trial* | 0.008 | -0.012 | 0.028 | 1 | 0.581 | .446 |
| *Amygdala vs. sham x happy – angry x feedback last trial* | -0.005 | -0.027 | 0.018 | 1 | 0.151 | .697 |
| *Amygdala vs. sham x neutral – angry x feedback last trial* | -0.013 | -0.033 | 0.008 | 1 | 1.475 | .225 |
| *Amygdala vs. sham x feedback last trial for happy* | 0.006 | -0.019 | 0.030 | 1 | 0.189 | .663 |
| *Amygdala vs. sham x feedback last trial for neutral* | -0.010 | -0.039 | 0.019 | 1 | 0.451 | .502 |
| *Amygdala vs. sham x feedback last trial for angry* | 0.015 | -0.016 | 0.045 | 1 | 0.904 | .342 |
| Amygdala vs. sham x emotion x bilaterally stimulated tissue |  |  |  | 2 | 0.062 | .969 |
| *Amygdala vs. sham x happy – neutral x bilaterally stimulated tissue* | 0.002 | -0.016 | 0.021 | 1 | 0.053 | .819 |
| *Amygdala vs. sham x happy – angry x bilaterally stimulated tissue* | 0.001 | -0.022 | 0.023 | 1 | 0.003 | .955 |
| *Amygdala vs. sham x neutral – angry x bilaterally stimulated tissue* | -0.002 | -0.022 | 0.019 | 1 | 0.024 | .876 |
| *Amygdala vs. sham x bilaterally stimulated tissue for happy* | 0.007 | -0.042 | 0.057 | 1 | 0.082 | .774 |
| *Amygdala vs. sham x bilaterally stimulated tissue for neutral* | 0.003 | -0.044 | 0.049 | 1 | 0.012 | .912 |
| *Amygdala vs. sham x bilaterally stimulated tissue for angry* | 0.006 | -0.044 | 0.057 | 1 | 0.064 | .800 |
| Amygdala vs. sham x feedback last trial x bilaterally stimulated tissue | 0.004 | -0.013 | 0.020 | 1 | 0.181 | .671 |
| Emotion x feedback last trial x bilaterally stimulated tissue |  |  |  | 2 | 2.052 | .358 |
| *Happy – neutral x feedback last trial x bilaterally stimulated tissue* | -0.009 | -0.028 | 0.009 | 1 | 0.935 | .334 |
| *Happy – angry x feedback last trial x bilaterally stimulated tissue* | -0.013 | -0.031 | 0.005 | 1 | 2.058 | .151 |
| *Neutral – angry x feedback last trial x bilaterally stimulated tissue* | -0.003 | -0.024 | 0.017 | 1 | 0.093 | .761 |
| Amygdala vs. sham x emotion x feedback last trial x bilaterally stimulated tissue |  |  |  | 2 | 3.128 | .209 |
| *Amygdala vs. sham x happy – neutral x feedback last trial x bilaterally stimulated tissue* | 0.004 | -0.016 | 0.024 | 1 | 0.147 | .702 |
| *Amygdala vs. sham x happy – angry x feedback last trial x bilaterally stimulated tissue* | 0.020 | -0.003 | 0.043 | 1 | 3.048 | .081 |
| *Amygdala vs. sham x neutral – angry x feedback last trial x bilaterally stimulated tissue* | 0.016 | -0.005 | 0.036 | 1 | 2.214 | .137 |
| *Amygdala vs. sham x bilaterally stimulated tissue for happy* | 0.020 | -0.005 | 0.045 | 1 | 2.488 | .115 |
| *Amygdala vs. sham x feedback last trial x bilaterally stimulated tissue for neutral* | 0.012 | -0.017 | 0.041 | 1 | 0.651 | .420 |
| *Amygdala vs. sham x feedback last trial x bilaterally stimulated tissue for angry* | -0.021 | -0.052 | 0.009 | 1 | 1.850 | .174 |
| *Overview results from mixed-effects linear regression models*. Mixed-effects linear regression models were fit to participants’ reaction times (RTs) with the lmer() function from lme4 package in R. The dependent variable was z-standardized. All factors were coded with sum-to-zero coding (required response: approach = 1, avoid = -1; emotion: happy/neutral/angry: 1/0/-1 and 0/1/-1; mini-block half: early = -1, late = 1) such that regression coefficients can be interpreted as standardized regression coefficients. Significant effects involving emotion were followed up by eliminating one factor level refitting the model with only the two remaining levels, respectively (indented; in italics). | | | | | | |

**Table S17:** Regression on RTs for the in-person cohort considered for BLA sessions (n = 20).

| Effect | *b* | 95% CI | | *df* | χ^2^ | *p* |
| --- | --- | --- | --- | --- | --- | --- |
| DV: repeat/switch response |  |  |  |  |  |  |
| Intercept | 1.881 | 1.586 | 2.175 | 1 | 156.560 | < .001 |
| Amygdala vs. sham | 0.005 | -0.136 | 0.145 | 1 | 0.004 | .948 |
| Feedback last trial | 1.793 | 1.590 | 1.996 | 1 | 299.819 | < .001 |
| Mini-block half | 0.209 | 0.091 | 0.327 | 1 | 12.019 | < .001 |
| Bilaterally stimulated tissue | -0.122 | -0.416 | 0.172 | 1 | 0.661 | .416 |
| Amygdala vs. sham x feedback last trial | -0.044 | -0.140 | 0.051 | 1 | 0.827 | .363 |
| Amygdala vs. sham x mini-block half | -0.018 | -0.070 | 0.034 | 1 | 0.457 | .499 |
| Amygdala vs. sham x bilaterally stimulated tissue | -0.009 | -0.149 | 0.131 | 1 | 0.016 | .900 |
| Feedback last trial x bilaterally stimulated tissue | -0.127 | -0.329 | 0.076 | 1 | 1.505 | .220 |
| Mini-block half x bilaterally stimulated tissue | 0.011 | -0.107 | 0.129 | 1 | 0.033 | .855 |
| Amygdala vs. sham x feedback last trial x bilaterally stimulated tissue | -0.026 | -0.120 | 0.069 | 1 | 0.282 | .596 |
| Amygdala vs. sham x mini-block half x bilaterally stimulated tissue | 0.038 | -0.014 | 0.090 | 1 | 2.075 | .150 |
| *Overview results from mixed-effects logistic regression models*. Mixed-effects linear regression models were fit to participants’ approach/avoid choices with the glmer() function from lme4 package in R. The dependent variable was coded as approach = 1, avoid = 0. All factors were coded with sum-to-zero coding (required response: approach = 1, avoid = -1; emotion: happy/neutral/angry: 1/0/-1 and 0/1/-1; mini-block half: early = -1, late = 1) such that regression coefficients can be interpreted as standardized regression coefficients. Significant effects involving emotion were followed up by eliminating one factor level refitting the model with only the two remaining levels, respectively (indented; in italics). | | | | | | |

**Table S18:** Regression on choices to repeat/switch the previous response for the in-person cohort considered for BLA sessions (n = 20).

##### Overview tables behavioural results with insula vs. sham TUS, N = 29

| Effect | *b* | 95% CI | | *Df* | χ^2^ | *p* |
| --- | --- | --- | --- | --- | --- | --- |
| DV: approach/avoid response |  |  |  |  |  |  |
| Intercept | 0.140 | 0.098 | 0.181 | 1 | 44.152 | < .001 |
| Insula vs. sham | 0.002 | -0.025 | 0.029 | 1 | 0.020 | .888 |
| Emotion |  |  |  | 2 | 43.103 | < .001 |
| *Happy – neutral* | 0.059 | 0.027 | 0.090 | 1 | 13.537 | < .001 |
| *Happy – angry* | 0.123 | 0.086 | 0.160 | 1 | 43.286 | < .001 |
| *Neutral – angry* | 0.064 | 0.035 | 0.093 | 1 | 18.698 | < .001 |
| Insula vs. sham x emotion |  |  |  | 2 | 0.172 | .918 |
| *Insula vs. sham x happy – neutral* | -0.005 | -0.033 | 0.024 | 1 | 0.098 | .755 |
| *Insula vs. sham x happy – angry* | -0.008 | -0.047 | 0.032 | 1 | 0.149 | .699 |
| *Insula vs. sham x neutral – angry* | -0.003 | -0.042 | 0.036 | 1 | 0.019 | .890 |
| *Insula vs. sham for happy* | -0.006 | -0.057 | 0.044 | 1 | 0.062 | .804 |
| *Insula vs. sham for neutral* | 0.003 | -0.042 | 0.049 | 1 | 0.019 | .891 |
| *Insula vs. sham for angry* | 0.009 | -0.043 | 0.062 | 1 | 0.125 | .724 |
| *Overview results from mixed-effects logistic regression models*. Mixed-effects linear regression models were fit to participants’ approach/avoid choices with the glmer() function from lme4 package in R. The dependent variable was coded as approach = 1, avoid = 0. All factors were coded with sum-to-zero coding (required response: approach = 1, avoid = -1; emotion: happy/neutral/angry: 1/0/-1 and 0/1/-1; mini-block half: early = -1, late = 1) such that regression coefficients can be interpreted as standardized regression coefficients. Significant effects involving emotion were followed up by eliminating one factor level refitting the model with only the two remaining levels, respectively (indented; in italics). | | | | | | |

**Table S19:** Regression on choices to approach/avoid for the in-person cohort considered for mIns sessions (n = 29).

| Effect | *b* | 95% CI | | *df* | χ^2^ | *p* |
| --- | --- | --- | --- | --- | --- | --- |
| DV: RTs |  |  |  |  |  |  |
| Intercept | -0.066 | -0.203 | 0.071 | 1 | 0.889 | .346 |
| Insula vs. sham | 0.001 | -0.040 | 0.042 | 1 | 0.003 | .960 |
| Emotion |  |  |  | 2 | 13.953 | < .001 |
| *Happy – neutral* | -0.016 | -0.032 | -0.001 | 1 | 4.174 | .041 |
| *Happy – angry* | -0.027 | -0.041 | -0.013 | 1 | 14.873 | < .001 |
| *Neutral – angry* | -0.010 | -0.026 | 0.006 | 1 | 1.616 | .204 |
| Feedback last trial | 0.063 | 0.046 | 0.080 | 1 | 53.571 | < .001 |
| Insula vs. sham x emotion |  |  |  | 2 | 2.575 | .276 |
| *Insula vs. sham x happy – neutral* | 0.013 | -0.006 | 0.032 | 1 | 1.736 | .188 |
| *Insula vs. sham x happy – angry* | 0.013 | -0.003 | 0.030 | 1 | 2.467 | .116 |
| *Insula vs. sham x neutral – angry* | 0.001 | -0.014 | 0.014 | 1 | 0.003 | .958 |
| *Insula vs. sham for happy* | 0.020 | -0.027 | 0.066 | 1 | 0.678 | .410 |
| *Insula vs. sham for neutral* | -0.007 | -0.051 | 0.036 | 1 | 0.113 | .737 |
| *Insula vs. sham for angry* | -0.008 | -0.052 | 0.036 | 1 | 0.128 | .720 |
| Insula vs. sham x feedback last trial | 0.001 | -0.011 | 0.013 | 1 | 0.023 | .879 |
| Emotion x feedback last trial |  |  |  | 2 | 6.283 | .043 |
| *Happy – neutral x feedback last trial* | -0.007 | -0.025 | 0.011 | 1 | 0.564 | .453 |
| *Happy – angry x feedback last trial* | 0.013 | -0.003 | 0.029 | 1 | 2.539 | .111 |
| *Neutral – angry x feedback last trial* | 0.020 | 0.004 | 0.035 | 1 | 5.965 | .015 |
| Insula vs. sham x emotion x feedback last trial |  |  |  | 2 | 0.190 | .909 |
| *Insula vs. sham x happy – neutral x feedback last trial* | -0.002 | -0.017 | 0.013 | 1 | 0.049 | .824 |
| *Insula vs. sham x happy – angry x feedback last trial* | 0.002 | -0.014 | 0.019 | 1 | 0.065 | .799 |
| *Insula vs. sham x neutral – angry x feedback last trial* | 0.003 | -0.013 | 0.019 | 1 | 0.175 | .676 |
| *Insula vs. sham x feedback last trial for happy* | 0.002 | -0.018 | 0.022 | 1 | 0.039 | .844 |
| *Insula vs. sham x feedback last trial for neutral* | 0.005 | -0.016 | 0.026 | 1 | 0.182 | .669 |
| *Insula vs. sham x feedback last trial for angry* | -0.003 | -0.026 | 0.021 | 1 | 0.047 | .828 |
| *Overview results from mixed-effects linear regression models*. Mixed-effects linear regression models were fit to participants’ reaction times (RTs) with the lmer() function from lme4 package in R. The dependent variable was z-standardized. All factors were coded with sum-to-zero coding (required response: approach = 1, avoid = -1; emotion: happy/neutral/angry: 1/0/-1 and 0/1/-1; mini-block half: early = -1, late = 1) such that regression coefficients can be interpreted as standardized regression coefficients. Significant effects involving emotion were followed up by eliminating one factor level refitting the model with only the two remaining levels, respectively (indented; in italics). | | | | | | |

**Table S20:** Regression on RTs for the in-person cohort considered for mIns sessions (n = 29).

| Effect | *b* | 95% CI | | *df* | χ^2^ | *p* |
| --- | --- | --- | --- | --- | --- | --- |
| DV: repeat/switch response |  |  |  |  |  |  |
| Intercept | 1.952 | 1.674 | 2.231 | 1 | 188.466 | < .001 |
| Insula vs. sham | 0.103 | 0.007 | 0.198 | 1 | 4.404 | .036 |
| Feedback last trial | 1.821 | 1.583 | 2.060 | 1 | 223.796 | < .001 |
| Mini-block half | 0.247 | 0.159 | 0.334 | 1 | 30.581 | < .001 |
| Insula vs. sham x feedback last trial | 0.046 | -0.037 | 0.128 | 1 | 1.169 | .280 |
| Insula vs. sham x mini-block half | 0.004 | -0.041 | 0.049 | 1 | 0.035 | .852 |
| *Overview results from mixed-effects logistic regression models*. Mixed-effects linear regression models were fit to participants’ repeat/switch choices (with respect to the previous response) with the glmer() function from lme4 package in R. Note that the first trial of each mini-block was omitted from these analyses. The dependent variable was coded as repeat = 1, switch = 0. All factors were coded with sum-to-zero coding (feedback last trial: positive = 1, negative = -1; mini-block half: early = -1, late = 1) such that regression coefficients can be interpreted as standardized regression coefficients. | | | | | | |

**Table S21:** Regression on choices to repeat/switch the previous response for the in-person cohort considered for mIns sessions (n = 29).

#### Full results for all computational modelling results reported in the manuscript

##### Overview tables computational modelling results collapsed over TUS conditions, n = 29 participants/ n = 87 sessions.

|  | M1 | M2 | M3 | M4 |
| --- | --- | --- | --- | --- |
| α | 0.152 (0.113), 0.055 – 0.572 |  |  |  |
| α_POS_ |  | 0.666 (0.099),  0.188 – 0.826 | 0.643 (0.112),  0.173 – 0.829 | 0.616 (0.121), 0.166 – 0.821 |
| α_NEG_ |  | 0.255 (0.076),  0.118 – 0.533 | 0.257 (0.074),  0.122 – 0.522 | 0.254 (0.074),  0.120 – 0.514 |
| β | 2.989 (0.667), 1.216 – 4.133 | 1.982 (0.359),  1.204 – 2.784 | 2.026 (0.367),  1.251 – 2.838 | 2.068 (0.359),  1.338 – 2.908 |
| γ |  |  | 0.032 (0.024), -0.003 – 0.112 |  |
| γ_HAPPY_ |  |  |  | 0.046 (0.026),  -0.008 – 0.112 |
| γ_NEUTRAL_ |  |  |  | 0.032 (0.025),  -0.032 – 0.094 |
| γ_ANGRY_ |  |  |  | 0.011 (0.027),  -0.034 – 0.092 |
| Mean (standard deviation), minimum – maximum parameter values in computational reinforcement learning models. α = learning rate, β = inverse temperature, γ = approach bias (per emotion). Parameter were fitted with hierarchical Bayesian inference using the CBM toolbox in MATLAB. Trials were treated as nested in 87 sessions. | | | | |

**Table S22:** Parameter estimates obtained from models M1–M4 using the data of all sessions (n = 87, 3 x n = 29) for the in-person TUS cohort.

##### Overview tables computational modelling results online sample, n = 210

|  | M1 | M2 | M3 | M4 |
| --- | --- | --- | --- | --- |
| α | 0.274 (0.245), 0.036 – 0.913 |  |  |  |
| α_POS_ |  | 0.711 (0.123), 0.155 – 0.881 | 0.690 (0.131), 0.144 – 0.876 | 0.665 (0.145), 0.119 – 0.877 |
| α_NEG_ |  | 0.327 (0.155), 0.045 – 0.836 | 0.325 (0.153), 0.045 – 0.833 | 0.321 (0.152), 0.044 – 0.829 |
| β |  | 1.904 (0.455), 0.790 – 3.379 | 1.936 (0.452), 0.808 – 3.476 | 1.978 (0.468), 0.808 – 3.556 |
| γ |  |  | 0.013 (0.028), -0.091 – 0.121 |  |
| γ_HAPPY_ |  |  |  | 0.031 (0.033),  -0.055 – 0.153 |
| γ_NEUTRAL_ |  |  |  | 0.016 (0.028),  -0.054 – 0.111 |
| γ_ANGRY_ |  |  |  | -0.009 (0.022), -0.093 – 0.057 |
| Mean (standard deviation), minimum – maximum parameter values in computational reinforcement learning models. α = learning rate, β = inverse temperature, γ = approach bias (per emotion). Parameter were fitted with hierarchical Bayesian inference using the CBM toolbox in MATLAB. | | | | |

**Table S23:** Parameter estimates obtained from models M1-M4 using the data of the online cohort in Experiment 2.

##### Overview tables computational modelling results with BLA vs. sham TUS, n = 20

|  | M1 | | M2 | | M3 | | M4 | |
| --- | --- | --- | --- | --- | --- | --- | --- | --- |
|  | BLA | sham | BLA | sham | BLA | sham | BLA | sham |
| α | 0.156 (0.141),  0.051 – 0.592 | 0.145 (0.120),  0.056 – 0.539 |  |  |  |  |  |  |
| α_POS_ |  |  | 0.647 (0.086), 0.426 – 0.802 | 0.622 (0.136),  0.162 – 0.810 | 0.624 (0.105), 0.389 – 0.811 | 0.607 (0.139),  0.154 – 0.802 | 0.594 (0.113),  0.374 – 0.800 | 0.584 (0.143),  0.150 – 0.790 |
| α_NEG_ |  |  | 0.257 (0.084),  0.169 – 0.550 | 0.256 (0.086),  0.147 – 0.488 | 0.259 (0.081), 0.170 – 0.539 | 0.257 (0.084),  0.151 – 0.475 | 0.256 (0.080), 0.168 – 0.529 | 0.255 (0.084),  0.149 – 0.475 |
| β | 3.034 (0.656), 1.629 – 3.846 | 3.117 (0.682),  1.649 – 4.192 | 2.021 (0.327),  1.303 – 2.521 | 2.062 (0.328),  1.288 – 2.577 | 2.061 (0.366), 1.336 – 2.831 | 2.098 (0.353),  1.341 – 2.848 | 2.107 (0.355),  1.487 – 2.891 | 2.134 (0.344),  1.511 – 2.895 |
| γ |  |  |  |  | 0.034 (0.025), -0.004 – 0.080 | 0.028 (0.021),  -0.012 – 0.064 |  |  |
| γ_HAPPY_ |  |  |  |  |  |  | 0.042 (0.033), -0.021 – 0.117 | 0.046 (0.031),  0.005 – 0.131 |
| γ_NEUTRAL_ |  |  |  |  |  |  | 0.031 (0.035), -0.020 – 0.107 | 0.027 (0.023),  -0.018 – 0.063 |
| γ_ANGRY_ |  |  |  |  |  |  | 0.023 (0.031), -0.022 – 0.081 | 0.007 (0.026),  -0.035 – 0.061 |
| Mean (standard deviation), minimum – maximum parameter values in computational reinforcement learning models. α = learning rate, β = inverse temperature, γ = approach bias (per emotion). Parameter were fitted with hierarchical Bayesian inference using the CBM toolbox in MATLAB. Values are fitted and reported separately for BLA and sham sessions. | | | | | | | | |

**Table S24:** Parameter estimates obtained from models M1-M4 for the in-person TUS cohort, comparing sham and BLA sonication conditions (n = 20).

##### Overview tables computational modelling results with mIns vs. sham TUS, N = 29

|  | M1 | | M2 | | M3 | | M4 | |
| --- | --- | --- | --- | --- | --- | --- | --- | --- |
|  | mIns | sham | mIns | sham | mIns | sham | mIns | sham |
| α | 0.146 (0.085),  0.063 – 0.412 | 0.149 (0.094),  0.064 – 0.506 |  |  |  |  |  |  |
| α_POS_ |  |  | 0.693 (0.089),  0.370 – 0.805 | 0.624 (0.131),  0.165 – 0.807 | 0.676 (0.093),  0.358 – 0.801 | 0.608 (0.132),  0.158 – 0.797 | 0.660 (0.092),  0.349 – 0.789 | 0.583 (0.136),  0.154 – 0.783 |
| α_NEG_ |  |  | 0.258 (0.059),  0.156 – 0.419 | 0.247 (0.087),  0.097 – 0.490 | 0.262 (0.058),  0.161 – 0.420 | 0.248 (0.084),  0.102 – 0.476 | 0.262 (0.058),  0.172 – 0.418 | 0.247 (0.083),  0.106 – 0.473 |
| β | 3.046 (0.740),  1.472 – 4.172 | 2.960 (0.545),  1.703 – 3.943 | 2.031 (0.377),  1.286 – 2.796 | 2.024 (0.346),  1.292 – 2.573 | 2.065 (0.390),  1.309 – 2.839 | 2.058 (0.363),  1.313 – 2.845 | 2.091 (0.387),  1.378 – 2.851 | 2.097 (0.354),  1.391 – 2.893 |
| γ |  |  |  |  | 0.029 (0.025),  -0.023 – 0.102 | 0.028 (0.019),  -0.011 – 0.061 |  |  |
| γ_HAPPY_ |  |  |  |  |  |  | 0.048 (0.032),  -0.012 – 0.106 | 0.046 (0.028),  0.007 – 0.123 |
| γ_NEUTRAL_ |  |  |  |  |  |  | 0.028 (0.026),  -0.022 – 0.093 | 0.028 (0.022),  -0.016 – 0.082 |
| γ_ANGRY_ |  |  |  |  |  |  | 0.008 (0.030),  -0.044 – 0.102 | 0.005 (0.027),  -0.036 – 0.059 |
| Mean (standard deviation), minimum – maximum parameter values in computational reinforcement learning models. α = learning rate, β = inverse temperature, γ = approach bias (per emotion). Parameter were fitted with hierarchical Bayesian inference using the CBM toolbox in MATLAB. Values are fitted and reported separately for mIns and sham sessions. | | | | | | | | |

**Table S25:** Parameter estimates obtained from models M1-M4 for the in-person TUS cohort, comparing sham and mIns sonication conditions (n = 29).

#### Overview tables regression results MRS

##### Overview tables MRS results BLA voxel, n = 11

| Model ID | Effect | *b* | 95% CI | | *df* | χ^2^ | *p* |
| --- | --- | --- | --- | --- | --- | --- | --- |
|  | *DV: E/I balance* |  |  |  |  |  |  |
| 1 | Intercept | -0.155 | -0.504 | 0.194 | 1 | 0.760 | .383 |
|  | BLA TUS vs. sham | -0.233 | -0.500 | 0.034 | 1 | 2.924 | .087 |
| 2 | Intercept | -0.244 | -0.615 | 0.167 | 1 | 1.256 | .262 |
|  | BLA TUS vs. sham | -0.233 | -0.471 | 0.005 | 1 | 3.676 | .055 |
|  | Gender (female vs. male) | 0.024 | -0.439 | 0.487 | 1 | 0.010 | .919 |
|  | Age | -0.565 | -1.068 | -0.062 | 1 | 4.849 | .028 |
|  | Time of day (morning vs. afternoon) | 0.243 | -0.150 | 0.635 | 1 | 1.470 | .225 |
|  | Overlap simulated beam & MRS voxel | -0.203 | -0.748 | 0.341 | 1 | 0.536 | .464 |
|  | *DV: GABA concentration* |  |  |  |  |  |  |
| 3 | Intercept | 0.078 | -0.243 | 0.400 | 1 | 0.226 | .634 |
|  | BLA TUS vs. sham | 0.369 | 0.047 | 0.690 | 1 | 5.057 | .025 |
| 4 | Intercept | 0.122 | -0.228 | 0.472 | 1 | 0.463 | .496 |
|  | BLA TUS vs. sham | 0.369 | 0.043 | 0.695 | 1 | 4.921 | .027 |
|  | Gender (male vs. female) | -0.145 | -0.557 | 0.267 | 1 | 0.474 | .491 |
|  | Age | 0.299 | -0.164 | 0.762 | 1 | 1.606 | .205 |
|  | Time of day (morning vs. afternoon) | -0.111 | -0.521 | 0.298 | 1 | 0.284 | .594 |
|  | Overlap simulated beam & MRS voxel | -0.001 | -0.490 | 0.488 | 1 | 0 | .998 |
|  | *DV: Glutamate concentration* |  |  |  |  |  |  |
| 5 | Intercept | 0.020 | -0.555 | 0.595 | 1 | 0.005 | .946 |
|  | BLA TUS vs. sham | 0.149 | -0.055 | 0.353 | 1 | 2.054 | .152 |
| 6 | Intercept | 0.085 | -0.375 | -0.546 | 1 | 0.133 | .716 |
|  | BLA TUS vs. sham | 0.149 | -0.057 | 0.355 | 1 | 2.012 | .156 |
|  | Gender (male vs. female) | -0.416 | -0.964 | 0.131 | 1 | 2.220 | .136 |
|  | Age | -0.605 | -1.183 | -0.027 | 1 | 4.206 | .040 |
|  | Time of day (morning vs. afternoon) | -0.101 | -0.490 | 0.288 | 1 | 0.259 | .611 |
|  | Overlap simulated beam & MRS voxel | -0.179 | -0.819 | 0.461 | 1 | 0.301 | .583 |
| *Overview results from mixed-effects linear regression models*. Mixed-effects linear regression models were fit with the lmer() function from lme4 package in R. All continuous variables (E/I balance, GABA, glutamate, age, overlap) were z-standardized. All factors were coded with sum-to-zero coding (BLA = 1, sham = -1; female = 1, male = -1; morning = 1, afternoon = -1) such that regression coefficients can be interpreted as standardized regression coefficients. | | | | | | | |

**Table S26:** MRS regression comparing BLA-TUS and sham in n = 10 participants.

##### Overview tables MRS results mIns voxel, n = 18

| Model ID | Effect | *b* | 95% CI | | *df* | χ^2^ | *p* |
| --- | --- | --- | --- | --- | --- | --- | --- |
|  | *DV: E/I balance* |  |  |  |  |  |  |
| 1 | Intercept | -0.112 | -0.488 | 0.265 | 1 | 0.340 | .560 |
|  | mIns TUS vs. sham | 0.321 | 0.079 | 0.563 | 1 | 6.757 | .009 |
| 2 | Intercept | -0.133 | -0.570 | 0.305 | 1 | 0.352 | .553 |
|  | mIns TUS vs. sham | 0.309 | 0.060 | 0.557 | 1 | 5.905 | .015 |
|  | Gender (female vs. male) | -0.269 | -0.711 | 0.173 | 1 | 1.425 | .233 |
|  | Age | 0.207 | -0.231 | 0.644 | 1 | 0.857 | .354 |
|  | Time of day (morning vs. afternoon) | -0.114 | -0.576 | 0.349 | 1 | 0.232 | .630 |
|  | Overlap simulated beam & MRS voxel | 0.188 | -0.233 | 0.609 | 1 | 0.763 | .382 |
|  | *DV: GABA concentration* |  |  |  |  |  |  |
| 3 | Intercept | 0.102 | -0.253 | 0.457 | 1 | 0.315 | .547 |
|  | mIns TUS vs. sham | -0.347 | -0.585 | -0.110 | 1 | 8.207 | .004 |
| 4 | Intercept | 0.047 | -0.386 | 0.479 | 1 | 0.045 | .833 |
|  | mIns TUS vs. sham | -0.359 | -0.603 | -0.115 | 1 | 8.318 | .004 |
|  | Gender (male vs. female) | 0.069 | -0.368 | 0.506 | 1 | 0.097 | .756 |
|  | Age | -0.052 | -0.485 | 0.381 | 1 | 0.056 | .813 |
|  | Time of day (morning vs. afternoon) | -0.107 | -0.562 | 0.349 | 1 | 0.211 | .646 |
|  | Overlap simulated beam & MRS voxel | -0.132 | -0.549 | 0.284 | 1 | 0.387 | .534 |
|  | *DV: Glutamate concentration* |  |  |  |  |  |  |
| 5 | Intercept | -0.063 | -0.527 | 0.402 | 1 | 0.070 | .792 |
|  | mIns TUS vs. sham | 0.092 | -0.127 | 0.311 | 1 | 0.678 | .410 |
| 6 | Intercept | -0.422 | -0.953 | 0.109 | 1 | 2.425 | .119 |
|  | mIns TUS vs. sham | -0.013 | -0.162 | 0.136 | 1 | 0.030 | .864 |
|  | Gender (male vs. female) | -0.549 | -1.083 | -0.014 | 1 | 4.042 | .044 |
|  | Age | 0.537 | -0.011 | 1.085 | 1 | 3.689 | .055 |
|  | Time of day (morning vs. afternoon) | -0.946 | -1.332 | -0.559 | 1 | 23.013 | < .001 |
|  | Overlap simulated beam & MRS voxel | 0.057 | -0.482 | 0.596 | 1 | 0.043 | .837 |
| *Overview results from mixed-effects linear regression models*. Mixed-effects linear regression models were fit with the lmer() function from lme4 package in R. All continuous variables (E/I balance, GABA, glutamate, age, overlap) were z-standardized. All factors were coded with sum-to-zero coding (mIns = 1, sham = -1; female = 1, male = -1; morning = 1, afternoon = -1) such that regression coefficients can be interpreted as standardized regression coefficients. | | | | | | | |

**Table S27:** MRS regression comparing mIns-TUS and sham in n = 8 participants.

##### Overview tables MRS results contrasting both voxels, n = 7

| Model ID | Effect | *b* | 95% CI | | *df* | χ^2^ | *p* |
| --- | --- | --- | --- | --- | --- | --- | --- |
|  | *DV: E/I balance* |  |  |  |  |  |  |
| 1 | Intercept | .001 | -0.462 | 0.462 | 1 | 0 | 1 |
|  | BLA vs. mIns voxel | -0.705 | -0.956 | -0.454 | 1 | 30.317 | < .001 |
| 2 | Intercept | -0.006 | -0.205 | 0.192 | 1 | 0.004 | .950 |
|  | BLA vs. mIns voxel | -0.759 | -0.952 | -0.565 | 1 | 59.182 | < .001 |
|  | Gender (female vs. male) | -0.045 | -0.458 | 0.368 | 1 | 0.045 | .831 |
|  | Age | 0.265 | -0.116 | 0.646 | 1 | 1.856 | .173 |
|  | Time of day (morning vs. afternoon) | 0.374 | 0.123 | 0.625 | 1 | 8.544 | .003 |
|  | Overlap BLA sim. beam & MRS voxel | 0.004 | -0.511 | 0.518 | 1 | 0.000 | .989 |
|  | Overlap mIns sim. beam & MRS voxel | 0.707 | 0.325 | 1.089 | 1 | 13.136 | < .001 |
|  | *DV: GABA concentration* |  |  |  |  |  |  |
| 3 | Intercept | 0.000 | -0.469 | 0.469 | 1 | 0.000 | 1 |
|  | BLA vs. mIns voxel | 0.631 | 0.286 | 0.977 | 1 | 12.842 | < .001 |
| 4 | Intercept | -0.016 | -0.370 | 0.337 | 1 | 0.008 | .929 |
|  | BLA vs. mIns voxel | 0.670 | 0.332 | 1.009 | 1 | 15.111 | < .001 |
|  | Gender (female vs. male) | -0.113 | -0.846 | 0.621 | 1 | 0.090 | .764 |
|  | Age | -0.273 | -0.951 | 0.404 | 1 | 0.626 | .429 |
|  | Time of day (morning vs. afternoon) | -0.273 | -0.718 | 0.171 | 1 | 1.449 | .229 |
|  | Overlap BLA sim. beam & MRS voxel | -0.064 | -0.978 | 0.849 | 1 | 0.019 | .890 |
|  | Overlap mIns sim. beam & MRS voxel | -0.689 | -1.368 | -0.009 | 1 | 3.948 | .047 |
|  | *DV: Glutamate concentration* |  |  |  |  |  |  |
| 5 | Intercept | 0.000 | -0.630 | 0.630 | 1 | 0.000 | 1 |
|  | BLA vs. mIns voxel | 0.080 | -0.360 | 0.521 | 1 | 0.128 | .720 |
| 6 | Intercept | 0.074 | -0.872 | 1.019 | 1 | 0.023 | .879 |
|  | BLA vs. mIns voxel | 0.013 | -0.412 | 0.438 | 1 | 0.003 | .953 |
|  | Gender (female vs. male) | 0.515 | -1.341 | 2.372 | 1 | 0.296 | .586 |
|  | Age | -0.261 | -2.056 | 1.535 | 1 | 0.081 | .776 |
|  | Time of day (morning vs. afternoon) | 0.474 | -0.406 | 1.355 | 1 | 1.113 | .291 |
|  | Overlap BLA sim. beam & MRS voxel | -1.320 | -3.545 | 0.906 | 1 | 1.350 | .245 |
|  | Overlap mIns sim. beam & MRS voxel | 0.009 | -1.796 | 1.814 | 1 | 0 | .992 |
| *Overview results from mixed-effects linear regression models*. Mixed-effects linear regression models were fit with the lmer() function from lme4 package in R. All continuous variables (E/I balance, GABA, glutamate, age, overlap) were z-standardized. All factors were coded with sum-to-zero coding (BLA = 1, mIns = -1; female = 1, male = -1; morning = 1, afternoon = -1) such that regression coefficients can be interpreted as standardized regression coefficients. | | | | | | | |

**Table S28:** MRS regression comparing mIns-TUS, BLA-TUS and sham in n = 5 participants.

#### Overview of TUS side effects

| **Symptom** | **c2** | **p** | **FDR corrected p** |
| --- | --- | --- | --- |
| Burning Heat | 11.62 | 0.003 | 0.0777 |
| Tingling | 9.48 | 0.0087 | 0.1136 |
| Hearing | 4 | 0.1353 | 0.7037 |
| Scalp Sensations | 4.19 | 0.1232 | 0.7037 |
| Twitches | 4.31 | 0.116 | 0.7037 |
| Balance Problems | 2 | 0.3679 | 0.8695 |
| Cold Sensation | 2.33 | 0.3114 | 0.8695 |
| Happy | 2 | 0.3679 | 0.8695 |
| Headache | 2.15 | 0.3406 | 0.8695 |
| Itchiness | 2.25 | 0.3247 | 0.8695 |
| Vision | 2 | 0.3679 | 0.8695 |
| Forgetfulness | 1.37 | 0.5045 | 1.0931 |
| Attention | 1.19 | 0.5512 | 1.1023 |
| Other Emotions | 1 | 0.6065 | 1.1264 |
| Dizziness | 0.73 | 0.6951 | 1.1296 |
| Nerve Pain | 0.74 | 0.6918 | 1.1296 |
| Nausea | 0.5 | 0.7788 | 1.1911 |
| Anxiety | 0.15 | 0.926 | 1.2037 |
| Sleepiness | 0.27 | 0.8717 | 1.2037 |
| Stress | 0.2 | 0.9048 | 1.2037 |
| Breathing Problems | NaN | NaN | NaN |
| Muscle Pain | NaN | NaN | NaN |
| Neck Pain | NaN | NaN | NaN |
| Other | NaN | NaN | NaN |
| Speech Problems | NaN | NaN | NaN |
| Tooth Pain | NaN | NaN | NaN |

**Table S29:** Results for the non-parametric repeated measures Friedman test with false discovery rate correction for 26 comparisons conducted on symptom severity scores after the three TUS sessions (BLA TUS, mIns TUS and sham). Here we included only the severity scores rated as at least mild (severity > 1) and with a reported relation to TUS as at least possible (relation > 2). We note that while no side effects were significantly different between the two active sessions, mIns/BLA differed from sham on two sensory scalp sensation measures: burning heat and tingling.

| **Symptom** | **BLA TUS** | **mIns TUS** | **Sham** |
| --- | --- | --- | --- |
| Burning Heat | 13 | 9 | 3 |
| Tingling | 13 | 6 | 4 |
| Scalp Sensations | 12 | 9 | 5 |
| Attention | 10 | 8 | 7 |
| Sleepiness | 9 | 7 | 7 |
| Itchiness | 5 | 2 | 2 |
| Dizziness | 4 | 5 | 6 |
| Cold Sensation | 3 | 4 | 3 |
| Forgetfulness | 3 | 5 | 2 |
| Headache | 3 | 3 | 6 |
| Nerve Pain | 3 | 2 | 2 |
| Vision | 3 | 4 | 2 |
| Balance Problems | 2 | 4 | 2 |
| Nausea | 2 | 1 | 1 |
| Twitches | 2 | 5 | 5 |
| Anxiety | 1 | 2 | 2 |
| Happy | 1 | 4 | 1 |
| Muscle Pain | 1 | 0 | 0 |
| Other Emotions | 1 | 1 | 0 |
| Stress | 1 | 2 | 1 |
| Breathing Problems | 0 | 0 | 1 |
| Hearing | 0 | 0 | 2 |
| Neck Pain | 0 | 1 | 0 |
| Other | 0 | 1 | 0 |
| Speech Problems | 0 | 0 | 0 |
| Tooth Pain | 0 | 0 | 1 |

**Table S30:** Frequency of occurrence of each symptom following TUS at times when the relation to the stimulation was reported as more likely than “possible”. Among the symptoms more frequently experienced during BLA TUS as at least mild and perceived as possibly related to TUS are burning/heat sensations (n = 13), tingling (n = 13), and scalp sensations (n = 12). These were reported in the insula sessions with similar frequency: burning/heat sensations (n = 9 for insula TUS, n = 3 for sham), tingling (n = 6 for insula TUS, n = 4 for sham), scalp sensations (n = 9 for insula TUS, n = 5 for sham).
